# Reconstructing the spatiotemporal patterns of admixture during the European Holocene using a novel genomic dating method

**DOI:** 10.1101/2022.01.18.476710

**Authors:** Manjusha Chintalapati, Nick Patterson, Priya Moorjani

**Author notes:** To whom correspondence should be addressed: Manjusha Chintalapati < >, Nick Patterson < > and Priya Moorjani < >.

## Abstract

Recent studies have shown that gene flow or admixture has been pervasive throughout human history. While several methods exist for dating admixture in contemporary populations, they are not suitable for sparse, low coverage data available from ancient specimens. To overcome this limitation, we developed *DATES* that leverages ancestry covariance patterns across the genome of a single individual to infer the timing of admixture. By performing simulations, we show that *DATES* provides reliable results under a range of demographic scenarios and outperforms available methods for ancient DNA applications. We apply *DATES* to ~1,100 ancient genomes to reconstruct gene flow events during the European Holocene. Present-day Europeans derive ancestry from three distinct groups, local Mesolithic hunter-gatherers, Anatolian farmers, and Yamnaya Steppe pastoralists. These ancestral groups were themselves admixed. By studying the formation of Anatolian farmers, we infer that the gene flow related to Iranian Neolithic farmers occurred no later than 9,600 BCE, predating agriculture in Anatolia. We estimate the early Steppe pastoralist groups genetically formed more than a millennium before the start of steppe pastoralism, providing new insights about the history of proto-Yamnaya cultures and the origin of Indo-European languages. Using ancient genomes across sixteen regions in Europe, we provide a detailed chronology of the Neolithization across Europe that occurred from ~6,400–4,300 BCE. This movement was followed by a rapid spread of steppe ancestry from ~3,200–2,500 BCE. Our analyses highlight the power of genomic dating methods to elucidate the legacy of human migrations, providing insights complementary to archaeological and linguistic evidence.

**Significance:** The European continent was subject to two major migrations during the Holocene: the movement of Near Eastern farmers during the Neolithic and the migration of Steppe pastoralists during the Bronze Age. To understand the timing and dynamics of these movements, we developed *DATES* that leverages ancestry covariance patterns across the genome of a single individual to infer the timing of admixture. Using ~1,100 ancient genomes spanning ~8,000–350 BCE, we reconstruct the chronology of the formation of the ancestral populations and the fine-scale details of the spread of Neolithic farming and Steppe pastoralist-related ancestry to Europe. Our analysis demonstrates the power of genomic dating methods to provide an independent and complementary timeline of population origins and movements using genetic data.

## Introduction

Recent studies have shown that population mixture (or “admixture”) is pervasive throughout human history, including mixture between the ancestors of modern humans and archaic hominins (i.e., Neanderthals and Denisovans), as well as in the history of many contemporary human groups such as African Americans, South Asians and Europeans (1, 2). Many admixed groups are formed due to population movements involving ancient migrations that pre-date historical records. The recent availability of genomic data for a large number of present-day and ancient genomes provides an unprecedented opportunity to reconstruct population events using genetic data, providing evidence complementary to linguistics and archaeology. Understanding the timing and signatures of admixture offers insights into the historical context in which the mixture occurred and enables the characterization of the evolutionary and functional impact of the gene flow.

To characterize patterns of admixture, genetic methods use the insight that the genome of an admixed individual is a mosaic of chromosomal segments inherited from distinct ancestral populations (3). Due to recombination, these ancestral segments get shuffled in each generation and become smaller and smaller over time. The length of the segments is inversely proportional to the time elapsed since the mixture (3, 4). Several genetic approaches—ROLLOFF (4), ALDER (5), Globetrotter (2), and Tracts (6)— have been developed that use this insight by characterizing patterns of admixture linkage disequilibrium (LD) or haplotype lengths across the genome to infer the timing of mixture. Haplotype-based methods perform chromosome painting or local ancestry inference at each locus in the genome and characterize the distribution of ancestry tract lengths to estimate the time of mixture (2, 6). This requires accurate phasing and inference of local ancestry, which is often difficult when the admixture events are old (as ancestry blocks become smaller over time) or when reference data from ancestral populations is unavailable. Admixture LD-based methods, on the other hand, measure the extent of the allelic correlation across markers to infer the time of admixture (4, 5). They do not require phased data from the target or reference populations and work reliably for dating older admixture events (>100 generations). However, they tend to be less efficient in characterizing admixture events between closely related ancestral groups.

While highly accurate for dating admixture events using data from present-day samples, current methods do not work reliably for dating admixture events using ancient genomes. Ancient DNA samples often have high rates of DNA degradation, contamination (from human and other sources) and low sequencing depth, leading to a large proportion of missing variants and uneven coverage across the genome. Additionally, most studies generate pseudo-homozygous genotype calls—consisting of a single allele call at each diploid site—that can lead to some issues in the inference. In such sparse datasets, estimating admixture LD can be noisy and biased (see Simulations below). Moreover, haplotype-based methods require phased data from both admixed and reference populations which remains challenging for ancient DNA specimens.

An extension of admixture LD-based methods, recently introduced by Moorjani et al. (2016), leverages ancestry covariance patterns that can be measured in a single sample using low coverage data. This approach measures the allelic correlation across neighboring sites, but instead of measuring admixture LD across multiple samples, it integrates data across markers within a single diploid genome. Using a set of ascertained markers that are informative for Neanderthal ancestry (where sub-Saharan Africans are fixed for the ancestral alleles and Neanderthals have a derived allele), Moorjani et al. (2016) inferred the timing of Neanderthal gene flow in Upper Paleolithic Eurasian samples and showed the approach works accurately in ancient DNA samples (1). However, this approach is inapplicable for dating admixture events within modern human populations, as there are very few fixed differences across populations (7).

Motivated by the single sample statistic in Moorjani et al. (2016), we developed *DATES* (*Distribution of Ancestry Tracts of Evolutionary Signals*) that measures the ancestry covariance across the genome in a single admixed individual, weighted by the allele frequency difference between two ancestral populations. This method was first introduced in Narasimhan et al. (2019), where it was used to infer the date of gene flow between groups related to Ancient Ancestral South Indians, Iranian farmers, and Steppe pastoralists in ancient South and Central Asian populations (8). In this study, we evaluate the performance of *DATES* by performing extensive simulations for a range of demographic scenarios and compare the approach to other published genomic dating methods. We then apply *DATES* to infer the chronology of the genetic formation of the ancestral populations of Europeans and the spatiotemporal patterns of admixture during the European Holocene using data from ~1,100 ancient DNA specimens spanning ~8,000–350 BCE.

## Results

### Overview of *DATES*: Model and simulations

*DATES* estimates the time of admixture by measuring the weighted ancestry covariance across the genome using data from a single diploid genome and two reference populations (representing the ancestral source populations). *DATES* works like haplotype-based methods as it is applicable to dating admixture in a single genome and not like admixture LD-based methods, which by definition require multiple genomes to be co-analyzed; but unlike haplotype-based methods, it is more flexible as it does not require local ancestry inference. There are three main steps in *DATES*: we start by first learning the genome-wide ancestry proportions by performing a simple regression analysis to model the observed genotypes in an admixed individual as a linear mix of allele frequencies from the two reference populations. For each marker, we then compute the likelihood of the observed genotype in the admixed individual using the estimated ancestry proportions and allele frequencies in each reference population (this is similar in spirit to local ancestry inference). This information is, in turn, used to compute the joint likelihood for two neighboring markers to test if they derive ancestry from the same ancestral group, accounting for the probability of recombination between the two markers. Finally, we compute the covariance across pairs of markers located at a particular genetic distance, weighted by the allele frequency differences in the reference populations (Note S1).

Following (1), we bin the markers that occur at a similar genetic distance across the genome, rather than estimating admixture LD for each pair of markers, and compute the covariance across increasing genetic distance between markers. The estimated covariance is expected to decay exponentially with genetic distance, and the rate of decay is informative of the time of the mixture (4). Assuming the gene flow occurred instantaneously, we infer the average date of gene flow by fitting an exponential distribution to the decay pattern (Methods). In cases where data for multiple individuals is available, we compute the likelihood by summing over all individuals. To make *DATES* computationally tractable, we implemented the fast Fourier transform (FFT) for calculating ancestry covariance as described in ALDER (5). This provides a speedup from *O*(*n*^2^) to *O*(*n* log *n*), which reduces the typical runtimes from hours to seconds (Note S1).

To assess the reliability of *DATES*, we performed simulations where we constructed ten admixed diploid genomes by randomly sampling haplotypes from two source populations (Note S2). Briefly, we simulated individual genomes with 20% European and 80% African ancestry by using phased haplotypes of Northern Europeans (Utah European Americans, CEU) and west Africans (Yoruba from Nigeria, YRI) from the 1000 Genomes Project respectively (7). As reference populations in *DATES*, we used closely related surrogate populations of French and Yoruba respectively, from the Human Genome Diversity Panel (9). We first investigated the accuracy of *DATES* by varying the time of admixture between 10–300 generations. For comparison, we also applied ALDER (5) to these simulations. Both methods reliably recovered the time of admixture up to 200 generations or ~5,600 years ago, assuming a generation time of 28 years (1), though *DATES* was more precise than ALDER for older admixture events (>100 generations) (Table S2.4). Further, *DATES* shows accurate results even for single samples (Figure 1).

**Figure 1:**
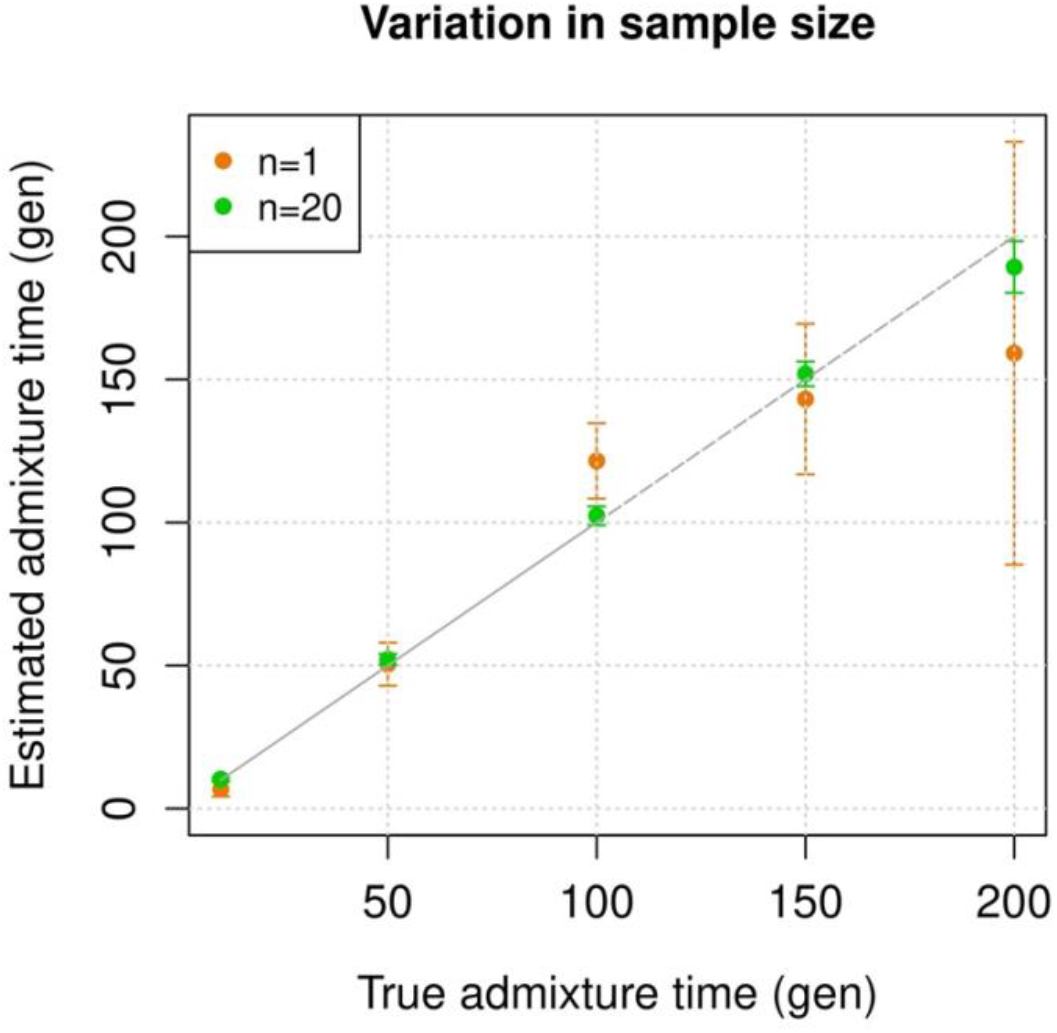
Simulation results. We constructed admixed individuals with 20% European (CEU) and 80% Africa (YRI) ancestry for admixture dates ranging between 10–200 generations where the sample size of the target group is as shown in the legend. We applied *DATES* using French and Yoruba as reference populations. We show the true time of admixture (X-axis, in generations) and the estimated time of admixture (± 1 SE) (Y-axis, in generations). Standard errors were calculated using a weighted block jackknife approach by removing one chromosome in each run (Methods).

Next, we tested *DATES* for features such as varying admixture proportions and use of surrogate populations as reference groups. By varying of European ancestry proportion between ~1–50% (the rest derived from west Africans), we observed *DATES* accurately estimated the timing in all cases (Figure S2.2A). However, the inferred admixture proportion was overestimated was lower admixture proportions (<10%) (Figure S2.2B). Thus, we caution against using *DATES* for estimating ancestry proportions and recommend other methods based on *f*-statistics (10). Using reference populations which are divergent from true admixing source, we found that the inferred dates were accurate even when we used Khomani San instead of Yoruba as the reference population (*F_ST_* ~ 0.1) (Figure S2.5). We also found that using the admixed samples themselves as one of the reference populations also works reliably as ALDER (i.e., single reference setup) (5).

An important feature of *DATES* is that it does not require phased data and is applicable to datasets with small sample sizes, making it in principle useful for ancient DNA applications. To test the reliability of *DATES* for ancient genomes, we simulated data mimicking the relevant features of ancient genomes, namely with large proportions of missing genotypes (between 10–60%), and pseudo-homozygous genotype calls (instead of diploid genotype calls). *DATES* showed reliable results in both cases, even only a single admixed individual was available (Figure S2.7). In contrast, admixture LD-based methods require more than one sample and do not work reliably with missing data. For example, ALDER estimates were very unstable for simulations with >40% missing data. For older dates (>100 generations), we observed slight bias even with >10% missing genotypes (Figure S2.17). As LD calculations leverage shared patterns across samples, variable missingness of genotypes across individuals leads to substantial loss of data leading to unstable and noisy inference. This highlights a major advantage of *DATES* for ancient DNA studies as it provides reliable results even in sparse datasets (Note S2.5).

*DATES* assumes a model of instantaneous gene flow with a single pulse of mixture between two source populations. However, many human populations have a history of multiple pulses of gene flow. To test the performance of *DATES* for multi-way admixture events, we generated admixed individuals with ancestry from three sources (East Asians, Africans, and Europeans) where the gene flow occurred at two distinct time points (Note S2, Figure S2.10). By applying *DATES* with pairs of reference populations at a time and fitting a single exponential to the ancestry covariance patterns, we observed that *DATES* recovered both admixture times in case of equal ancestry proportion from the three ancestral groups when the associated reference groups were used for dating (Figure S2.11). In the case of unequal admixture proportions from three ancestral groups, *DATES* inferred the timing of the recent admixture event in most cases, though some confounding was observed, especially when the ancestry proportion of the recent event was low (Figure S2.12). However, if the reference populations were set up to match the model of gene flow, we observed that we could reliably recover the time of the recent gene flow event. For example, there is limited confounding if the two references used in *DATES* include (i) the source population for the recent event and (ii) either the pooled ancestral populations contributing to the first (or earlier) event or the intermediate admixed group formed after the first event (Table S2.1). This highlights how the choice of reference populations can help to tune the method to infer the timing of specific admixture events reliably.

Finally, we explored the impact of more complex demographic events, including continuous admixture and founder events using coalescent simulations (Note S2). In the case of continuous admixture, *DATES* inferred an intermediate timing between the start and the end of the gene flow period, similar to other methods like ALDER and Globetrotter (2, 5) (Table S2.2). In the case of populations with founder events, we inferred unbiased dates of admixture in most cases except when the founder event was extreme (*N_e_* ~ 10) or the population had maintained a low population size (*N_e_* < 100) until present (i.e., no recovery bottleneck) (Figure S2.13, Table S2.3). In humans, few populations have such extreme founder events, and thus, in most other cases, our inferred admixture dates should be robust to founder events (11). We note that while *DATES* is not a formal test of admixture, in simulations, we find that in the absence of gene flow, the method does not infer significant dates of admixture even when the target has a complex demographic history (Figure S2.15, S2.16).

### Comparison to other methods

We assessed the reliability of *DATES* in real data by comparing our results with published methods: Globetrotter, ALDER, and ROLLOFF. These methods are designed for the analysis of present-day samples that typically have high-quality data with limited missing variants. In addition, Globetrotter uses phased data which is challenging for ancient DNA samples. Thus, instead of rerunning other methods, we took advantage of the published results for contemporary samples presented in Hellenthal et al., (2014) (2). Following (2), we created a merged dataset including individuals from Human Genome Diversity Panel (9), Behar et al. (2010) (12), and Henn et al. (2012) (13) (Methods). We applied *DATES* and ALDER to 29 target groups using the reference populations reported in Hellenthal et al. 2014 (Table S12), excluding one group where the population label was unclear. Interestingly, the majority of these groups (25/29) failed ALDER’s formal test of admixture; either because the results of the single reference and two reference analyses yielded inconsistent estimates or because the target had long-range shared LD with one of the reference populations (Table S4.1). Using *DATES*, we inferred significant dates of admixture in 20 groups, and 14 of those were consistent with estimates based on Globetrotter. In most remaining cases, recent studies suggest the target populations may have ancestry from multiple gene flow events, either involving the same source populations or additional ancestral groups. The estimated admixture timing based on *DATES*, ROLLOFF, and ALDER (assuming two-way admixture regardless of the formal test results) were found to be highly concordant (Table S4.1).

### Fine-scale patterns of population mixtures in ancient Europe

Recent ancient DNA studies have shown that present-day Europeans derive ancestry from three distinct sources: (a) hunter-gatherer-related ancestry that is closely related to Mesolithic hunter-gatherers (HG) from Europe; (b) Anatolian farmer-related ancestry related to Neolithic farmers from the Near East and associated to the spread of farming to Europe; and (c) Steppe pastoralist-related ancestry that is related to the Yamnaya pastoralists from Russia and Ukraine (16–19). Many open questions remain about the timing and dynamics of these population interactions, in particular related to the formation of the ancestral groups (which were themselves admixed) and their expansion across Europe. To characterize the spatial and temporal patterns of mixtures in Europe in the past 10,000 years, we used 1,096 ancient European samples from 152 groups from the publicly available Allen Ancient DNA Resource (AADR) spanning a time range of ~8,000–350 BCE (Methods, Table SA). Using *DATES*, we characterized the timing of the various gene flow events, and below, we describe the key events in chronological order focusing on three main periods.

#### Holocene to Mesolithic

Pre-Neolithic Europe was inhabited by hunter-gatherers until the arrival of the first farmers from the Near East (20, 21). There was large diversity among hunter-gatherers with four main groups— western hunter-gatherers (WHG) that were related to the Villabruna cluster in central Europe, eastern hunter-gatherers (EHG) from Russia and Ukraine related to the Upper Paleolithic group of Ancestral North Eurasians (ANE) ancestry, Caucasus hunter-gatherers (CHG) from Georgia associated to the first farmers from Iran, and the GoyetQ2-cluster associated to the Magdalenian culture in Spain and Portugal (18, 22–25). Most Mesolithic HGs fall on two main clines of relatedness: one cline that extends from Scandinavia to central Europe showing variable WHG–EHG ancestry, and the other in southern Europe with WHG–GoyetQ2 ancestry (23). This ancestry is already present in the 17,000 BCE *El Mirón* individual from Spain, suggesting that the GoyetQ2-related gene flow occurred well before the Holocene. However, the WHG-EHG cline was formed more recently during the Mesolithic period, though the precise timing of the spread of EHG ancestry remains less well understood.

To characterize the formation of the WHG–EHG cline, we used genomic data from 16 ancient HG groups (*n*=101) with estimated ages of ~7,500–3,600 BCE. We first verified the ancestry of each HG group using *qpAdm* that compares the allele frequency correlations between the target and a set of source populations to formally test the model of admixture and then infer the ancestry proportions for the best fitted model (16). For each target population, we chose the most parsimonious model, i.e., fitting the data with the minimum number of source populations. Consistent with previous studies, our *qpAdm* analysis showed that most HGs from Scandinavia, the Baltic Sea region, and central Europe could be modeled as a two-way mixture of WHG and EHG-related ancestry (Table S5.1, Note S5). To confirm that the target populations do not harbor Anatolian farmer-related ancestry (that could lead to some confounding in estimated admixture dates), we applied *D*-statistics of the form *D*(Mbuti, *target*, WHG, Anatolian farmers) where *target* = Mesolithic HGs. We observed that none of the target groups had a stronger affinity to Anatolian farmers than WHG, suggesting that the mixtures we date below reflect pre-Neolithic contacts between the HGs (Table S5.2).

To infer the timing of the mixtures in the history of Mesolithic European HGs, we applied *DATES* to hunter-gatherers from Scandinavia, the Baltic regions, and central Europe. We inferred that the earliest admixture occurred in Scandinavian HGs from Norway and Sweden around ~80–113 generations before the samples lived (Figure SB). Accounting for the average sampling age of the specimens and the mean human generation time of 28 years (1), this translates to a timing of admixture of ~10,200 BCE for Norway and Sweden Mesolithic individuals, though dates are more recent (~8,000 BCE) in the Motala HG’s. In the Baltic region, we inferred admixture dates of ~8,700–6,000 BCE in Latvia and Lithuania HGs, postdating the mixture in Scandinavia (Figure 3). In southeast Europe, the Iron Gates region of the Danube Basin shows widespread evidence of mixtures between hunter-gatherer groups and, in the case of some outliers, mixture of hunter-gatherers and Anatolian farmer-related ancestry as early as the Mesolithic period (26). Further, these groups showed strong affinity to the WHG-related ancestry in Anatolian populations, suggesting ancient interactions with Near Eastern populations (26). We applied *qpAdm* to test the model of admixture in Iron Gates HG and found that the parsimonious model with WHG and EHG provides a good fit to the data. Further, when we tested the model with Anatolian-related ancestry using Anatolian HG (AHG) as an additional source population, AHG was not required as the AHG ancestry proportion was not significant (Table S5.1.1 and S5.1.2). Applying *DATES* to Iron Gates HG with WHG and EHG as reference populations, we inferred this group was genetically formed in ~10,000–8400 BCE. Our samples of the Iron Gates HGs include a wide range of C14 dates between 8,800–5,700 BCE. We confirmed our dates were robust to the sampling age of the individuals as we obtained statistically consistent dates when all samples were combined as one group or when subsets of samples were grouped in bins of 500 years (Figure SA). The most recent dates of ~7,500 BCE were inferred in eastern Europe in Ukraine HGs, highlighting how the WHG-EHG cline was formed over a period ~2000–3000 years (Figure 3, Table SC).

**Figure 2:**
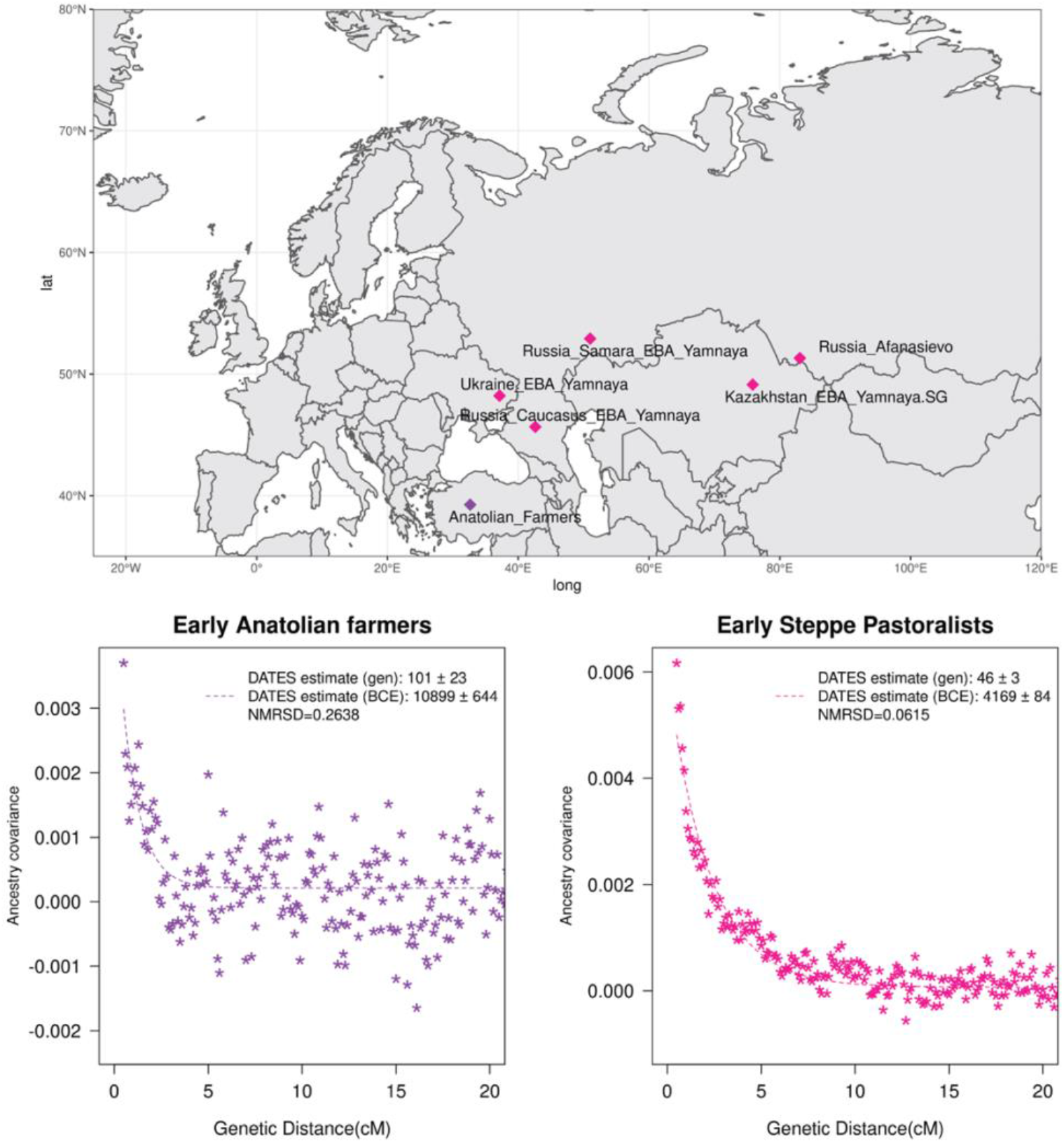
Genetic formation of early Anatolian farmers and early Bronze Age Steppe pastoralists. The top panel shows a map with sampling locations of the target groups analyzed for admixture dating. The bottom panels show the inferred times of admixture for each target using *DATES* by fitting an exponential function with an affine term *y* = *Ae^−λd^* + *c*, where *d* is the genetic distance in Morgans and *λ* = (*t*+1) is the number of generations since admixture (*t*) (Methods). We start the fit at a genetic distance (*d*) > 0.5cM to minimize confounding with background LD and estimate a standard error by performing a weighted block jackknife removing one chromosome in each run. For each target, in the legend, we show the inferred average dates of admixture (± 1 SE) in generations before the individual lived, in BCE accounting for the average age of all the individuals and the mean human generation time, and the NRMSD values to assess the fit of the exponential curve (Methods). The bottom left shows the ancestry covariance decay curve for early Anatolian farmers inferred using one reference group as a set of pooled individuals of WHG-related and Levant Neolithic farmers-related individuals as a proxy of AHG ancestry and the second reference group containing Iran Neolithic farmer-related individuals. The bottom right shows the ancestry covariance decay curve for early Steppe pastoralists groups, including all Yamnaya and Afanasievo individuals as the target group and EHG-related and Iran Neolithic farmer-related groups as reference populations.

**Figure 3:**
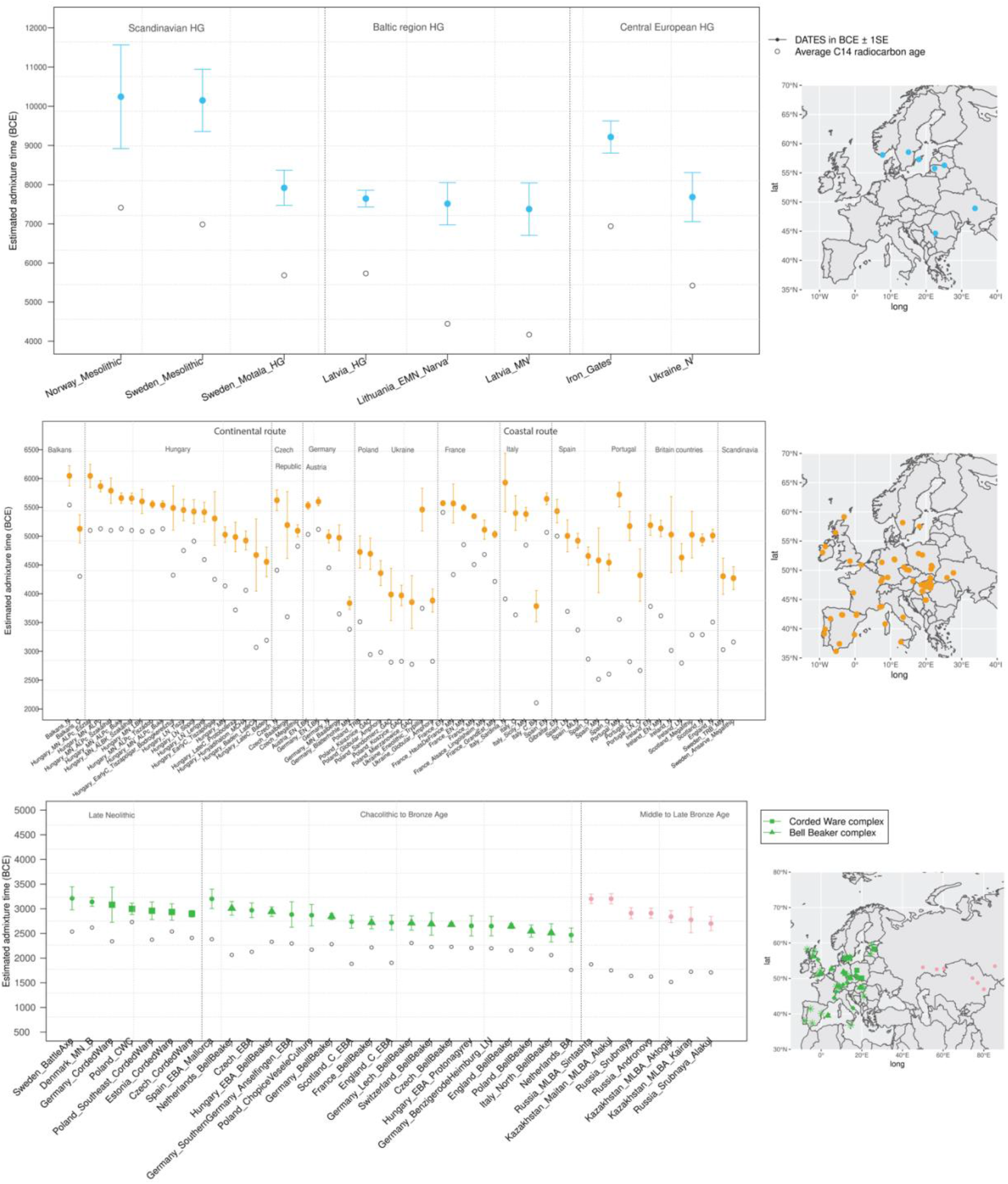
Timeline of admixture events in ancient Europe. We applied *DATES* to ancient samples from Europe. In the right panel, we show the sampling locations of the ancient specimens, and in the left panel, we show the admixture dates for each target group listed on the X-axis. The inferred dates in generations were converted to dates in BCE by assuming a mean generation time of 28 years (4) and accounting for the average sampling age (shown as grey dots) of all ancient individuals in the target group (Methods). The top panel shows the formation of WHG–EHG cline (in blue) using Mesolithic hunter-gatherers as the target and EHG and WHG as reference populations. The middle panel shows admixture dates of local HGs and Anatolian farmers (in orange) using Neolithic European groups as targets and Anatolian farmers-related groups and WHG-related groups as reference populations. The bottom panel shows the spread of Steppe pastoralist-related ancestry (in green) estimated using middle and late Neolithic, Chalcolithic, and Bronze Age samples from Europe as target populations and early Steppe pastoralist-related groups (Afanasievo and Yamnaya Samara) and a set of Anatolian farmers and WHG-related groups as reference populations. For the middle to late Bronze Age samples from Eurasia, we used the early Steppe pastoralist-related groups and the Neolithic European groups as reference populations. The cultural affiliation (CWC, BBC, or Steppe MLBA cultures) of the individuals is shown in the legend.

#### Early to middle Neolithic

Neolithic farming began in the Near East—the Levant, Anatolia, and Iran—and spread to Europe and other parts of the world (18, 20, 27). The first farmers of Europe were related to Anatolian farmers, whose origin remains unclear. The early Neolithic Anatolian farmers (Aceramic Anatolian farmers) had majority ancestry from AHG with some gene flow from the first farmers from Iran (26). AHG, in turn, had ancestry from Levant HG (Natufians) and some mysterious hunter-gatherer group related to the ancestors of WHG individuals from central Europe— a gene flow event that likely occurred in the late Pleistocene (26). Using *qpAdm*, we confirmed that early Anatolian farmers could be modeled as a mixture of AHG and Iran Neolithic farmer-related groups (Note S5). To learn about the timing of the genetic formation of early Anatolian farmers, we applied *DATES* using one reference group as a set of pooled individuals of WHG-related and Levant Neolithic farmers-related individuals as a proxy of AHG ancestry and the second reference group containing pooled Iran Neolithic farmer-related individuals. We note that the application of *DATES* to three-way admixed groups can lead to intermediate dates between the first and second pulse of gene flow unless the reference populations are chosen carefully (Table S2.1). Our setup for early Anatolian farmers should have limited confounding and should recover the timing of the most recent event (in this case, the gene flow from CHG or Iran Neolithic-related groups) reliably (Table S2.1). We infer the Iran Neolithic farmer-related gene flow occurred ~10,900 BCE (12,200–9,600 BCE), predating the origin of farming in Anatolia (28). During the subsequent millennia, these early farmers further admixed with Levant Neolithic groups to form Anatolian Neolithic farmers who spread towards the west to Europe and in the east to mix with Iran Neolithic farmers, forming the Chalcolithic groups of Seh Gabi and Hajji Firuz. Using *DATES*, we inferred the Chalcolithic groups were genetically formed in ~7,600–5,700 BCE (Table SC).

In Europe, the Anatolian Neolithic farmers mixed with the local indigenous hunter-gatherers replacing between ~3-50% ancestry of Neolithic Europeans. To elucidate the fine-scale patterns and regional dynamics of these mixtures, we applied *DATES* to time transect samples from 94 groups (*n*=657) sampled from sixteen regions in Europe, ranging from ~6,000-1,900 BCE and encompassing individuals from the early Neolithic to Chalcolithic periods (Table SB). Using *qpAdm*, we first confirmed that the Neolithic Europeans could be modeled as a mixture of European hunter-gatherer-related ancestry and Anatolian farmer-related ancestry and inferred their ancestry proportions (Table SD). For most target populations (~80%), we found the model of gene flow between Anatolian farmer-related and WHG-related ancestry provided a good fit to the data (*p*-value > 0.05). In some populations, we found variation in the source of the HG-related ancestry and including either EHG or GoyetQ2 improved the fit of the model. In five groups, none of the models fit, despite excluding outlier individuals whose ancestry profile differed from the majority of the individuals in the group (Table SD, Table SE). To confirm that the target populations do not harbor Steppe pastoralist-related ancestry, we applied *D*-statistics of the form *D*(Mbuti, *target*, Anatolian farmers, Steppe pastoralists) where *target* = Neolithic European groups. We observed that four groups had a stronger affinity to Steppe pastoralists compared to Anatolian farmers, and hence we excluded these from further analysis (Table SF). After filtering, we applied *DATES* to 86 European Neolithic groups using WHG-related individuals and Anatolian farmers as reference populations.

Earlier analysis has suggested that farming spread along two main routes in Europe, from southeast to central Europe (‘continental route’) and along the Mediterranean coastline to Iberia (‘coastal route’) (23, 29, 30). Consistent with this, we inferred one of the earliest timings of gene flow in the Balkans around 6,400 BCE. Using the most comprehensive time-transect in Hungary with 19 groups (*n*=63) spanning from middle Neolithic to late Chalcolithic, we inferred that the admixture occurred between ~6,100–4,500 BCE. Under a model of a single shared gene flow event in the common ancestors of all individuals, we would expect to obtain similar dates of admixture (before present) after accounting for the age of the ancient specimens. Similar to Lipson et al. (2017), we observed that the estimated dates in middle Neolithic individuals were substantially older than those inferred in late Neolithic or Chalcolithic individuals (Figure 3). This would be expected if the underlying model of gene flow involved multiple pulses of gene flow, such that the timing in the middle Neolithic samples reflects the initial two-way mixture and the timing in the Chalcolithic samples captures both recent and older events. Interestingly, Lipson et al. (2017) and other recent studies have documented increasing HG ancestry from ~3-15% from the Neolithic to Chalcolithic period (16, 23, 31), suggesting that there was additional HG gene flow after the initial mixture. This highlights that the interactions between local hunter-gatherers and incoming Anatolian farmers were complex with multiple gene flow events between these two groups, which explains the increasing HG ancestry and more recent dates in Chalcolithic individuals (Table SD).

Mirroring the pattern in Hungary, we documented the resurgence of HG ancestry in the Czech Republic, France, Germany, and southern Europe. In central Europe, we inferred that the Anatolian farmer-related gene flow occurred ~5,600-5,000 BCE, with some exceptions. In the Blätterhöhle site from Germany, we inferred the gene flow occurred more recently (~4,000 BCE), consistent with the occupation of both hunter-gatherers and farmers in this region until the late Neolithic (31). In eastern Europe, using samples related to the Funnel Beaker culture (TRB; from German *Trichterbecher*) from Poland, we dated the Anatolian farmer-related gene flow occurred ~5,300–4,200 BCE. Following the TRB decline, the Baden culture and the Globular Amphora culture appeared in many areas of Poland and Ukraine (25). These cultures had close contacts with Corded Ware complex and steppe societies, though we did not find any evidence of Steppe pastoralist-related ancestry in the GAC individuals (Table SD). Applying *DATES*, we inferred the Anatolian farmer-related and HG mixture occurred ~5,200-3,100 BCE, predating the spread of Steppe pastoralists to eastern Europe (16, 19).

Along the Mediterranean route, we characterized Anatolian farmer-related gene flow in Italy, Iberia, France, and the British Isles. Using samples from five groups in Italy, we inferred the earliest dates of Anatolian farmer-related gene flow of ~6,100 BCE, and within the millennium, the ancestry spread from Sardinia to Sicily (Figure 3). In Iberia, the Anatolian farmer-related mixture occurred ~6,000–3,400 BCE and showed evidence for an increase in HG ancestry from ~9–20% after the initial gene flow. In France, previous studies have shown that Anatolian farmer-related ancestry came from both routes, along the Danubian in the north and along the Mediterranean in the south (23). This is reflected in the source of the HG ancestry, which is predominantly EHG and WHG-related in the north and includes WHG and Goyet-Q2 ancestry in the south (23). Consistently, we also observed that the admixture dates in France were structured along these routes, with the median estimate of ~5,100 BCE in the east and much older ~5,500 BCE in the south (Table SC). In Scandinavia, we inferred markedly more recent dates of admixture of ~4,300 BCE using samples from Sweden associated with the TRB culture and Ansarve Megalithic tombs, consistent with a late introduction of farming to Scandinavia (33).

Finally, we inferred recent dates of admixture in Neolithic samples from the British Isles (England, Scotland, and Ireland) with the median timing of ~5,000 BCE across the three regions. Interestingly, unlike in western and southern Europe, there was no resurgence in HG ancestry during the Neolithic in Britain (34). This suggests our dates can be interpreted as the time of the main mixture of HGs and Anatolian farmers in this region, implying that the farmer-related ancestry reached Britain a millennium after its arrival in continental Europe. By 4,300 BCE, we find that Anatolian farmer-related ancestry is present in nearly all regions in Europe.

#### Late Neolithic to Bronze Age

The beginning of the Bronze Age was a period of major cultural and demographic change in Eurasia, accompanied by the spread of Yamnaya Steppe Pastoralist-related ancestry from Pontic-Caspian steppes into Europe and South Asia (16). The archaeological record documents that the early Steppe pastoralists cultures of Yamnaya and Afanasievo, with characteristic burial styles and pottery, appeared around ~3,300 to 2,600 BCE (35). These groups were likely the result of a genetic admixture between the descendants of EHG-related groups and CHG-related groups associated with the first farmers from Iran (8, 22, 36). Using *qpAdm*, we first tested how well this model fits the data from 8 early Steppe pastoralist groups, including seven groups associated with Yamnaya culture and one group related to the Afanasievo culture (Methods). For all but two Yamnaya groups (from Hungary Baden and Russia Kalmykia), we found this model provides a good fit to the data (Table S5.4). We note that the samples from Kalmykia in our dataset were shotgun sequenced, and in the *qpAdm* analysis, we are mixing shotgun and capture data that could potentially lead to technical issues. To understand the timing of the formation of the early Steppe pastoralist-related groups, we applied *DATES* using pooled EHG and pooled Iranian Neolithic farmers. Focusing on the groups with the largest sample sizes, Yamnaya Samara (*n*=10) and Afanasievo (*n*=19), we inferred the admixture occurred between 40–45 generations before the individuals lived, translating to an admixture timing of ~4,100 BCE (Table S6.1). We obtained qualitatively similar dates across four Yamnaya and one Afanasievo groups, consistent with the findings that these groups descend from a recent common ancestor (for Ozera samples from Ukraine, the dates were not significant). This is also further supported by the insight that the genetic differentiation across early Steppe pastoralist groups is very low (*F_ST_* ~ 0.000-0.006) (Table S6.2). Thus, we combined all early Steppe pastoralist individuals in one group to obtain a more precise estimate for the genetic formation of proto-Yamnaya of ~4,400 to 4,000 BCE (Figure 2). These dates are noteworthy as they pre-date the archaeological evidence by more than a millennium (37) and have important implications for understanding the origin of proto-Pontic Caspian cultures and their spread to Europe and South Asia.

Over the following millennium, the Yamnaya-derived groups of the Corded Ware Complex (CWC) and Bell Beaker complex (BBC) cultures brought Steppe pastoralist-related ancestry to Europe. Present-day Europeans derive between ~10-60% Steppe pastoralist-related ancestry, which was not seen in Neolithic samples. To obtain a precise chronology of the spread of Steppe pastoralist-related ancestry across Europe, we analyzed 109 late Neolithic, Chalcolithic, and BA samples dated between 3,000-750 CE from 18 regions, including samples associated with the CWC and BBC cultures. We first confirmed that most target samples had Steppe pastoralist-related ancestry, in addition to European HG-related and Anatolian farmer-related ancestry using *qpAdm*. We excluded 20 groups that could not be parsimoniously modeled as a three-way mixture even after removing individual outliers. After filtering, we retained 79 groups for dating Steppe pastoralist-related gene flow across Europe (Note S5, Table SH). As Bronze Age Europeans have ancestry from three distinct groups, we applied *DATES* using the following two reference populations, one group including early Steppe pastoralists (Yamnaya and Afanasievo) and the other group with pooled samples of WHG-related and Anatolian farmer-related individuals, which is the proxy for the ancestral Neolithic Europe population.

To learn about the spread of CWC culture across Europe, we used seven late Neolithic and Bronze age groups, including five associated with CWC artifacts. Using *DATES*, we inferred that the oldest date of Steppe pastoralists gene flow in Europe was ~3,200 BCE in Scandinavia in samples associated with Battle Axe Culture in Sweden and Single Grave Culture in Denmark that were both contemporary to CWC. The samples from Scandinavia showed large heterogeneity in ancestry, including some individuals with majority Steppe pastoralist-related ancestry (and negligible amounts of Anatolian farmer-related ancestry), consistent with patterns expected from recent gene flow (38). Strikingly, we inferred the timing of admixture in central Europe (Germany and the Czech Republic) and eastern Europe (Estonia and Poland) to be remarkably similar. These dates fall within a narrow range of ~3,000–2,900 BCE across diverse regions, suggesting that the mixed population associated with the Corded Ware culture formed over a short time and spread across Europe rapidly with very little further mixture (Table SC).

Following the Corded Ware culture, from around 2,800 to 2,300 BCE, Bell Beaker pottery became widespread across Europe (39). Using 19 Chalcolithic and Bronze Age samples, including ten associated with Beaker-complex artifacts, we inferred the dynamics of the spread of the Beaker complex across Europe. We inferred the oldest date of Steppe pastoralist-related admixture was ~3,200 BCE (3600–2800 BCE) in EBA Mallorca samples from Iberia. We note the EBA Mallorca sample is not directly associated with Beaker culture, but *qpAdm* modeling suggests that this individual is clade with the small subset of Iberian Beaker-complex-associated individuals who carried Steppe pastoralist-related (40). Most individuals from Iberia, however, had negligible Steppe pastoralist-related ancestry suggesting the Beaker culture was not accompanied by major gene flow in Iberia despite the earliest dates (Table SH). In central and western Europe, where steppe gene flow was more pervasive, we inferred the median date of the mixture was ~2,700 BCE with the oldest dates in the Netherlands, followed by Germany and France (Figure 3). There was, however, large heterogeneity in the dates across Europe and even within the same region. For example, comparing two BA groups from the Netherlands suggests a wide range of dates ~3,000 BCE and 2,500 BCE, and four groups from Germany indicate a range of ~2,900–2,700 BCE. From central Europe, the Steppe pastoralist-related ancestry spread quickly to the British Isles, where people with steppe ancestry replaced 90% of the genetic ancestry of individuals from Britain. Our estimates for the time of gene flow in Bell Beakers samples from England suggest that the gene flow occurred ~2,700 BCE (2770-2550 BCE). Our estimated dates of admixture are older than the dates of arrival of this ancestry in Britain (41) and, interestingly, overlap the dates in central Europe. Given that a significant fraction of the Beaker individuals were recent migrants from central Europe, we interpret our dates reflect the admixture into ancestors of the British Beaker people, occurring in mainland Europe (41).

The middle to late Bronze age led to the final integration of Steppe pastoralist-related ancestry in Europe. In southern Europe, early BA samples had limited Steppe pastoralist-related ancestry, though present-day individuals have between ~5–30% steppe ancestry (16). Using pooled samples of middle to late BA from Spain, we inferred major mixture occurred ~2,500 BCE in Iberia. We inferred a similar timing in Italy using individuals associated with the Bell Beaker culture and early BA samples from Sicily (Table SC). In Sardinia, a majority of the BA samples do not have Steppe pastoralist-related ancestry. In a few individuals, we found evidence for steppe ancestry, though in most cases, the Steppe pastoralist-related ancestry proportion overlapped 0, and the dates were very noisy (Table SH). Using Iron Age samples from Sardinia, we inferred the gene flow occurred ~2,600 BCE, though there is large uncertainty associated with this estimate (2,614 +/− 560 BCE). In other parts of continental Europe and the British Isles, the Steppe pastoralist-related gene flow got diluted over time, as evidenced by more recent dates in LBA than EBA or MBA samples in Germany, England, and Scotland, and increase in Neolithic farmer ancestry during this period (42) (Table SC).

Finally, the Corded Ware Complex expanded to the east to form the archaeological complexes of Sintashta, Srubnaya, Andronovo, and the Bronze Age cultures of Kazakhstan. Samples associated with these cultures harbor mixed ancestry from the Yamnaya Steppe pastoralist-related groups (CWC, in some cases) and Neolithic individuals from central Europe (Table S5.5) (8). Applying *DATES* to 8 Middle to late Bronze Age (MLBA) Steppe pastoralist groups, we inferred the precise timing for the formation of these groups beginning in the third millennium BCE. These groups were formed chronologically, with the date of genetic formation of ~3,200 BCE for Sintashta culture, followed by ~2,900 BCE for Srubnaya and Andronovo cultures. In the central Steppe region (present-day Kazakhstan), we obtained median dates of ~2,800 BCE for the expansion of Steppe pastoralist-related ancestry in four Kazakh cultures of Maitan Alakul, Aktogai, and Kairan. By ~2,700 BCE, most of these cultures had almost 60-70% Yamnaya Steppe pastoralist-related ancestry (Table SC). These groups, in turn, expanded eastwards, transforming the genetic composition of populations in South Asia.

## Discussion

We developed *DATES*, a novel method to measure ancestry covariance in a single diploid individual genome to estimate the time of admixture. Using extensive simulations, we show that *DATES* provides accurate estimates of the timing of admixture for a range of demographic scenarios. Application of *DATES* to present-day samples shows that the results are concordant with published methods—Rolloff, ALDER, and Globetrotter. For sparse datasets, *DATES* outperforms published methods as it does not require phased data and works reliably with limited samples, large proportions of missing variants as well as pseudodiploid genotypes. This makes *DATES* ideally suited for the analysis of ancient DNA samples.

We illustrate the application of *DATES* by reconstructing population movements and admixtures during the European Holocene. The European continent was subject to two major migrations during the Holocene: the movement of Anatolian farmers during the Neolithic and the migration of Yamnaya Steppe pastoralists during the Bronze Age. First, we document that the Mesolithic hunter-gatherers formed as a mixture of WHG and EHG ancestry ~10,200 to 7400 BCE. These dates are consistent with the archeological evidence for the appearance of lithic technology associated with eastern HGs in Scandinavia and the Baltic regions and the spread of WHG ancestry to east (17, 43, 44). Next, we studied the timing of the genetic formation of Anatolian farmers. The earliest evidence of agriculture comes from the Fertile Crescent, the southern Levant, and the Zagros Mountains of Iran and dates to around 10,000 BCE. In central Anatolia, farming has been documented c. 8,300 BCE (45, 46). It has been long debated if Neolithic farming groups from Iran and the Levant introduced agriculture to Anatolia or hunter-gatherers in the region locally adopted agricultural practices. The early Anatolian farmers can be modeled as a mixture of local hunter-gatherers people related to Caucasus hunter-gatherers or first farmers from Iran (26). By applying *DATES* (assuming single instantaneous admixture), we inferred that the Iran Neolithic gene flow occurred around 10,900 BCE (~12,200–9,600 BCE). An alternate possibility is that there was a long period of gradual gene flow between the two groups and our dates reflect intermediate dates between the start and end of the gene flow. An upper bound for such mixture comes from the lack of Iran Neolithic ancestry in Anatolian HGs at 13,000 BCE, and a lower bound comes from the C14 dates of early Anatolian farmers, one of which is directly dated at 8269–8210 BCE (26). In either case (instantaneous admixture or gradual gene flow), the genetic mixture that formed Anatolian farmers predates the advent of agriculture in this region. This supports the model that Anatolian hunter-gatherers locally transitioned to agricultural subsistence, and most probably, there was cultural diffusion from other regions in Near East (Iran and Levant) (26). Future studies with more dense temporal sampling will shed light on the demographic processes that led to the transition from foraging to farming in the Near East, and in turn, elucidate the relative roles of demic and cultural diffusion in the dispersal of technologies like agriculture across populations.

Using data from sixteen regions in Europe, we reconstruct a detailed chronology and dynamics of the expansion and admixture of Anatolian farmers during the Neolithic period. We infer that starting in ~6,400 BCE, gene flow from Anatolian farmers became widespread across Europe, and by ~4,300 BCE, it was present in almost all parts of continental Europe and the British Isles. These dates are significantly more recent than the estimates of farming based on archaeological evidence in some parts of Europe, suggesting that the local hunter-gatherers and farmers co-existed for more than a millennium before the mixture occurred (16, 31). In many regions, after the initial mixture, there was a resurgence of HG ancestry, highlighting the complexities of these ancient interactions. We note that our results are consistent with two previous genetic studies, Lipson et al. (2017) and Rivollat et al. (2020), that applied genetic dating methods to a subset of samples we used in our analysis. Lipson et al. (2017) used a modified version of ALDER to infer the timing of admixture in three regions (*n*=151), and we obtained statistically consistent results for all overlapping samples (within two standard errors). An advantage of our approach over the modified ALDER approach is that we do not rely on helper samples (higher coverage individuals combined with the target group) for dating; unless these have a similar ancestry profile, they could bias the inferred dates. Our results are concordant with Rivollat et al. (2020) that used a previous version of *DATES* to infer the timing of Neolithic gene flow in 32 groups (vs. 86 groups in our study). We find the performance of both versions of *DATES* is similar, though some implementation details have improved (see Note S3, Table S3.3).

The second major migration occurred when populations associated with the Yamnaya culture in the Pontic-Caspian steppe expanded to central and western Europe from far eastern Europe. Our analysis reveals the precise timing of the genetic formation of these early Steppe pastoralists groups–Yamnaya and Afanasievo–occurred ~4,400-4,000 BCE. This estimate predates the archaeological evidence by more than a millennium (37) and suggests the presence of an ancient “ghost” population of proto-Yamnaya around this time. Understanding the source and location of this ghost population will provide deep insights into the history of Pontic-Caspian cultures and the origin of Indo-European languages that have been associated to have spread with Steppe pastoralists ancestry to Europe and South Asia (16, 47). Starting in ~3,200 BCE, the Yamnaya-derived cultures of Corded Ware Complex and Bell Beaker complex spread westwards, bringing steppe ancestry to Europe. Our analysis reveals striking differences in the spread of these three cultures: the Yamnaya were genetically formed a millennium before the evidence for pastoralism, while CWC formation is coincident with the archaeological dates and similar across diverse regions separated by thousands of kilometers, suggesting a rapid spread after the initial formation of this group. In contrast, the formation and expansion of people with Steppe pastoralist-related ancestry associated with Bell Beakers cultural artifacts are much more complex and heterogeneous across regions. We find the earliest evidence of Steppe pastoralist-related ancestry in Iberia around 3200 BCE, though this ancestry only becomes widespread after 2,500 BCE. In central Europe, the gene flow occurred simultaneously with archaeological evidence and was coexisting with the Corded Ware complex in some parts (41, 48). Finally, in the British Isles, the Bell Beaker culture spreads rapidly from central Europe and replaces almost 90% of the ancestry of individuals in this region (41).

Recent analysis has shown remarkable parallels in the history of Europe and South Asia; with both groups deriving ancestry from local indigenous HGs, Near Eastern farmers, and Steppe pastoralist-related groups (8). Interestingly, however, the timing of the two major migrations events differs across the two subcontinents. Both mixtures occurred in Europe almost a millennium before they occurred in South Asia. In Europe, the Neolithic migrations primarily involved Anatolian farmers, while the source of Neolithic ancestry is closer to Iran Neolithic farmers in South Asia. The Steppe pastoralist-related gene flow occurred in the context of the spread of CWC and BBC cultures in Europe around 3,200-2,500 BCE; in South Asia, this ancestry arrived with Steppe MLBA cultures in 1,800-1,500 BCE (8). The Steppe MLBA groups were genetically formed as an admixture of Steppe pastoralist-derived groups and European Neolithic farmers following the eastward expansion of CWC groups between ~3,200–2,700 BCE. Understanding the origin and migration paths of the ancestral groups thus helps to illuminate the differences in the timeline of the spread of steppe genetics across the two subcontinents of Eurasia.

Genomic dating methods like *DATES* provide an independent and complementary approach for reconstructing population history. By focusing on genetic clocks like recombination rate, we provide an independent estimate of the timing of evolutionary events up to several thousands of years. Our analysis also has advantages over temporal sampling of ancient DNA, in that we can obtain direct estimates of when a population was formed, rather than inferring putative bounds for the timing based on the absence/presence of a particular ancestry signature (which may be sensitive to sampling choice and density). Genetic approaches provide complementary evidence to archaeology and linguistics as they date the time of gene flow and not migration. Both dates are similar in many contemporary populations like African Americans and Latinos, though this may not be generally true (2). This is underscored by our dates for the Neolithic farmer mixture, which post-dates evidence of material culture related to agriculture by almost two millennia in some regions. This suggests that European HGs and farmers resided side by side for several thousand years before gene exchange (49, 50). This highlights how genetic dates can provide complementary evidence to archaeology and help to build a comprehensive picture of population origins and movements.

## Methods and Materials

### Dataset

We analyzed 1,096 ancient European samples from 152 groups restricting to data from 1,233,013 autosomal SNP positions that were genotyped using the Affymetrix Human Origins array (the V44.3 release of the Allen Ancient DNA Resource (AADR); https://reich.hms.harvard.edu/allen-ancient-dna-resource-aadr-downloadable-genotypes-present-day-and-ancient-dna-data). We filtered this dataset to remove samples that were marked as contaminated, low coverage, outliers, duplicates or first- or second-degree relatives (Table SB). We grouped individuals together from a particular culture or region. Details of sample affiliation and grouping used is described in Table SA.

### Modeling admixture history

We applied *qpAdm* from ADMIXTOOLS to identify the best fitting model and estimate the ancestry proportions in a target population modeled as a mixture of *n* “reference” populations using a set of “Outgroup” populations (16). We set the details: YES parameter, which reports a normally distributed Z-score to evaluate the goodness of fit of the model (estimated with a Block Jackknife). For each target population, we chose the most parsimonious model, i.e., fitting the data with the minimum number of source populations. We excluded models where the *p*-value < 0.05 indicating a poor fit to the data. Details of the *qpAdm* analysis for each group are reported in Note S5. We also applied *D*-statistics in some cases using *qpDstat* in ADMIXTOOLS with default parameters.

### Dating admixture events

We applied *DATES* to infer the time of admixture for a given target population. We present the details of the model and implementation in Note S1. We applied *DATES* using genome-wide SNP data from the target population and two reference populations. To infer the allele frequency in the ancestral populations more reliably, where specified, we pooled individuals deriving the majority of their ancestry from the population of interest (Table SA). We computed the weighted ancestry covariance between 0.45cM (to minimize the impact of background LD) to 100 cM, with a bin size of 0.1 cM. We plotted the weighted covariance with genetic distance and obtained a date by fitting an exponential function with an affine term *y* = *Ae^−λd^* + *c*, where *d* is the genetic distance in Morgans and *λ* = (*t*+1) is the number of generations since admixture (*t*). The factor of (*t*+1) is because in the first-generation following admixture, the admixed population derives one chromosome from each ancestral group. The mixing of chromosomes only begins in the following generations as the chromosomes recombine. We computed standard errors using weighted block jackknife, where one chromosome was removed in each run (51). We examined the quality of the exponential fit by computing the normalized root-mean-square deviation (NRMSD) between the empirical ancestry covariance values *z* and the fitted ones *ẑ*, across all the genetic distance bins (11).

The estimated dates of admixture were considered <u>significant</u> if the *Z*-score > 2, *λ* < 200 generations and NRMSD < 0.7. We converted the inferred dates from generations to years by assuming a mean generation time of 28 years (1). For ancient samples, we added the sampling age of the ancient specimen (Table SA). When multiple individuals were available, we used the average sampling ages to offset the admixture dates. We report dates in BCE by assuming the 1950 convention.

## Supporting information

Chintalapati_etal_SI_Material

Chintalapati_etal_DataTables

## Software availability

The executable and source code for *DATES* will be available on GitHub: https://github.com/MoorjaniLab/DATES_v3600

## Acknowledgments

We thank Monty Slatkin, Ziyue Gao, David Reich, Iosif Lazaridis, and Vagheesh Narasimhan for their comments on the manuscript. We thank Iosif Lazaridis for helpful discussions about population models in the Near East, Remi Tournebize for suggestions for evaluating the fit of exponential decay curves, and Neel Alex for suggestions for implementation of FFT for an earlier version of DATES. We are grateful for support from the Burroughs Wellcome Fund Careers at the Scientific Interface, Sloan Research Fellowship, and NIH R35GM142978 awarded to PM. NP was a fellow at the Radcliffe Institute for Advanced Study at Harvard University.

## Notes

### Competing Interest Statement

The authors have declared no competing interest.

