## Supplementary material for "Reconstructing the spatiotemporal patterns of admixture during the European Holocene using a novel genomic dating method": Chintalapati_etal_SI_Material

#### Table of Contents

**Supplementary Notes**


Note S1. *DATES*: model and implementation 2

Note S2. Simulations to test the performance of *DATES* 8

Note S3. Comparison of versions of *DATES* 40

Note S4. Comparison to other published methods 45

Note S5. Modeling population structure in ancient Eurasia 49

Note S6. Formation of early Steppe pastoralists groups 55

References 57


**Supplementary Figures**


Figure SA. Timing of WHG and EHG admixture in Iron Gates samples 59

Figure SB. Decay curves for all ancient samples included in the study 59


**Supplementary Tables** – Included in separate excel spreadsheet.

### Note S1. *DATES*: model and implementation

*DATES* leverages the weighted ancestry covariance patterns across the genome of an admixed individual to infer the time of admixture. This method extends the idea introduced in ROLLOFF and ALDER and ref. (1) to be applicable to dating admixture events between modern human populations using a single genome (1).

*Basic model and notation*

Assume we have an admixed individual *C* with ancestry from source populations *A* and *B*, with ancestry proportion of $\alpha$ and $\beta = (1-\alpha)$ respectively. This mixture occurred $t$ generations ago. First, we model the genotypes of *C* as a linear mix of allele frequencies of populations *A* and *B*. For any SNP $i$, let the genotype of *C* be $g_i$ and allele frequency in *A* and *B* be $p_A(i)$ and $p_B(i)$. We can then infer the mixing fraction $\alpha$ from population *A* by solving the simple linear regression by minimizing the residuals.

$$R = \sum_i (g_i - (\alpha p_A(i) + (1-\alpha) p_B(i)))^2 \quad (1)$$

Let $a_i$ be the probability of observing $g_i$ in *C* given the observed genotype in *A,* and $b_i$ be the probability of observing $g_i$ in *C* given the observed genotype in *B*

$$a_i = P(g_i|A)$$
$$b_i = P(g_i|B)$$

We can then compute the likelihood $L_i$ of observing a genotype $g_i$ in the admixed individual

$$L_i = \alpha a_i + \beta b_i \quad (2)$$

For a pair of neighboring markers $S_1$, $S_2$ located at a genetic distance of *d* Morgans, the probability of no recombination between the two markers is given by $\theta = e^{-td}$. Accounting for recombination, the log likelihood that the two markers have the same ancestry is then given by:

$$\mathcal{L} = \log[(1-\theta) L_1 L_2 + \theta(\alpha a_1 a_2 + \beta b_1 b_2)] \quad (3)$$

Let $K_i$ represent the ancestry at marker $S_i$. Expanding as a power series in $\theta$, the coefficient of $\theta$ is $QK_1K_2$, where

$$Q = \alpha\beta$$
$$K_i = \frac{(a_i - b_i)}{L_i} \quad (4)$$

We can compute the ancestry covariance, $A(d)$, across pairs of markers $S_1$, $S_2$ separated by distance *d* as

$$A(d) = \frac{\sum_{S(d)} (K_1 - \overline{K_1})(K_2 - \overline{K_2})}{|S(d)|}$$

where $S(d)$ is a set of markers $S_1$, $S_2$ located *d* Morgans apart.

The ancestry covariance $A(d)$ is expected to follow an exponential decay with *d* with the rate of decay depending on the time since admixture $(t)$

$$A(d) \sim e^{-(t+1)d}$$

The factor of ($t$+1) is because, in the first-generation following admixture, the admixed population derives one chromosome from each ancestral group. The mixing of chromosomes only begins in the following generations as the chromosomes recombine. This means that if we fit *t* generations, we are likely to underestimate the time of admixture. We note that previous methods like ALDER and ROLLOFF, however, incorrectly fit $t$ generations to infer the time of mixture. In practice, however, this has little effect on the inference except for every recent admixture dates. We infer the time of the mixture by fitting an exponential distribution with affine term using least squares. *DATES* is applicable for dating admixture in a single individual. When multiple individuals from an admixed population are available, *DATES* computes the log-likelihood by summing over all individuals.

*Implementation*

In order to make *DATES* computationally tractable, we implemented the fast Fourier transform (FFT) for computing ancestry covariance as described in ALDER (2). Briefly, we perform an algebraic transformation of the ancestry covariance statistic and compute the FFT convolution in discrete equally sized bins (referred to as mesh points). This provides a speedup from $O(n^2)$ to $O(n \log n\ )$, which reduces the typical runtimes from hours to seconds.

In *DATES*, we compute the ancestry covariance $A(d)$ by expanding the numerator as follows:

$$X_2 = \sum_{s(d)} K_1 K_2 X_1(d) = 2 \sum_{s(d)} \overline{K_1} K_2 \mid \overline{K_2} K_1 X_0(d) = \sum_{s(d)} \overline{K_1}\, \overline{K_2}$$

We discuss an approximate calculation of $X_2$. The other terms are similar to ALDER (2).

Like ALDER, we divide the genome in windows based on the position in the genetic map (instead of genetic distance). We set a mesh on the genetic map (default mesh size is 0.01 centiMorgans (cM)), mapping every SNP to the nearest mesh point. For a mesh point, $u$ define $T_u$ to be the set of SNPs mapping to $u$ and

$$K(u) = \sum_{i \in T(u} K_i$$

We now set

$$X_2' = \sum_{u,v \mid u-v \mid = d} K(u) K(v)$$

Where $|u - v|$ is the genetic distance of $u, v$. $X_2'$ can be computed by FFT. We note that the use of the mesh is the only source of approximation in the FFT implementation to compute $X_2$. The mesh discretization parameter, *qbin*, provides a trade-off between runtime and accuracy, smaller mesh size leads to higher accuracy and longer run time.

To explore the impact of *qbin* on the estimated accuracy, we performed simulations for varying sample sizes (*n*=1 and *n*=20) and ran *DATES* using varying *qbin* between 1–100. We find the method works reliably for all *qbin* values (Figure S1.1). Moreover, there is almost a 5–10-fold speedup in a run between *qbin* values of 10 vs. 100 (Figure S1.2). The run time is invariant to the proportion of time of admixture. The default value of *qbin* in *DATES* is 10 but we advise the user to perform simulations for their dataset size and population model to set this parameter reliably.

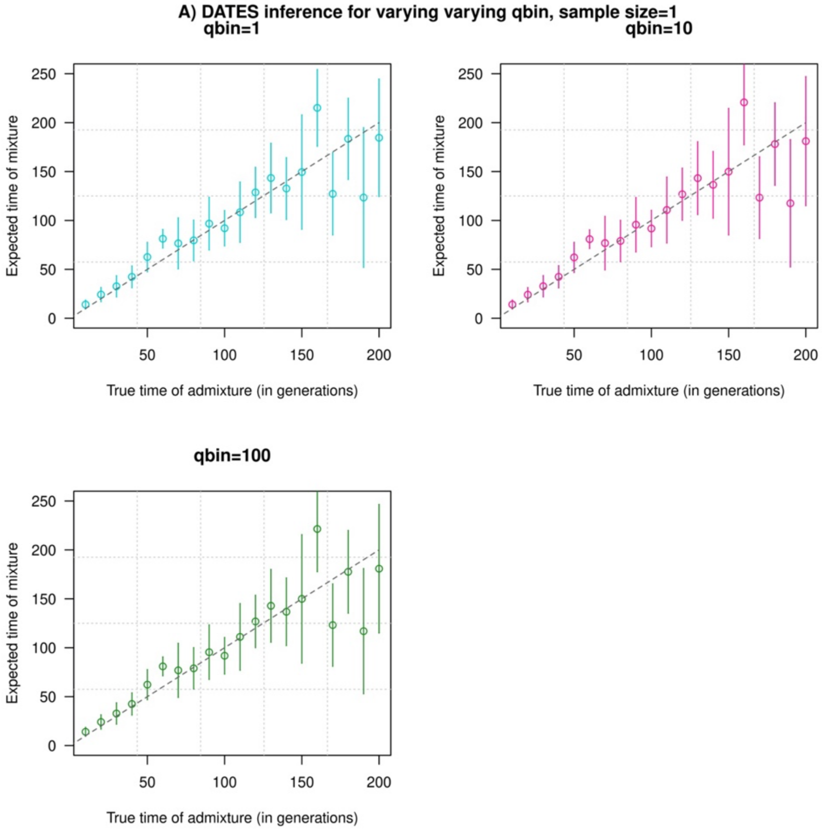

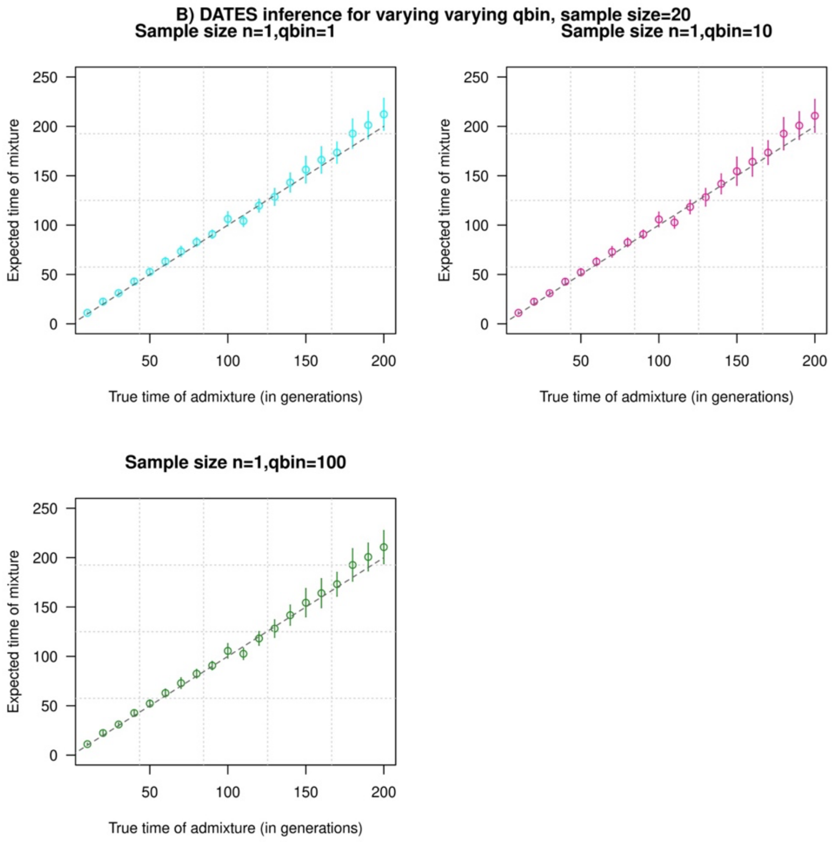


**Figure S1.1. Impact of the discretization parameter (*qbin*) on accuracy.** We show three subplots for a sample size of *n*=1 (Panel A) and *n*=20 (Panel B). For each subplot, we simulated data for *n* admixed individuals with 20% ancestry from Europeans (1000 Genomes, CEU) and 80% ancestry from Africans (1000 Genomes, YRI) with the time of admixture (*t*) shown on the X-axis and the estimated admixture time inferred using *DATES* on Y-axis. We ran *DATES* using varying *qbin* values shown in different colors.

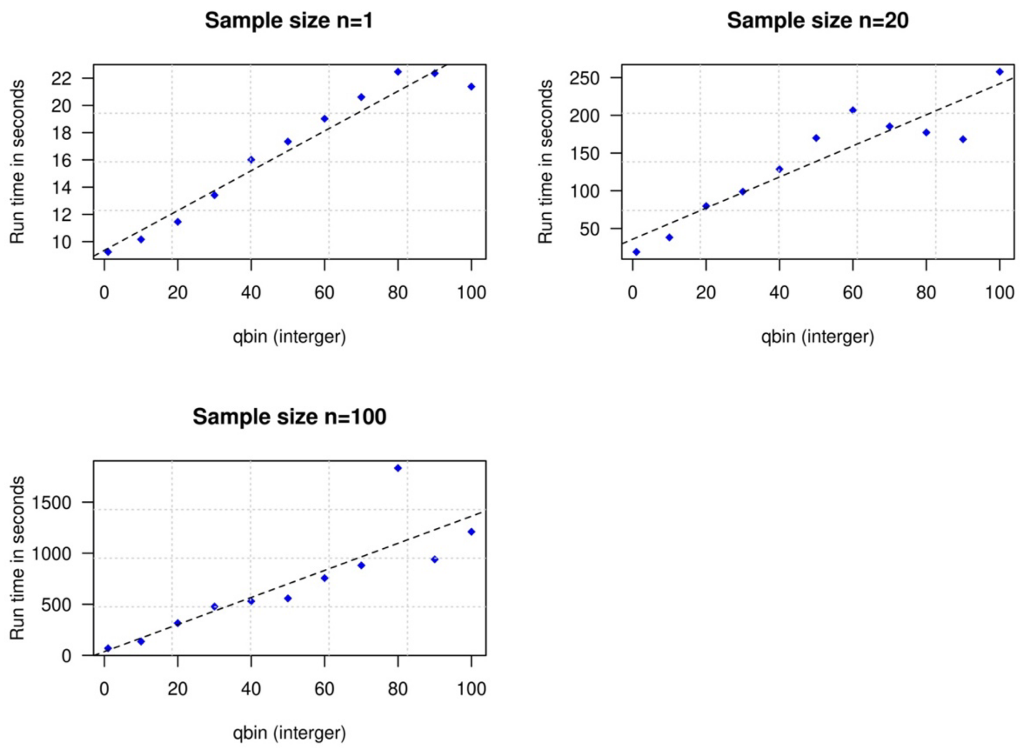


**Figure S1.2. Impact of the discretization parameter (*qbin*) on run time.** We show three subplots for sample size of *n=1* (top left), *n=20* (top right) and *n=100* (bottom). For each subplot, we simulated data for *n* admixed individuals with 20% ancestry from Europeans (1000 Genomes, CEU) and 80% ancestry from Africans (1000 Genomes, YRI) with the time of admixture (*t*) of 100 generations ago. We show the impact of *qbin* (X-axis) on the runtime measured in seconds. For sample sizes, $n > 1$, $r^2$ between *qbin* and run time is >0.99.

*Assessing the exponential fit*

Following Tournebize et al. (2020), we examined the quality of the exponential fit by computing the normalized root-mean-square deviation (NRMSD) between the empirical ancestry covariance values $z$ and the fitted ones $\hat{z}$, across all the genetic distance bins (where, $N$ the number of bins) (3).

$$NRMSD = \frac{1}{max(\hat{z}) - min(\hat{z})} \sqrt{\frac{\sum^{D} (z - \hat{z})^2}{N}}$$

We calculated NRMSD for all ancient DNA populations in our study and the distribution of these values is shown in (Figure S1.3). Focusing on the most extreme values of NRMSD, we show that the statistic is useful in identifying poor fits where the fitted line deviates from the data or the fit is highly dispersed (Figure 1.4). However, the absolute value of this statistic does not have any statistical meaning. Based on the empirical distribution of NRMSD values in our study samples (Figure 1.3), we use a threshold of 0.7 to flag poor fits. We caution that users should adjust this threshold based on their application and always visually inspect their exponential fits to ensure reliable results.

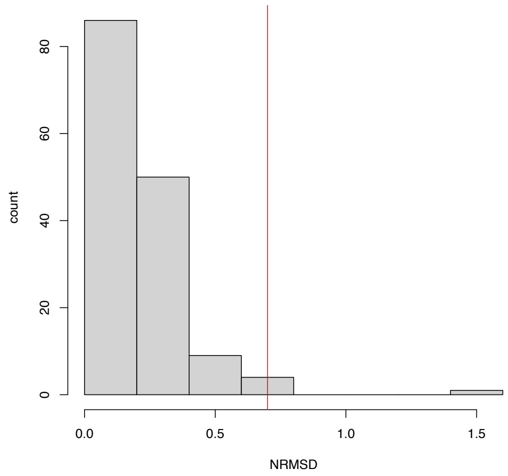


**Figure S1.3. Histogram of the NRMSD values computed as the normalized residual between the empirical and fitted decay curves, for all the ancient DNA populations reported in Supplementary Table SB.** The red vertical line represents the value NRMSD=0.7, which we used as the threshold to exclude populations from our analysis because visual inspection of fitted curves above this threshold suggests the results are too noisy to make a reliable inference (see Figure SA for all fitted decay curves).

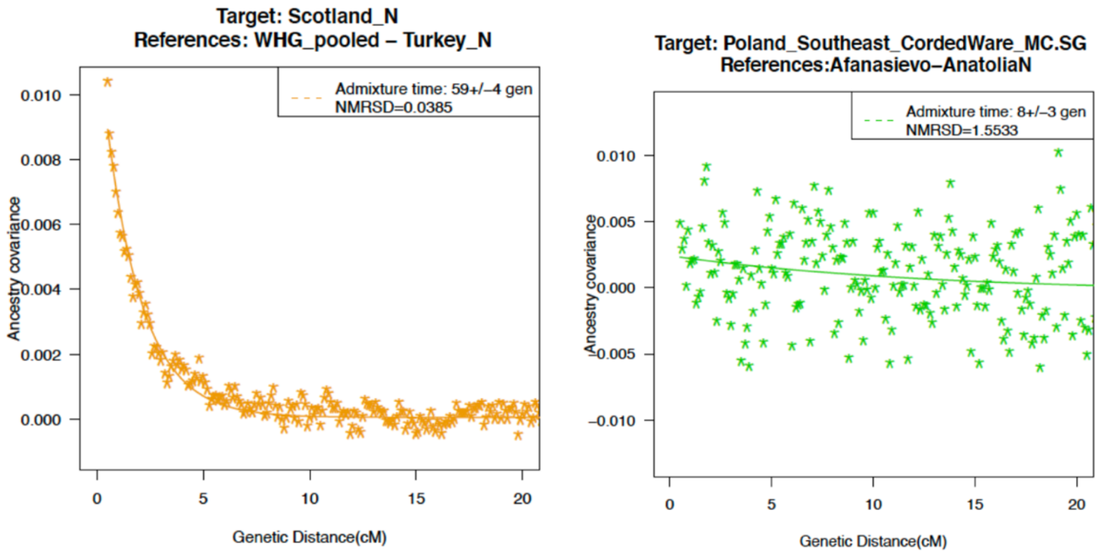


**Figure S1.4. Ancestry covariance curves for the lowest (left) and highest (right) values in our ancient DNA populations in our study**. For details of all curves and NRMSD, see Supplementary Table SB and Supplementary Figure S7.

### Note S2. Simulations to test the performance of *DATES*

To test the performance of *DATES*, we simulated data for admixed groups for a range of demographic scenarios and parameters that we detail below.

##### A. Haplotype-based simulations

We constructed admixed genomes following the procedure described in Moorjani et al. (2011) that simulates an admixed genome based on two parameters: (a) the mixture proportion ($\alpha$) that represents the probability that a particular sampled haplotype comes from one of the reference panels, namely $source_1$ and $source_2$, and (b) the time of mixture ($t$) which is the number of generations since mixture. To simulate an admixed individual, we begin at the start of the chromosome and sample a haplotype from either $source_1$ with a probability ($\alpha$) and $source_2$ with a probability $(1-\alpha)$. At each subsequent marker, we check if there was a recombination event between the two neighboring markers. A recombination event occurs with a probability of $(1-e^{-\lambda g})$, where $g$ is the genetic distance in Morgans. We use the time of $\lambda = (t+1)$ generations to account for the fact that in the first-generation following admixture, the offspring inherits one chromosome of each ancestry. In the next generation, the crossovers lead to a mixing of ancestry. Thus, when a recombination event occurs, we resample the ancestry between $source_1$ or $source_2$, otherwise, we copy the haplotype from the same source population (Note, a recombination event can lead to a switch to a haplotype of the same ancestry). Once the ancestry is chosen, we randomly pick a haplotype from the ancestral pool (without replacement) and copy its sequence to the genome of the admixed individual. This process is continued until we reach the end of the chromosome. Using this approach, we generate the genomes of $n$ admixed individuals. The simulated haploid chromosomes are merged at random to construct diploid admixed individuals. This algorithm requires more than $2n$ ancestral haplotypes for generating data for $n$ diploid admixed individuals (4). For more than two reference populations, the same algorithm is repeated iteratively.

For the simulations described below and in the main text, we used phased haplotype data from the 1000 Genomes Project (1000G) for 111 Europeans (Utah Residents CEPH with Northern European ancestry (CEU)), 88 East-Asians (Han Chinese in Beijing, China (CHB)), and 112 west Africans (Yoruba in Ibadan, Nigeria (YRI)) (5). Unless otherwise stated, we simulated data for 10 admixed individuals with ~380,000 SNPs with 20% European and 80% African ancestry where the time of admixture varied between 10–200 generations ago. For inference, we used French and Yoruba individuals from the Human Genome Diversity Panel (HGDP) (6) as the reference populations to represent the ancestral populations of CEU and YRI respectively. Standard errors were generated using a weighted block jackknife, where one chromosome was removed in each run (7). We evaluated the performance of *DATES* under the following scenarios:

#### 1. Varying the time of admixture

We investigated the accuracy of *DATES* for inferring the time of admixture by generating data for a range of admixture times, between 10–300 generations (in increments of 10 generations). We found that *DATES* provides accurate estimates up to 300 generations, though the precision is higher for dates below 200 generations (Figure S2.1). Thus, we focus on dates below 200 generations henceforth and in the main text.

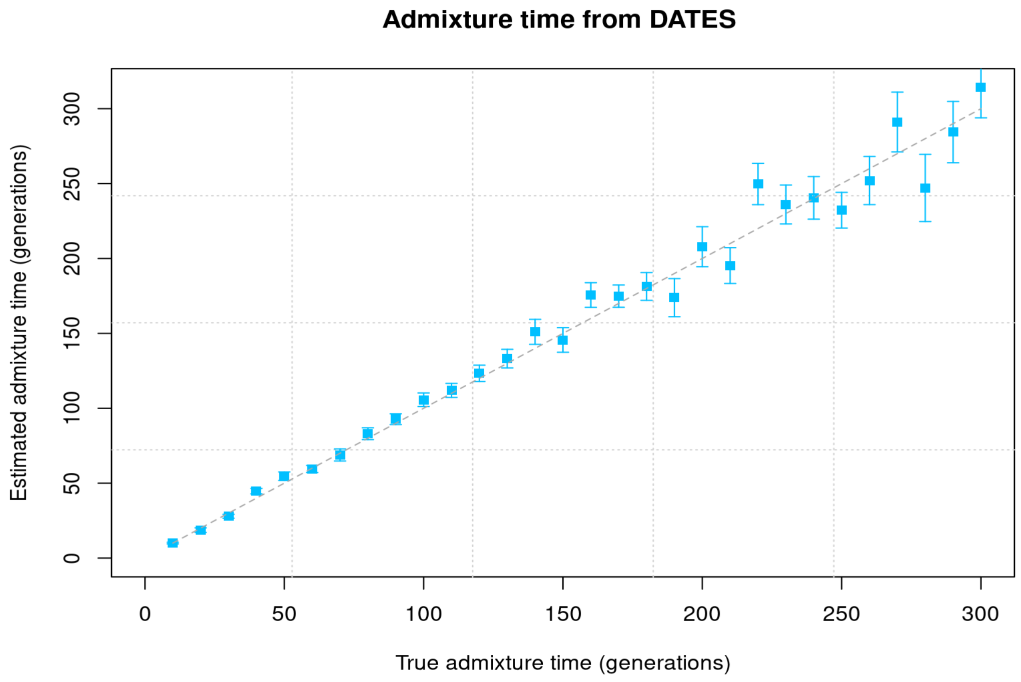


**Figure S2.1: Varying admixture time.** We simulated data for 10 admixed individuals with 20%/80% CEU and YRI ancestry respectively and varied the admixture times between 10 to 300 generations. The X-axis shows the true time of admixture, and the Y-axis shows the estimated time of admixture (± 1 SE) inferred using *DATES*.

#### 2. Varying the proportion of admixture

We explored the impact of admixture proportion on estimating the inferred time and admixture proportion by generating data for admixed groups with varying proportions of European ancestry between 1–50% (with the rest of the ancestry derived from Africans (YRI)). We found that the estimated dates were reliable, regardless of the admixture proportion, even for cases with only 1% European ancestry (Figure S2.2A).

*DATES* also estimates the ancestry proportion based on a simple regression analysis by modeling the observed genotypes in an admixed individual as a linear mix of allele frequencies from two reference populations (see Methods). To test the reliability of this simple approach (we do not account for the drift in the reference and target groups), we compared the inferred European ancestry proportion with the true ancestry proportion for the above-mentioned simulations where European ancestry varied between 1–50%. We observed that *DATES* accurately estimates the ancestry proportion for groups with >10% European ancestry, though for lower admixture proportions it tends to overestimate the ancestry proportion (Figure S2.2B). Thus, for low proportions of admixture, we suggest the users should use other methods such as *qpAdm* or $f_4$*-ratio* test that work reliably even with low ancestry proportions (8, 9).

**A) Admixture time inference for varying admixture proportions**

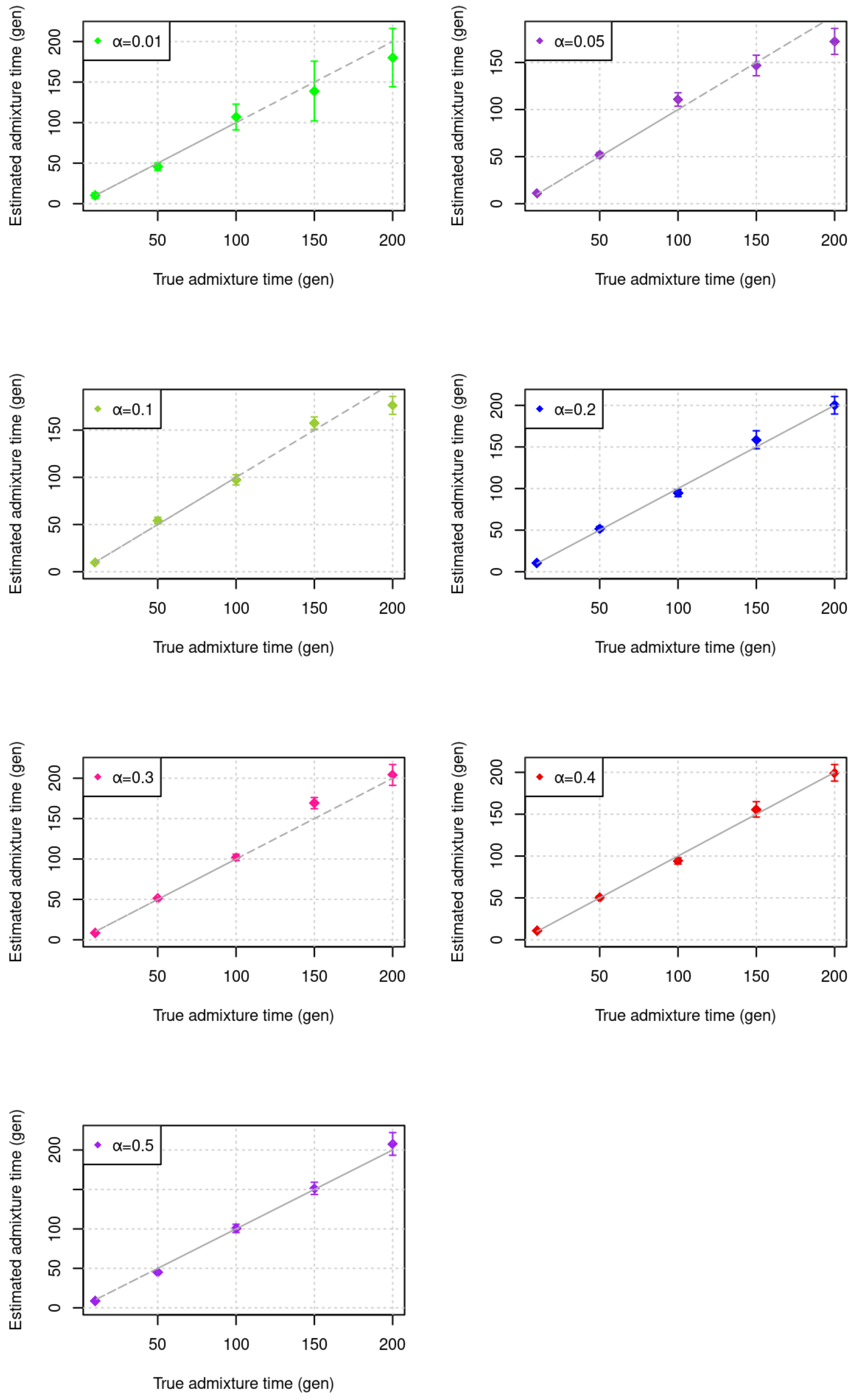

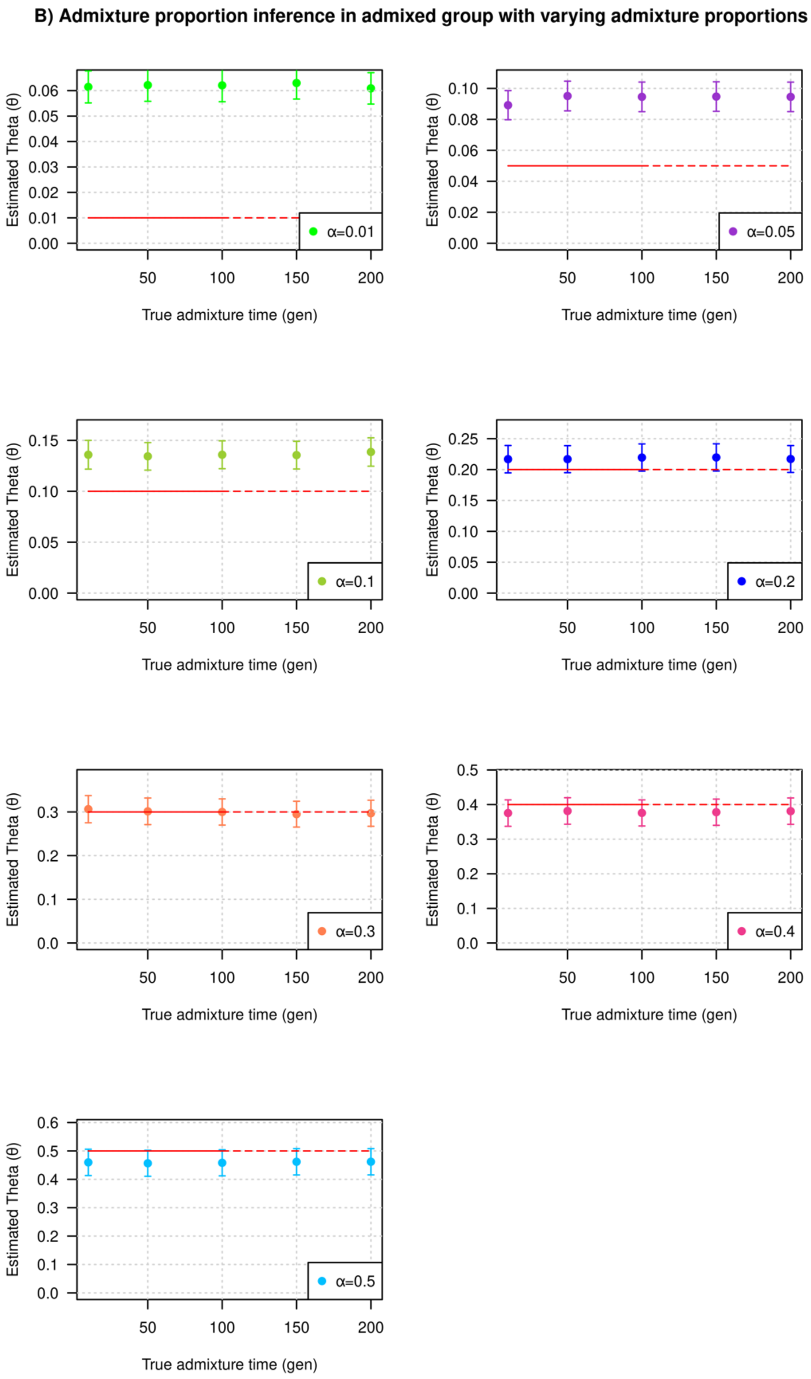


**Figure S2.2: Impact of variation in admixture proportion on the inference.** We simulated data for 10 admixed individuals with European ancestry ($\alpha$) in the range of 1-50% (the rest of the ancestry is derived from Africans). We ran *DATES* to infer the time of admixture and ancestry proportion. For each value of $\alpha$ (shown in different colors), we show A) Impact on the estimated time of admixture: The estimated time of admixture (± 1 SE) is shown on the Y-axis and the true time of admixture is shown on the X-axis; B) Impact on estimated ancestry proportion: The true time of admixture is shown on X-axis with the inferred proportion of admixture on Y-axis. The true admixture proportion used in the simulation is shown as a red dashed line and indicated in the legend.

#### 3. Varying sample size

We investigated the impact of sample size on the inference of admixture time. Because *DATES* uses allele frequencies in the ancestral populations for computing weighted ancestry covariance in the target, the sample size of the reference populations can also impact the estimation of the dates in the target (see Methods). Thus, we investigated the impact of the sample size of both target and reference populations.

(i) The sample size of the target population

We generated target populations with sample sizes ($n$) ranging between 1–50 individuals. For these analyses, the sample size for the reference populations was 28 French individuals and 21 Yoruba individuals from HGDP. For each target group of $n$ individuals, we applied *DATES* and observed that the inferred time of admixture was unbiased for all cases, including when only one individual was available from the target population. The estimated dates were more precise with larger sample sizes (Figure S2.3), similar to admixture disequilibrium-based dating methods like *ROLLOFF* and ALDER (2, 4, 9).

(ii) The sample size of the reference populations

To investigate the impact of the number of reference samples used in the analysis, we applied *DATES* to 10 simulated target individuals and varied the sample size of the reference panel between 1–20 individuals. *DATES* uses the reference samples to compute the allele frequency in the ancestral populations, which is in turn used to perform the simple regression to infer the ancestry proportion at each site and compute the weighted ancestry covariance across pairs of sites. Thus, this information is critical for reliable inference of estimated dates. Our simulations show that the inference can be very noisy if only a single diploid individual is used as the reference population. The estimated dates are reliable when at least 5 individuals are available from the reference groups (Figure S2.4).

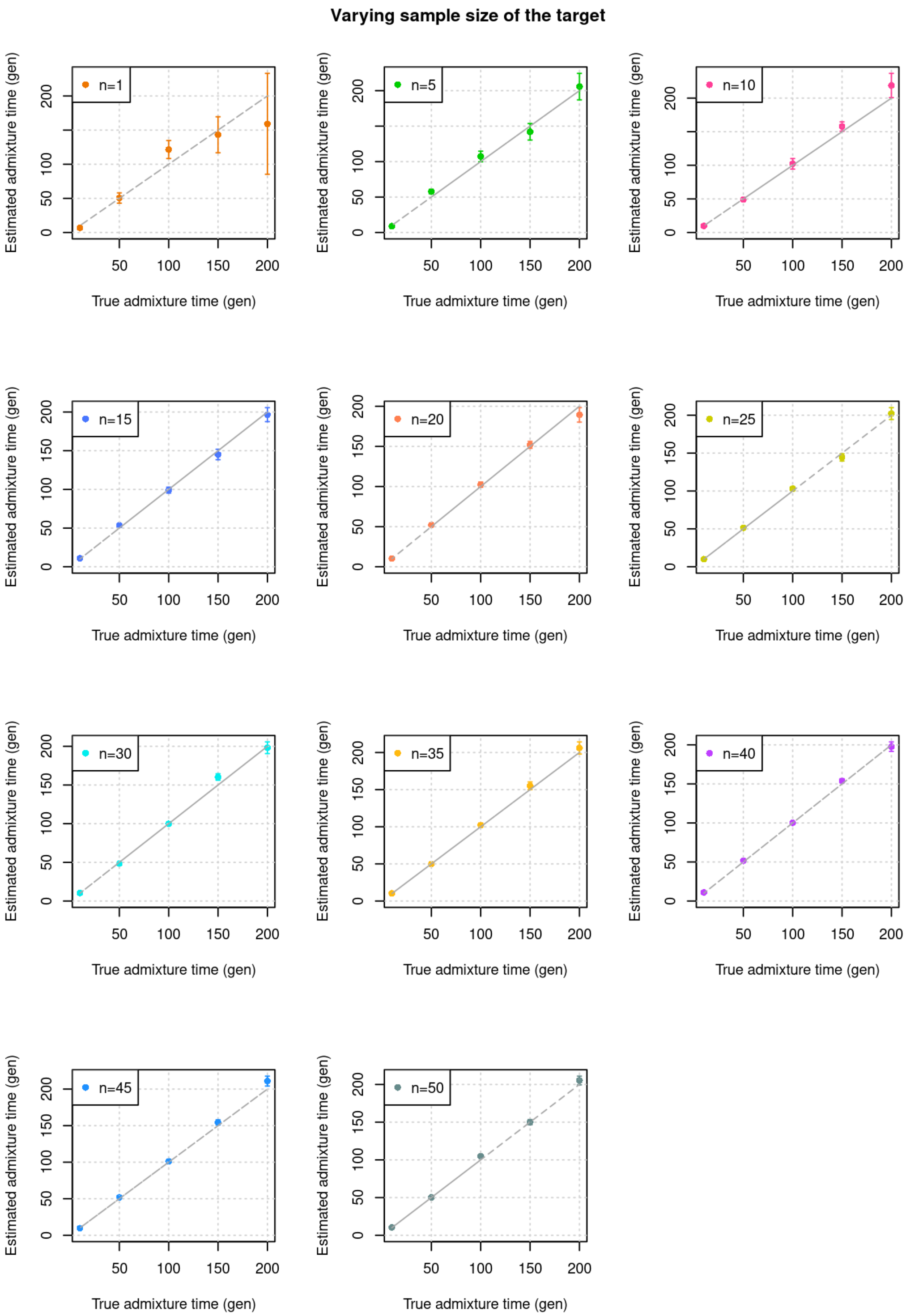


**Figure S2.3: Varying sample size of the target population**: We simulated admixed individuals with 20% European and 80% African ancestry with varying sample sizes ($n$) from 1 to 50 in increments and applied *DATES* to infer the time of admixture. The true admixture time is shown on X-axis, and the estimated time of admixture (± 1 SE) is shown on Y-axis. The different panels include results for different sample sizes of the target group.

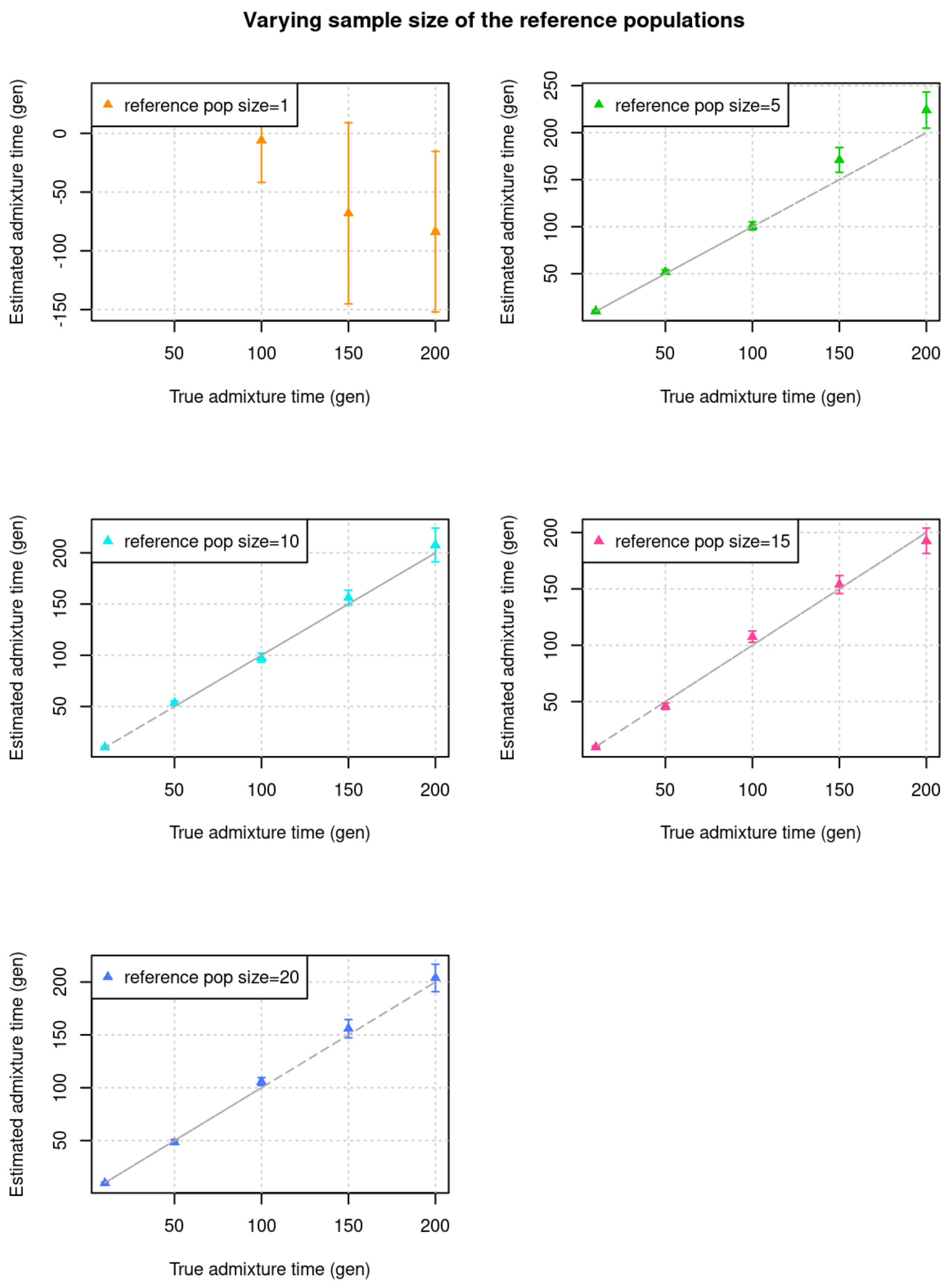


**Figure S2.4: Varying sample size of the reference populations**: We simulated 10 admixed individuals with 80% European and 20% African ancestry. We applied *DATES* with reference populations with sample sizes shown separately in each panel in the text box. The true admixture time is shown on X-axis, and the estimated time of admixture (± 1 SE) is shown on Y-axis.

#### 4. Impact of divergence between the true and reference populations

One of the advantages of *DATES* is that it does not require data from the true mixing populations for the estimation of admixture time, but reference populations that are related to the true ancestral groups work reliably for inferring the dates. We investigated the efficiency of *DATES* in estimating the admixture time with inaccurate reference populations, by using reference populations that were increasingly divergent from the true ancestral sources. Application of *DATES* in this setting showed that the inferred timing was reliable for all cases even when highly divergent groups such as Khomani San were used as the reference group instead of Yoruba in *DATES* ($F_{ST}$ ~ 0.1) (Figure S2.5).

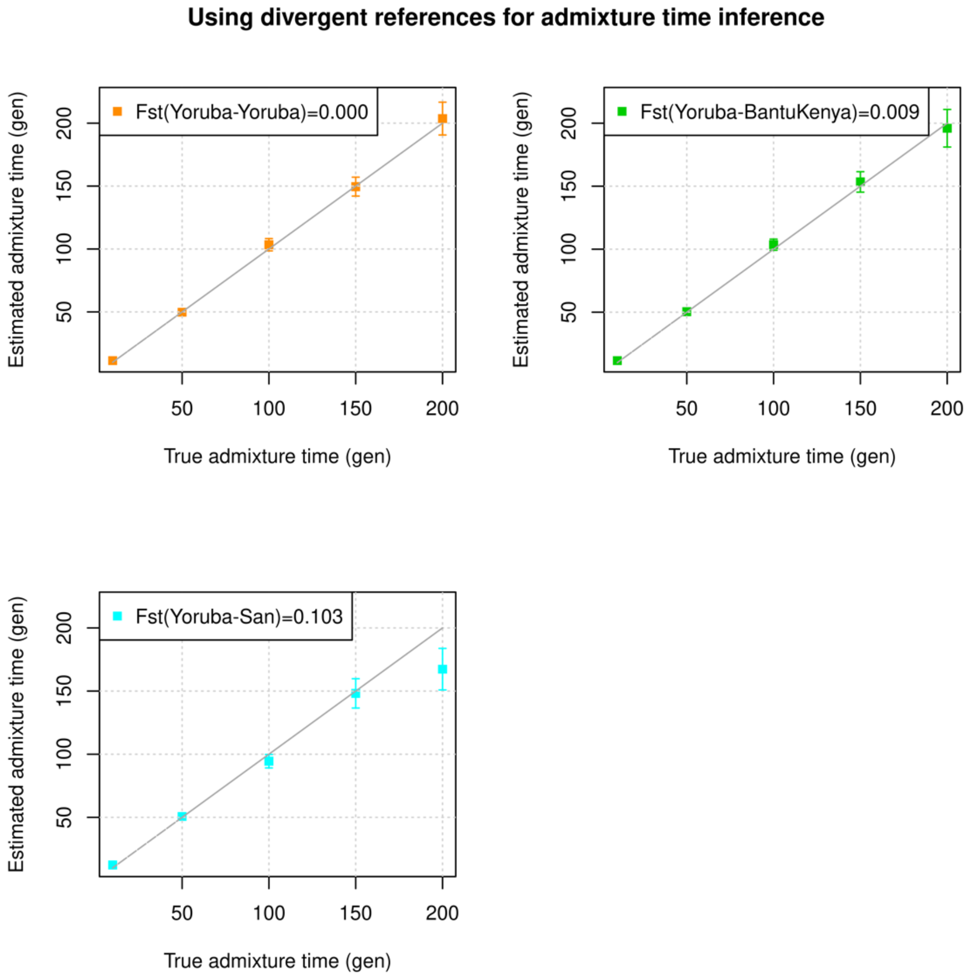


**Figure S2.5: Impact of divergence between the true and reference populations:** We simulated 10 admixed individuals with 80% European and 20% African ancestry and applied *DATES* to infer the timing of mixture using reference populations that were increasingly divergent to one of the true ancestral (French and Yoruba). For inference, we used French and a population related to Yoruba. In each panel, we show the analysis with a group that is increasingly divergent from Yoruba (the legend shows the $F_{ST}$ between Yoruba (true ancestral source) and the reference population used in *DATES*. The true admixture time is shown on X-axis, and the estimated time of admixture (± 1 SE) is shown on Y-axis.

#### 5. Impact of sparse data from the target population

One of the major goals of developing *DATES* was to apply it to ancient DNA samples to learn about past evolutionary events. However, ancient DNA specimens tend to have a low quality of data due to poor preservation and high degradation of DNA over time. Thus, ancient DNA samples often have a high proportion of missing data and limited or low coverage. Further, in most ancient DNA applications, it is a common practice to make pseudo-homozygous genotype calls using a random allele observed in the reads at each site in the genome (8) that can further lead to biases in the inference. To test the robustness of *DATES*, we simulated datasets mimicking the features of ancient DNA datasets, namely large proportions of missing data, pseudo-homozygous genotype calls, and small sample sizes.

(i) Missing genotypes
We simulated 10 admixed genomes with missing genotypes (by setting the genotype call at a site as "missing" or "unknown"; in eigenstrat format as *9*) where the proportion of missing genotypes ranged between 5–60% (in increments of 5%). We then applied *DATES* and estimated the timing of the admixture. We observed that *DATES* inferred the time of admixture accurately even for target individuals with a high missing proportion (~60%) of missing data (Figure S2.6).

(ii) Use of pseudo-homozygous genotype calls.
We simulated data for the target population with a missing genotype rate of ~10–60% (as described above (i)), with small sample sizes (*n*=1 and *n*=10) and replacing the diploid genotype calls with pseudo-homozygous calls (i.e., for every heterozygous site, we randomly assigned one of the alleles as the homozygous genotype at that site). We then applied *DATES* to infer the timing of the admixture. We observed the inference was accurate in all cases, even using a single admixed individual with only pseudo-homozygous genotypes (Figure S2.7). This shows *DATES* is reliable and applicable for the analysis of ancient DNA genomes.

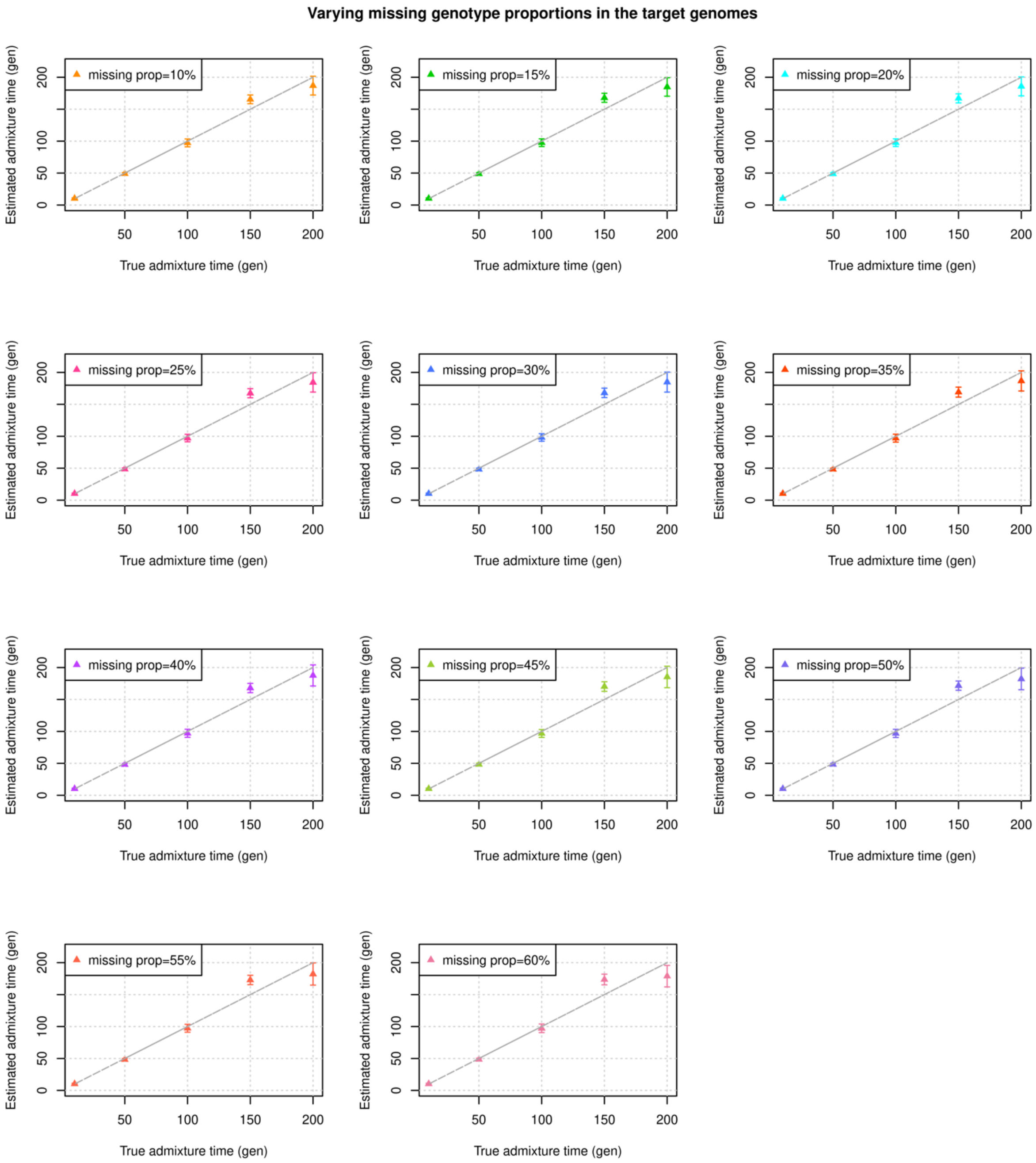


**Figure S2.6: Impact of missing genotypes in the target population**: We simulated data for 10 admixed individuals with varying proportions of missing data between ~10%-60%. Each panel shows the results for simulations with *x*% of missing genotypes (shown in the legend). The true admixture time is shown on X-axis, and the estimated time of admixture (± 1 SE) is shown on Y-axis.

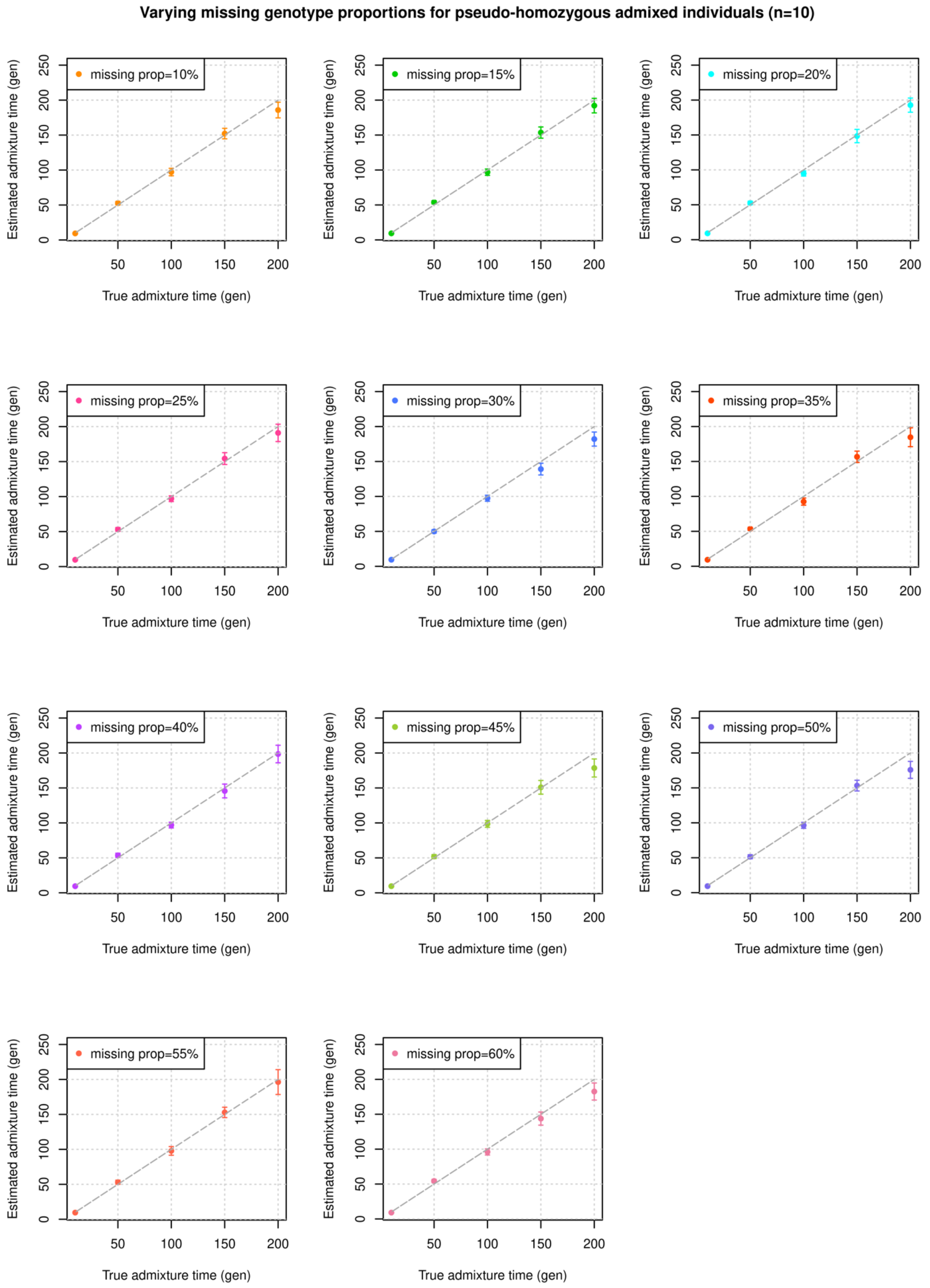
Varying missing genotype proportions for pseudo-homozygous admixed individuals (n=10)
missing prop=10%
missing prop=15%
missing prop=20%
missing prop=25%
missing prop=30%
missing prop=35%
missing prop=40%
missing prop=45%
missing prop=50%
missing prop=55%
missing prop=60%
Estimated admixture time (gen)
True admixture time (gen)
0
50
100
150
200
250

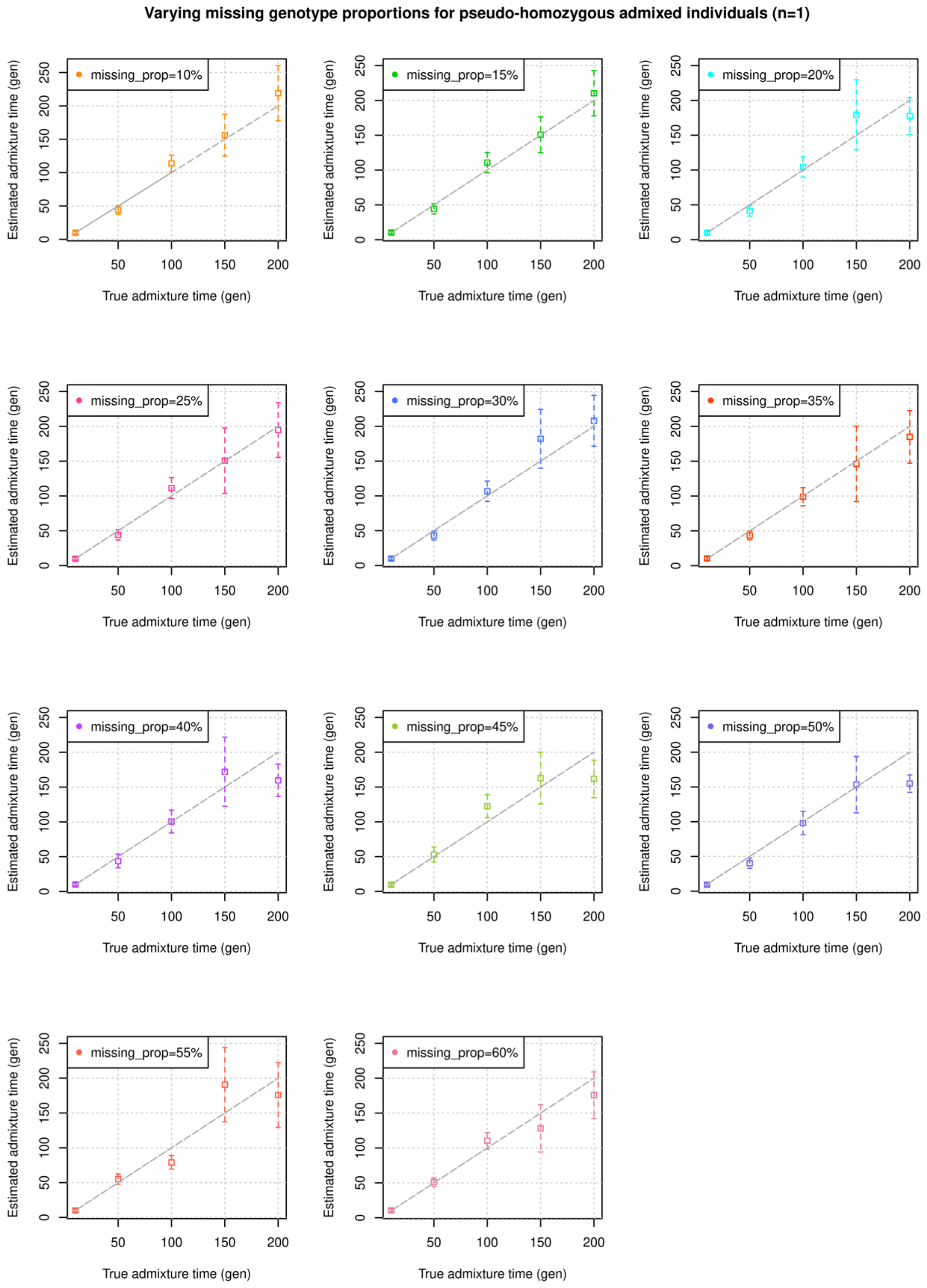


**Figure S2.7: Impact of features of ancient genomes:** We simulated data for 10 admixed individuals (Panel A) and a single admixed individual (Panel B) and matched the features of ancient DNA datasets, with large proportions of missing data (10-60%) and generating pseudo-homozygous genotypes. Each panel shows the results for simulations with *x*% of missing genotypes (shown in the legend). The true admixture time is shown on X-axis, and the estimated time of admixture (± 1 SE) is shown on Y-axis.

#### 6. Impact of divergence between the two ancestral populations

To investigate how the accuracy of *DATES* is impacted when the source populations are closely related, we simulated admixed populations with 20% Northern Europeans (CEU) and 80% ancestry from another reference population, with decreasing genetic similarity to Europeans. Specifically, the other reference population we used was either West Africans, YRI ($F_{ST} = 0.154$), East Africans, LWK ($F_{ST} = 0.142$), East Asians, CHB ($F_{ST} = 0.11$), South Asians, ITU ($F_{ST} = 0.042$), South Americans, MXL ($F_{ST} = 0.037$) or Southern Europeans, TSI ($F_{ST} = 0.004$). To minimize any issues with overfitting, we used the following reference populations for the inference: one reference as French and the other reference as either Yoruba, Bantu Kenya, Tujia, Sindhi, Maya, or Italian respectively. Application of *DATES* showed that for most cases the inferred timing was accurate, except when the ancestral groups were very closely related such as CEU and TSI ($F_{ST} = 0.004$). (Figure S2.8). This is likely due to the fact that the allele frequencies within continental groups such as Northern and South Europeans are correlated and so the method is unable to differentiate between background correlation and admixture correlation thus leading to noisy inference.

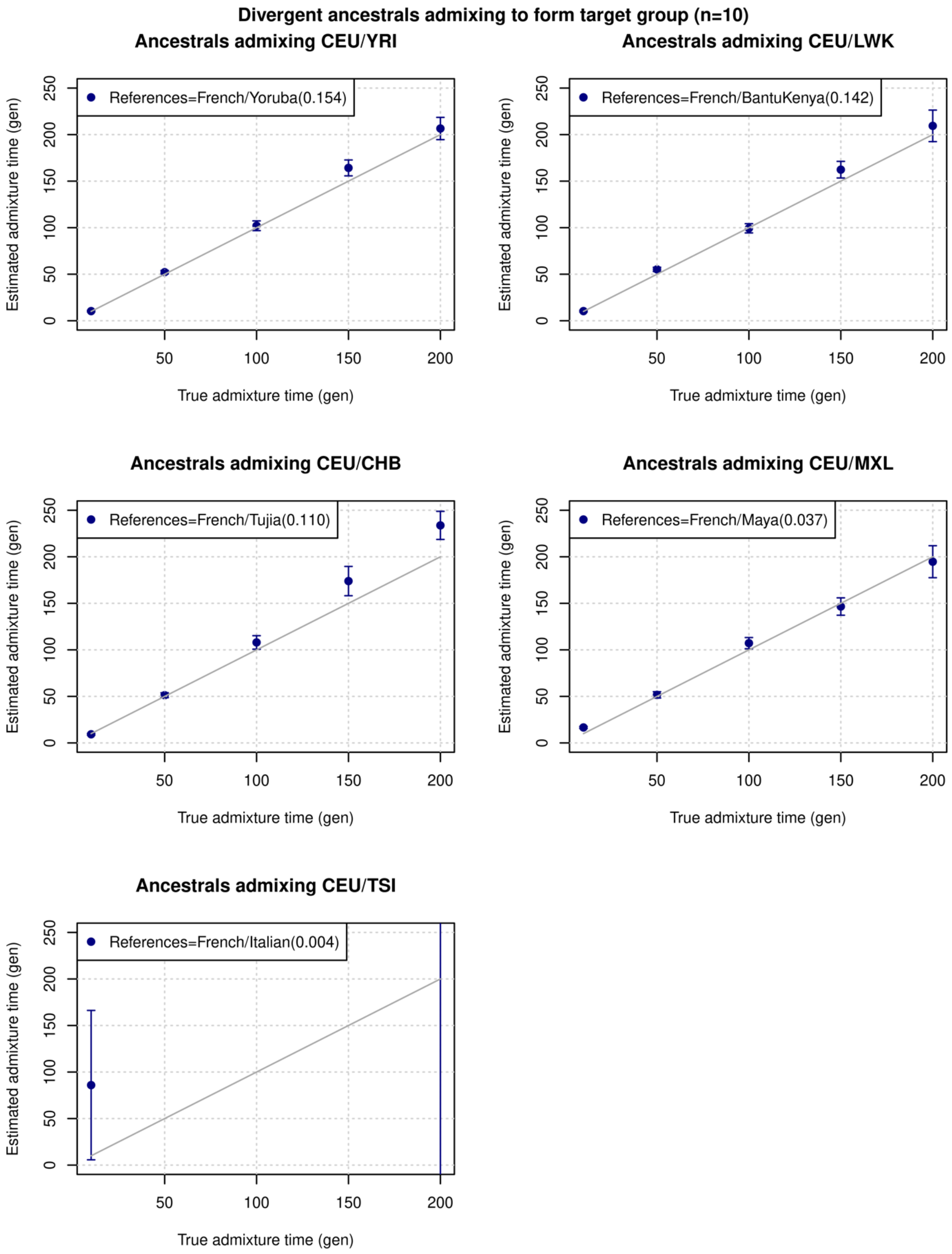
Divergent ancestrals admixing to form target group (n=10)
Ancestrals admixing CEU/YRI
References=French/Yoruba(0.154)
Ancestrals admixing CEU/LWK
References=French/BantuKenya(0.142)
Ancestrals admixing CEU/CHB
References=French/Tujia(0.110)
Ancestrals admixing CEU/MXL
References=French/Maya(0.037)
Ancestrals admixing CEU/TSI
References=French/Italian(0.004)
Estimated admixture time (gen)
True admixture time (gen)
0
50
100
150
200
250

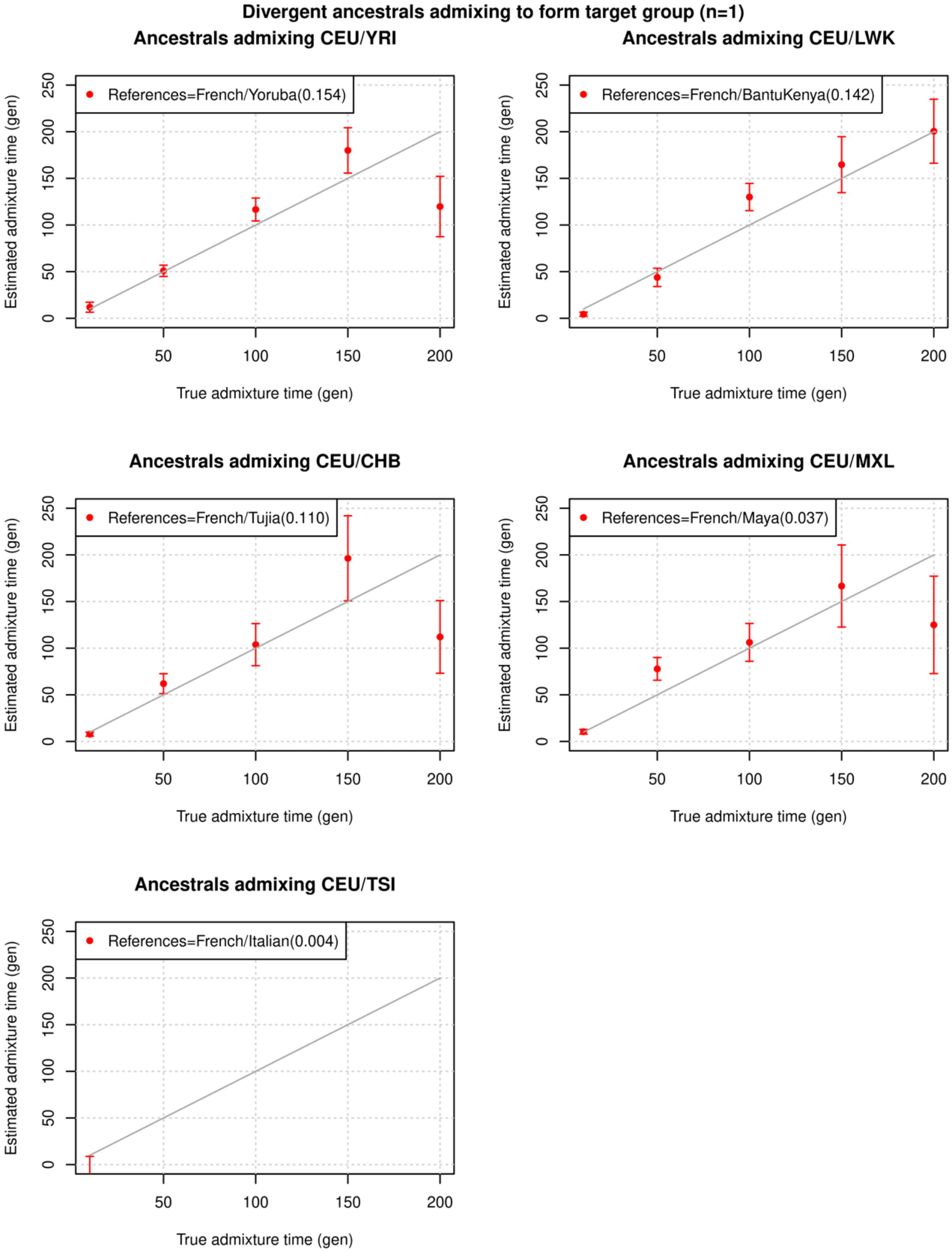


**Figure S2.8: Impact of divergence between the two ancestral populations:** We simulated data for 10 individuals (Panel A) and one individual (Panel B) with 20% European (CEU) and 80% ancestry from a range of populations (shown in each sub-panel). We used closely related reference populations (shown in the legend). The true admixture time is shown on X-axis, and the estimated time of admixture (± 1 SE) is shown on Y-axis.

#### 7. Using admixed populations as one of the reference groups

Previous methods such as ALDER have shown that one can leverage the admixed population as one of the reference populations for the inference (2). This is very desirable as in some cases, it is difficult to obtain reference data for more than one ancestral source population. To test if this feature provides reliable estimates of the timing of mixture using the *DATES*, we simulated two non-overlapping sets of admixed individuals– one set was used as the target and the other as one of the reference populations with either French (minor parent with ~20%) or Yoruba (major parent with ~80%) as the second reference. Figure S2.9 shows the inferred dates were accurate in all cases, even when the minor parental population was used as one of the reference populations. The ability to use the admixed population as one of the references opens the possibility of dating with only one reference as in ALDER.

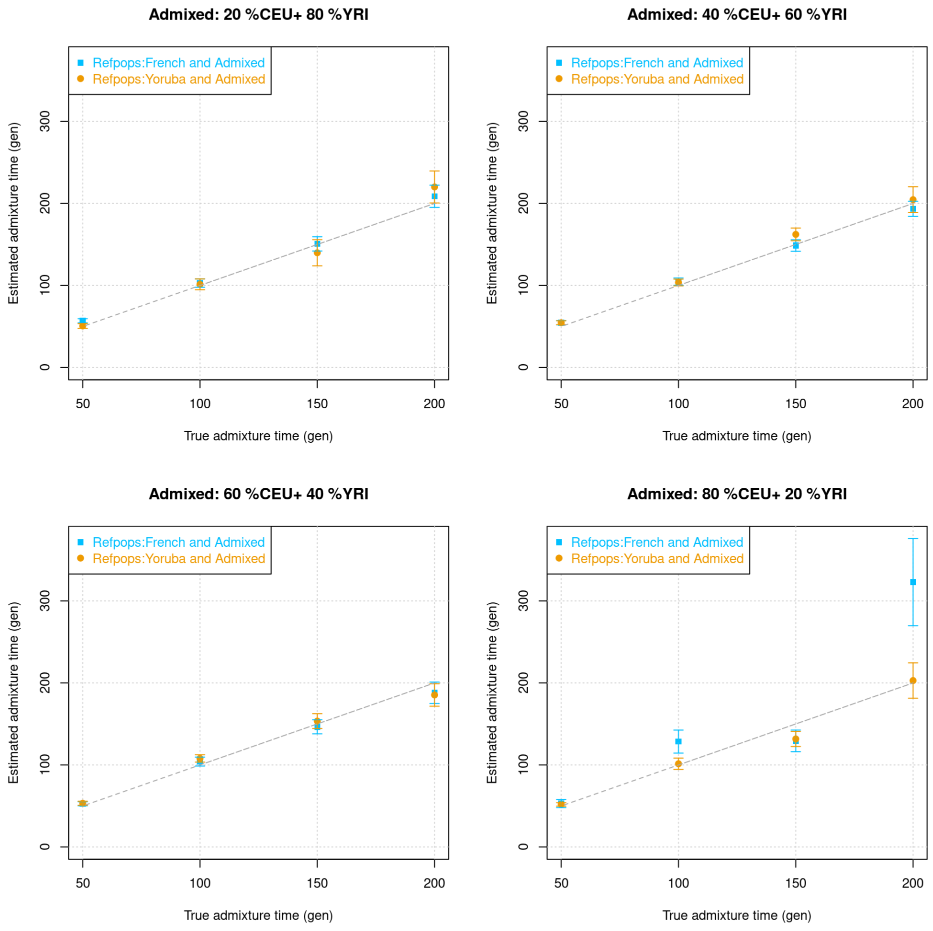


**Figure S2.9: Using admixed reference populations in *DATES*:** We simulated data for 20 individuals using two separate reference datasets, with varying proportions of European ($\alpha = 0.2, 0.4, 0.6, 0.8$) and African ancestry ($1 - \alpha$) shown above in four panels. Out of these 20 individuals, we used 10 individuals as the target and 10 individuals as one of the reference populations. For each admixed group, we ran *DATES* with French and simulated admixed individuals as the reference populations (shown in blue points), and Yoruba and simulated admixed individuals (shown as orange points). The true admixture time is shown on X-axis, and the estimated time of admixture (± 1 SE) is shown on Y-axis.

#### 8. Multiple pulses of admixture

*DATES* models admixed individuals as a mix of two source populations. However, in real data, the target could have ancestry from more than two source populations. To test the impact of this scenario, we generated a target population that has ancestry from three groups (*PopA*, *PopB*, *PopC*) that mixed at two distinct times ($t_1$ and $t_2$ generations ago) (Figure S2.10). Specifically, we generated data for three sets of admixed populations each with 10 individuals, where *PopA*, *PopB,* and *PopC* differ across runs. For each simulation, the older pulse of admixture occurred $t_2$ (=30, 60, 100) generations ago and *PopA* and *PopB* mixed with $\alpha_1/\alpha_2$ ancestry respectively. This mixture was followed by additional gene flow from *PopC* that contributed $\alpha_3$ ancestry at $t_1$ (=10) generation ago. We used CEU, YRI, and CHB as *PopA*, *PopB,* and *PopC* and varied the order of the three ancestral populations to generate multiple sets of simulated individuals (see Figure S2.11). We estimated admixture time for all three sets of simulated individuals using French, Tujia, and Yoruba as reference populations.

(i) Equal proportions of ancestry from three sources.

In this setup, we simulated admixed individuals with ancestry from *PopA*, *PopB,* and *PopC* with $\alpha_1 = 50\%$, $\alpha_2 = 50\%$, $\alpha_3 = 33\%$ thus the effective ancestry proportion in the admixed population would be 33% from each ancestral group. We varied the order of the three ancestrals and applied *DATES* to infer the timing of the mixture. Our results showed that when the references populations used for the inference correspond to the true admixing sources, in most cases, we reliably estimate the timing of both the pulses of admixture (Figure S2.11).

(ii) Unequal proportions of ancestry from three ancestrals.

In most real-world scenarios, the ancestry proportions of the admixing groups are unlikely to be exactly the same or similar. Thus, we generated data for groups with unequal proportions of ancestry from *PopA*, *PopB,* and *PopC* by setting $\alpha_1 = 20\%$, $\alpha_2 = 80\%$, and $\alpha_3 = 20\%$ or $80\%$. We varied the order of the three ancestral groups and applied *DATES* to infer the timing of the mixture. We observed that we recovered the timing of the recent pulse of admixture in most cases (Figure S2.12). In some cases, there was confounding in the timing of the recent event, when the % of ancestry from *PopC* was low (20%). In Note S2.9, we explore how choosing ancestral populations that are more aligned with the model of admixture (that can be reliably inferred using other methods like *qpAdm*) can alleviate this bias (Table S2.1).

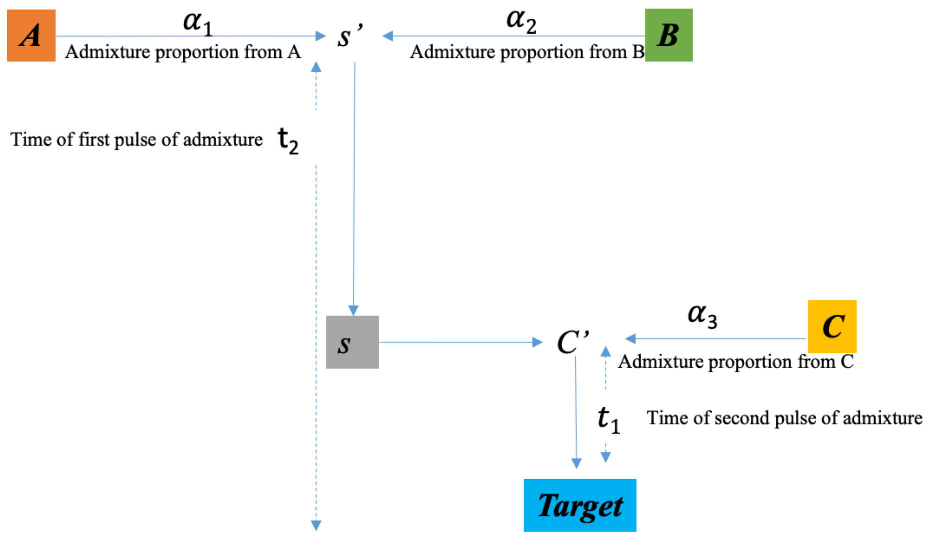


**Figure S2.10: Model for multiple pulses of admixture.** The admixed population (*Target)* derives ancestry from three populations, from the two gene flow events that occurred $t_2$ generations ago (older pulse) between *PopA* and *PopB* with $\alpha_1$ / $\alpha_2$ ancestries respectively resulting in an intermediate group *S*, which then mixes with *PopC* with $\alpha_3$ ancestry *at* $t_1$ generations ago (younger pulse).

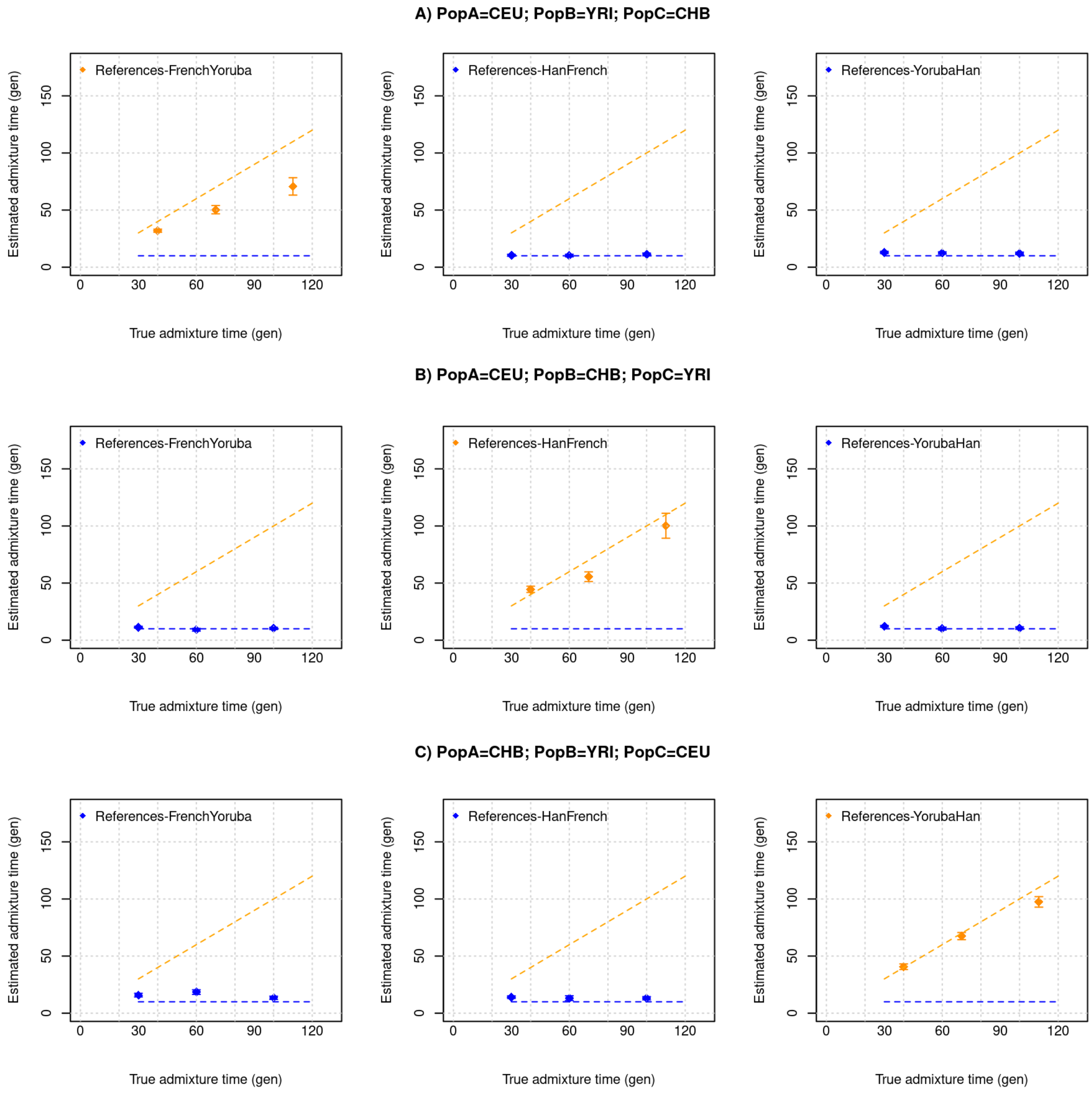


**Figure S2.11: Multiple pulses of admixture with equal proportions of ancestry from sources.** We generated a target population that has ancestry from three groups (*PopA*, *PopB*, *PopC*) with ancestry proportions of 33%, 33% and 33% respectively that mixed at two distinct times ($t_1$ = 30,60,100 and $t_2$ = 10 generations ago). We used CEU, YRI, and CHB as *PopA*, *PopB,* and *PopC* and varied the order of the three ancestrals, and applied *DATES* with pairs of populations as the reference to infer the timing of the mixture. We show the expected dates ($t_1$ or $t_2$ depending on the references used), the orange dashed line corresponds to the older pulse and the blue dashed line corresponds to the younger pulse of admixture. The blue points correspond to *DATES* estimates using Pop*A* and Pop*C* or Pop*B* and Pop*C* as references. The orange points correspond to *DATES* estimates using Pop*A* and Pop*B* as references. Panel (A) shows the admixture scenario with *PopA* = CEU, *PopB* = YRI and *PopC* = CHB, Panel (B) shows the admixture scenario with *PopA* = CEU, *PopB* = CHB and *PopC* = YRI, and C) shows the admixture scenario with *PopA* = CHB, *PopB* = YRI and *PopC* = CEU.

**Panel (I) PopA α1 = 20%; PopB α2 = 80%; PopC α3 = 80%**

**A) PopA=CEU; PopB=YRI; PopC=CHB**

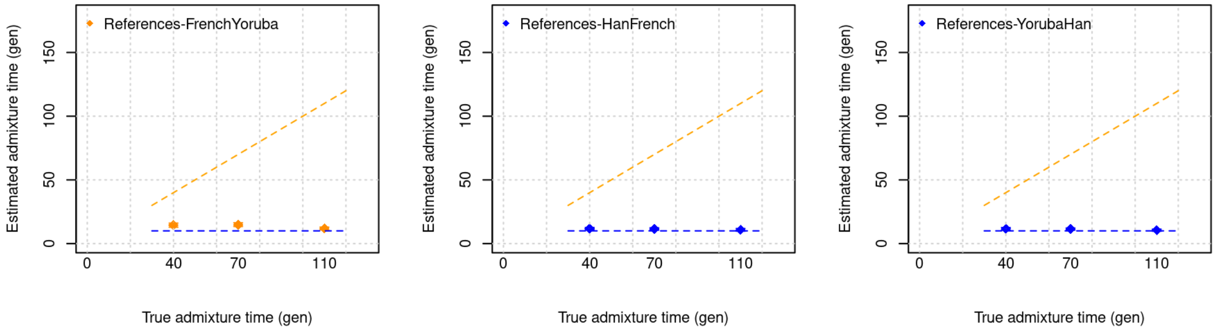


**B) PopA=CEU; PopB=CHB; PopC=YRI**

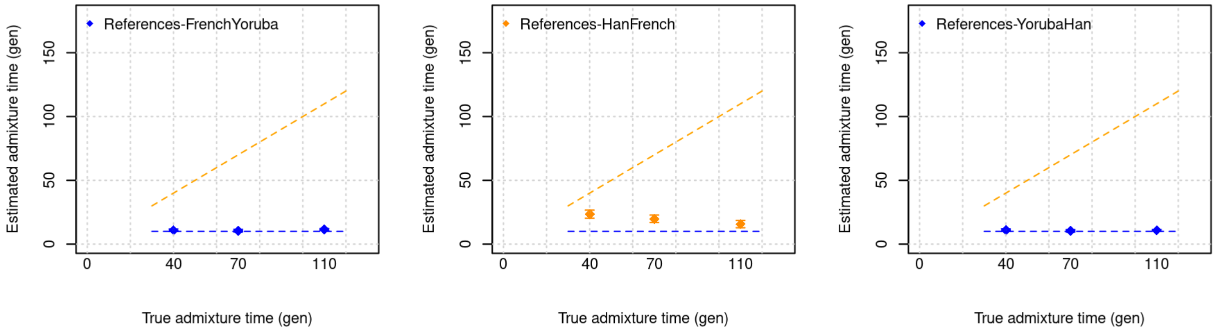


**C) PopA=CHB; PopB=YRI; PopC=CEU**

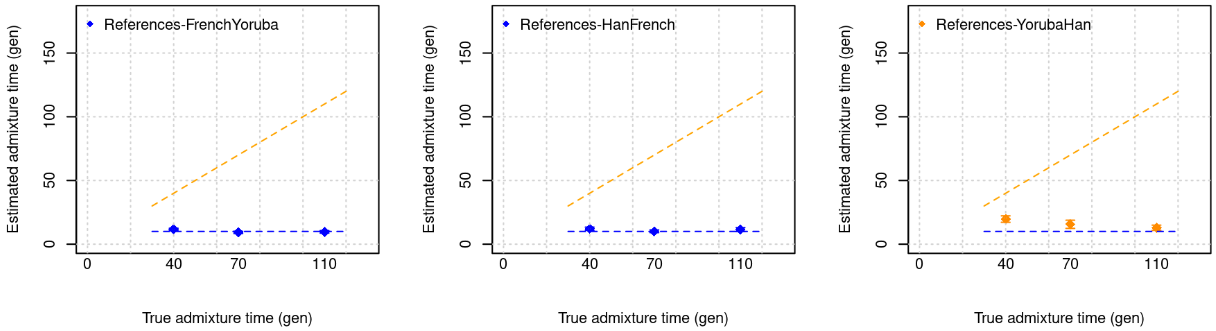

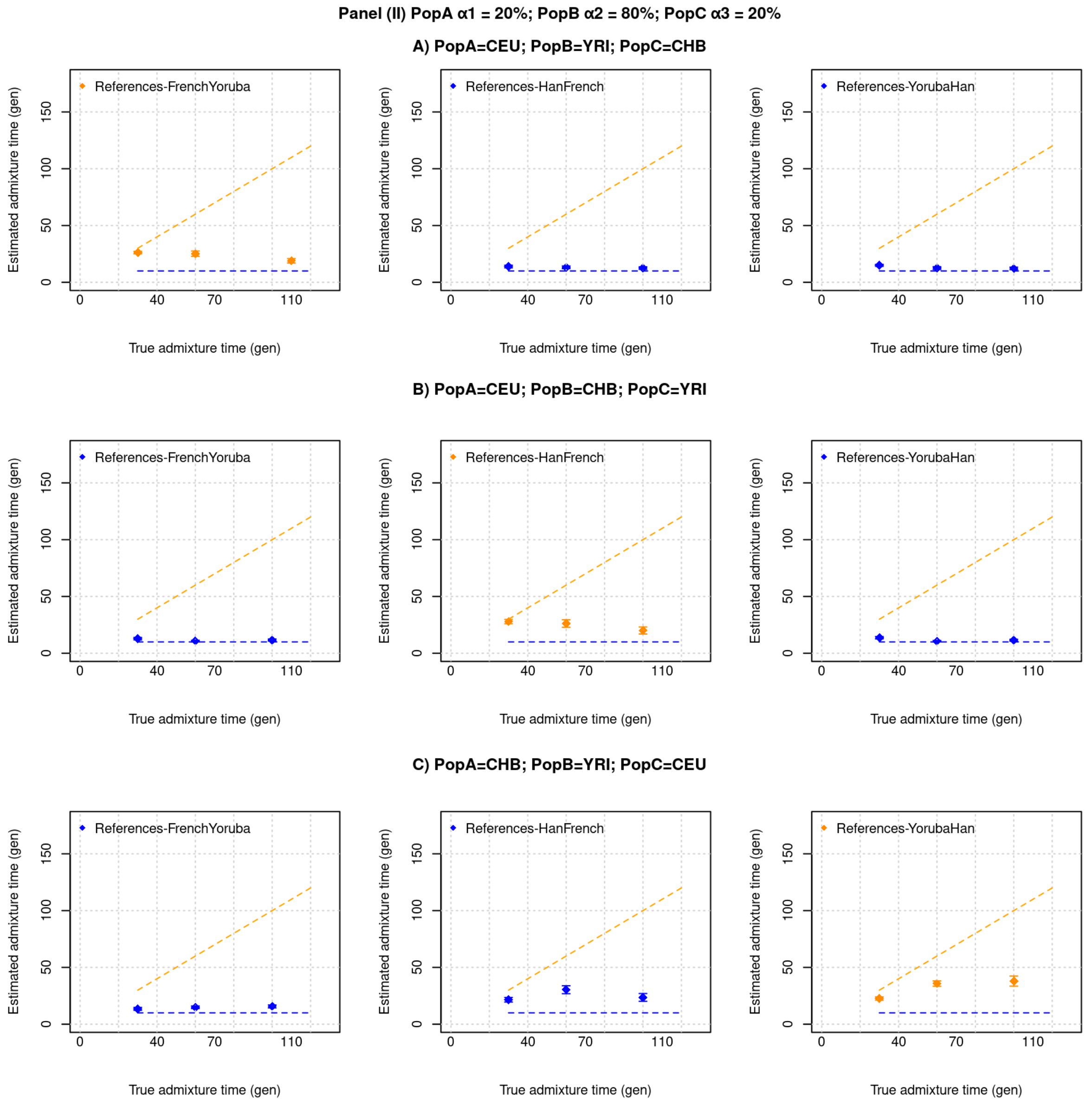


**Figure S2.12: Two pulses of admixture with unequal proportions of ancestry from reference populations.** We generated a target population that has ancestry from three groups with variation in ancestry from three sources. Panel (I):*PopA*, *PopB*, *PopC* with ancestry proportion of 4%, 16% and 80% respectively that mixed at two distinct times ($t_1$ = 30,60,100 and $t_2$ = 10 generations ago). Panel (II):*PopA*, *PopB*, *PopC* with ancestry proportion of 16%, 64% and 20% respectively that mixed at two distinct times ($t_1$ = 30,60,100 and $t_2$ = 10 generations ago). We used CEU, YRI, and CHB as *PopA*, *PopB,* and *PopC* and varied the order of the three ancestral populations, and applied *DATES* with pairs of populations as the reference to infer the timing of the mixture. Figures shows the true time of admixture on the X-axis and inferred time on Y-axis, the orange dashed line corresponds to the older pulse and the blue dashed line corresponds to the younger pulse of admixture. The blue points correspond to *DATES* estimates using Pop*A* and Pop*C* or Pop*B* and Pop*C* as references. The orange points correspond to *DATES* estimates using Pop*A* and Pop*B* as references. Panel (A) shows the admixture scenario with *PopA* = CEU, *PopB* = YRI and *PopC* = CHB, Panel (B) shows the admixture scenario with *PopA* = CEU, *PopB* = CHB and *PopC* = YRI, and C) shows the admixture scenario with *PopA* = CHB, *PopB* = YRI and *PopC* = CEU.

#### 9. Choice of references in a multi-pulse admixture

Using the admixture scenario described above, we examined how the choice of reference populations impacts the inferred dates of admixture. From Note S2.8, we find that using PopC as one of the references gives more reliable dates. In most real-world scenarios, the ordering of gene flow events is known. For instance, for present-day Europeans, it's known that Steppe pastoralist gene flow occurs after the gene flow between Anatolian farmers and local European hunter-gatherers. Thus, we fixed one reference group as *PopC* and explored how the choice of the second reference population impacts the recovery of the recent gene flow event. To ensure that our observed dates are not biased, we ran 10 replicates for each run and report the average of the 10 runs below. Specifically, for the target populations with ancestries from *PopA*, *PopB,* and *PopC*, we ran *DATES* with the following reference populations:

(i) Ref1: *PopC* and Ref2: *PopA*
(ii) Ref1: *PopC* and Ref2: *PopB*
(iii) Ref1: *PopC* and Ref1: admixed individuals with *PopA* and *PopB* ancestry (30% *PopA* / 70% *PopB*) ancestry
(iv) Ref1: *PopC* and Ref2: pooled samples from *PopA* and *PopB*

We observed that in all four cases using *PopC* (admixing source in the most recent pulse) as one of the sources allows us to recover the recent pulse of admixture. When the other reference population is either the pooled set of samples of *PopA* and *PopB* (admixing sources in the older pulse) or admixed individuals with ancestry from *PopA* and *PopB*, we reliably infer the younger pulse of admixture in all cases (Supplementary Table S2.1). In other cases, we observe some confounding in the inferred dates with the estimated dates falling intermediate of two dates.

**Table S2.1: Impact of reference populations in two-way admixed groups.**

| **Model: We generated target populations with two pulses of gene flow where *PopA* and *PopB* mixed at time $t_1$ generations ago with ancestry proportion of $\alpha_1$ and $\alpha_2$, followed by gene flow from *PopC* at $t_2$ generations ago with ancestry proportion of $\alpha_3$** | | | | | | | |
|---|---|---|---|---|---|---|---|
| **Target** | **$t_2/t_1$** | **Ref1** | **Ref 2** | **$\alpha_1$ =20%, $\alpha_2$ =80%, $\alpha_3$ =80%** | **$\alpha_1$ =50%, $\alpha_2$ =50%, $\alpha_3$ =50%** | **$\alpha_1$ =20%, $\alpha_2$ =80%, $\alpha_3$ =20%** | **$\alpha_1$ =20%, $\alpha_2$ =80%, $\alpha_3$ =10%** |
| **A. Using reference populations *PopA* and *PopC*** | | | | | | | |
| *PopA = CEU PopB = YRI PopC = CHB* | 100/10 | Han | French | 10.3 | 10.7 | 12.0 | 12.6 |
| *PopA = CEU PopB = CHB PopC = YRI* | 100/10 | Yoruba | French | 10.3 | 9.7 | 10.2 | 10.6 |
| *PopA = CHB PopB = YRI PopC = CEU* | 100/10 | French | Han | 10.8 | 11.6 | 20.7 | 44 |
| *PopA = CEU PopB = YRI PopC = CHB* | 60/10 | Han | French | 11.3 | 10.7 | 11.8 | 14.4 |
| *PopA = CEU PopB = CHB PopC = YRI* | 60/10 | Yoruba | French | 10.4 | 10.8 | 10.3 | 11.2 |
| *PopA = CHB PopB = YRI* | 60/10 | French | Han | 11.2 | 12.6 | 19.9 | 40.7 |

| | | | | | | | |
|---|---|---|---|---|---|---|---|
| *PopC = CEU* | | | | | | | |
| **B. Using reference populations *PopB* and *PopC*** | | | | | | | |
| *PopA = CEU*<br>*PopB = YRI*<br>*PopC = CHB* | 100/10 | Han | Yoruba | 10.2 | 11.5 | 11.3 | 11.9 |
| *PopA = CEU*<br>*PopB = CHB*<br>*PopC = YRI* | 100/10 | Yoruba | Han | 10.1 | 9.7 | 10.3 | 10.5 |
| *PopA = CHB*<br>*PopB = YRI*<br>*PopC = CEU* | 100/10 | French | Yoruba | 10.1 | 12.8 | 12.2 | 14 |
| *PopA = CEU*<br>*PopB = YRI*<br>*PopC = CHB* | 60/10 | Han | Yoruba | 11.0 | 12.6 | 12.2 | 14.6 |
| *PopA = CEU*<br>*PopB = CHB*<br>*PopC = YRI* | 60/10 | Yoruba | Han | 10.3 | 11.1 | 10.6 | 11.4 |
| *PopA = CHB*<br>*PopB = YRI*<br>*PopC = CEU* | 60/10 | French | Yoruba | 10.4 | 13.4 | 12.4 | 18 |
| **C. Using reference populations *PopC* and "admixed" individuals with ancestry from *PopA* and *PopB* *(30% PopA / 70% PopB)* ancestry** | | | | | | | |
| *PopA = CEU*<br>*PopB = YRI*<br>*PopC = CHB* | 100/10 | Han | Admixed<br>(30% CEU/<br>70% YRI) | 10.1 | 10.9 | 10.8 | 10.5 |
| *PopA = CEU*<br>*PopB = CHB*<br>*PopC = YRI* | 100/10 | Yoruba | Admixed<br>(30% CEU/<br>70% CHB) | 10.1 | 9.5 | 10.0 | 10 |
| *PopA = CHB*<br>*PopB = YRI*<br>*PopC = CEU* | 100/10 | French | Admixed<br>(30% CHB/<br>70% YRI) | 10.0 | 11.2 | 10.9 | 10.8 |
| *PopA = CEU*<br>*PopB = YRI*<br>*PopC = CHB* | 60/10 | Han | Admixed<br>(30% CEU/<br>70% YRI) | 11 | 11.3 | 10.9 | 11.8 |
| *PopA = CEU*<br>*PopB = CHB*<br>*PopC = YRI* | 60/10 | Yoruba | Admixed<br>(30% CEU/<br>70% CHB) | 10.3 | 10.8 | 10.3 | 10.5 |
| *PopA = CHB*<br>*PopB = YRI*<br>*PopC = CEU* | 60/10 | French | Admixed<br>(30% CHB/<br>70% YRI) | 10.2 | 11.3 | 10.6 | 12.1 |
| **D. Using reference populations *PopC* and pooled individuals of *PopA* and *PopB* ancestry** | | | | | | | |
| *PopA = CEU*<br>*PopB = YRI*<br>*PopC = CHB* | 100/10 | Han | French + Yoruba | 10.1 | 10.99 | 10.4 | 9.5 |
| *PopA = CEU*<br>*PopB = CHB*<br>*PopC = YRI* | 100/10 | Yoruba | French + Han | 10.2 | 9.5 | 10.03 | 9.9 |
| *PopA = CHB*<br>*PopB = YRI*<br>*PopC = CEU* | 100/10 | French | Han + Yoruba | 9.9 | 10.4 | 10.6 | 10 |
| *PopA = CEU*<br>*PopB = YRI*<br>*PopC = CHB* | 60/10 | Han | French + Yoruba | 10.9 | 10.8 | 10.3 | 10.3 |
| *PopA = CEU*<br>*PopB = CHB*<br>*PopC = YRI* | 60/10 | Yoruba | French + Han | 10.3 | 10.7 | 10.1 | 10.5 |
| *PopA = CHB*<br>*PopB = YRI*<br>*PopC = CEU* | 60/10 | French | Han + Yoruba | 10.0 | 10.3 | 9.8 | 11 |

Note: the estimated dates are shown per scenario are averages of 10 simulations.

#### B. Coalescent simulations

To evaluate the performance of *DATES* under more complex demographic models involving gradual gene flow over a long period of time and founder events, we performed simulations using the coalescent simulator, *macs* (10). For all the simulations described below, we generated data for three populations (*PopA*, *PopB,* and *PopC*) for a region of 100Mb with 22 replicates. The effective population size ($N_e$) of all three populations was assumed to be 12,500 with the mutation rate and recombination rate was assumed as 1.2 x $10^{-8}$ and 1 x $10^{-8}$ per base pair per generation respectively (11, 12). The divergence time between population $A$ and $B$ was assumed to be 1800 generations, which translates to an estimated $F_{ST}$ (*PopA*, *PopB*) of 0.067. *PopC* was formed by admixture between *PopA* to *PopB* that occurred either continuously over a period of $\lambda$ generations or instantaneously at time $t$. We combined two haploid chromosomes at random to generate one diploid chromosome.

##### (i) Impact of continuous gene flow

To model continuous gene flow, we simulated a gradual mixture in *PopC* from *PopA*/ *PopB* over a period of $\lambda\ (= 5 - 60)$ generations, leading to 20%/ 80% *PopA*/ *PopB* ancestry. Applying *DATES* to *PopC* with *Pop A* and *PopB* as the reference populations showed that the inferred time was intermediate between the start and end of the period of gene flow (Table S2.2). This is similar to the results of other admixture dating methods like Globetrotter, ALDER, and ROLLOFF (1, 2, 13) and can be explained by the fact that continuous admixture leads to mixtures of exponential curves, and resolving the timing in such case can be challenging due to the well-known difficulty of fitting a sum of exponentials to data with even a small amount of noise (14).

*macs command line for continuous admixture with $\lambda = 5$ generations:*
*macs* 120 1e8 -t 6e-4 -r 5e-4 -I 3 50 20 50 -em 0.0002 2 1 2000 -em 0.0003 2 1 0 -ej 0.00032 2 3 -ej 0.036 1 3

**Table S2.2: Impact of continuous gene flow.** The table shows true and inferred times of admixture in *PopC* using *PopA* and *PopB* used as the reference populations.

| The true period of continuous admixture, $\lambda$ generations | Inferred time of admixture (mean ± 1 SE) is shown on Y-axis. generations) |
|---|---|
| 10-15 | 15 ± 1 |
| 20-30 | 23 ± 2 |
| 40-60 | 53 ± 3 |
| 40-100 | 64 ± 4 |

#### (ii) Impact of founder event/ bottleneck post admixture

Many human populations have a history of founder events in their recent evolutionary past (3). A founder event generates long-range LD in the target population, which could in principle be spuriously inferred as admixture-related ancestry covariance, confounding the dates of admixture. To explore the effect of this scenario, we simulated *PopC* that has ancestry from *PopA* and *PopB* due to gene flow that occurred $T_A$ generations ago. Following admixture, *PopC* experienced a bottleneck that occurred $T_B$ (=10, 80 or 100) generations ago where the effective population size decreased from 12,500 to $N_B$ (=10, 100, 500, and 1000) for a duration of $D_B$ generations (=1, 5 or 10). After $T_B$, the population recovered to the original population size of 12,500 (Figure S2.13(a)). We applied *DATES* to *PopC* using *PopA* and *PopB* as reference populations and found that the estimated dates of admixture were accurate when the bottleneck was not extreme (Figure S2.13(b)). In the case of strong bottlenecks where $N_B$ is less than 100, we observed a downward bias in the estimated admixture time.

*Macs command line:*

*macs* 120 1e8 -t 6e-4 -r 5e-4 -I 3 50 20 50 -em 0.002 2 1 10000 -em 0.00202 2 1 0 -en 0.0002 2 0.0002 -en 0.0003 2 1 -ej 0.00204 2 3 -ej 0.036 1 3

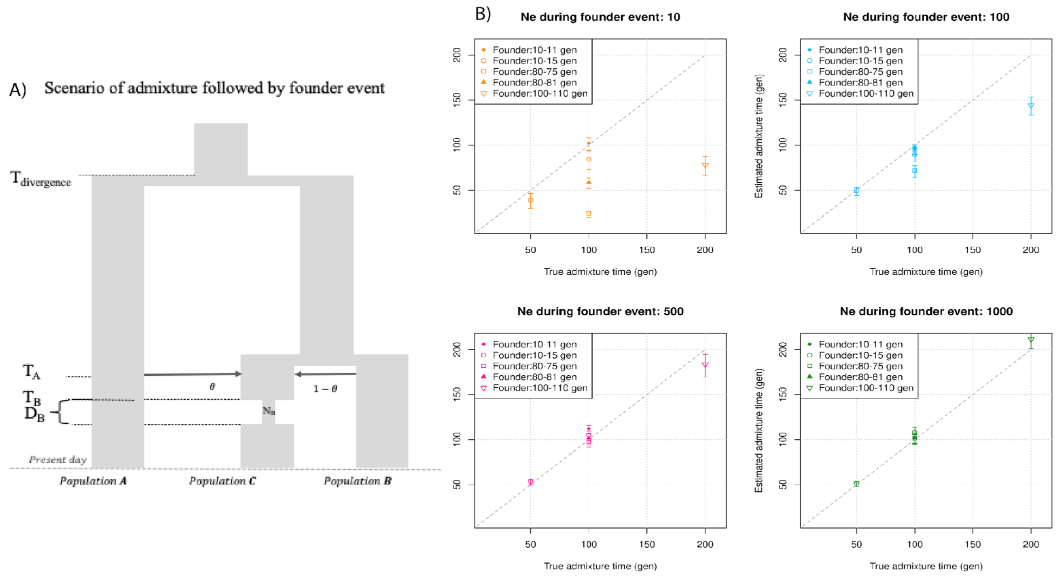


**Figure S2.13: Impact of founder events on inferred dates of admixture:** A) Schematic for demographic scenario shows that *PopC* was formed through admixture between *PopA* and *PopB* at time $T_A$. Following admixture, *PopC* experienced a severe bottleneck that occurred $T_B$ generations ago where the effective population size decreased from 12,500 to $N_B$ for a duration of $D_B$ generations. After $T_B$, the population recovered to the original population size. B) We simulated data for 10 individuals with admixture occurring at 50, 100, and 200 generations with bottleneck post admixture for a period of $D_B$ =1, 5, or 10 generations (shown in the legend) with the effective population size during bottleneck as $N_B$ =10, 100, 500 or 1000 individuals (shown as four panels). The true admixture time is shown on X-axis, and the estimated time of admixture (± 1 SE) is shown on Y-axis.

Another scenario we considered is when a population undergoes a severe bottleneck but does not recover (i.e., maintains a historically low population size to present). To test *DATES* for this demographic scenario, we simulated an admixed population that experienced a bottleneck post-admixture that occurred 100 generations ago. The effective population size was then reduced from 12,500 to $N_B$ (=100–4000). This population maintained a small size until the present. Using *DATES* with the target as *PopC* and *PopA* and *PopB* as reference populations, we observed the inferred admixture times can be biased when $N_B < 1000$; there is no bias when the effective population size is larger (Figure S2.14, Table S2.3).

*macs command line:*
macs 120 1e8 -t 6e-4 -r 5e-4 -I 3 50 20 50 -em 0.002 2 1 10000 -em 0.00202 2 1 0 -en 0 2 0.04 -en 0.00198 2 1 -ej 0.00204 2 3 -ej 0.036 1 3

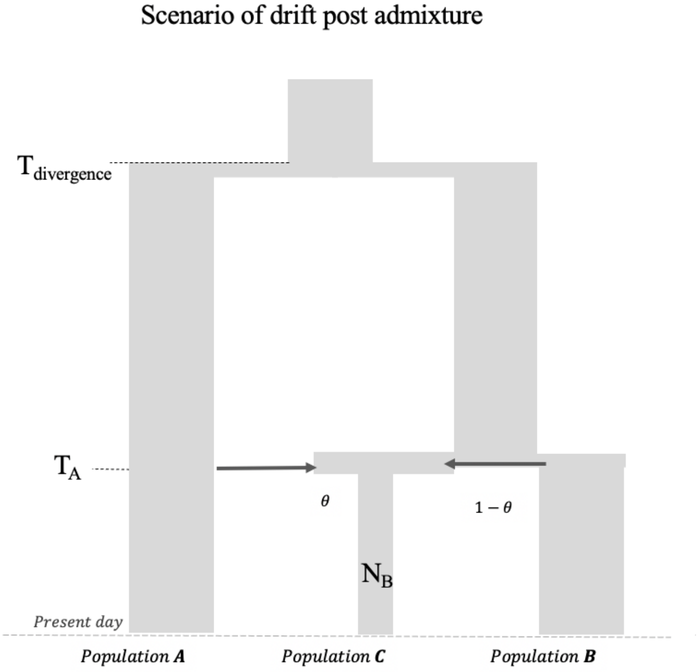


**Figure S2.14: Impact of founder event with no recovery in admixed population:** Schematic for the demographic history of the admixed group *PopC* that has ancestry from *PopA* and *PopB* followed by a severe bottleneck post-admixture without recovery to present (i.e., maintenance of historically low population size to present $N_B$). The $F_{ST}$ (*PopA*, *PopC)* is 0.202 and $F_{ST}$ (*PopB*, *PopC)* is 0.168.

**Table S2.3: Admixture time estimates from *DATES* for populations with extreme bottlenecks with a historically low population size that does not recover until the present.**

| Admixture | Ne before admixture | Ne post admixture | Inferred time of admixture |
|---|---|---|---|
| 100 | 12500 | 4000 | 96 ± 5 |
| | | 3500 | 96 ± 5 |
| | | 3000 | 88 ± 4 |
| | | 2500 | 99 ± 5 |
| | | 2000 | 92 ± 6 |
| | | 1500 | 78 ± 6 |
| | | 1000 | 86 ± 6 |
| | | 500 | 54 ± 9 |
| | | 100 | 42 ± 10 |

**(ii) Simulations with no admixture in the target population**

To investigate if *DATES* gives spurious results for admixture in the absence of gene flow from the reference populations, we generated data for populations without a history of recent admixture. We simulated individuals for three populations *PopA*, *PopB,* and *PopC*, where the divergence between *PopA* and *PopB* was 1800 generations and divergence between *PopC* and *PopB* was 1000 generations. *PopC* had a bottleneck that occurred $T_B$ (=100 or 10) generations ago where the population size reduced to $N_B$ (=100 or 10). Applying *DATES* with *PopC* as the target with *PopA* and *PopB* as reference populations, we observed no evidence of ancestry decay in *PopC* — the ancestry covariance curves were noisy and the 95% CI for the dates included 0 (Figure S2.15).

*macs command line:*

macs 120 1e8 -t 6e-4 -r 5e-4 -I 3 50 20 50 -en 0.0002 2 0.0002 -en 0.0003 2 1 -ej 0.02 2 3 -ej 0.036 1 3

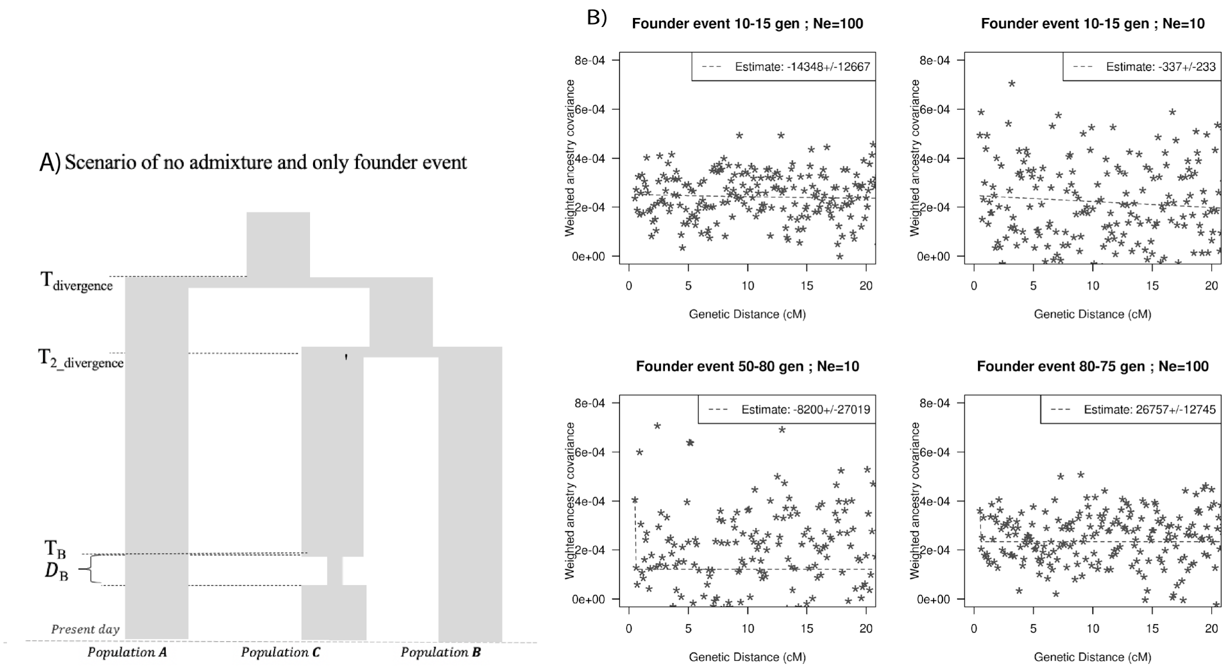


**Figure S2.15: Impact of models with no admixture, severe founder event: A)** Demographic scenario with three populations. *PopA* and *PopB* diverged 1800 generations ago and *PopB* and *PopC* diverged 1000 generations ago. *PopC* had a bottleneck at $T_B$ generations ago with population size during bottleneck $N_B$. B) Ancestry covariance curves for *PopC*. We simulated data for 25 individuals from *PopA* and *PopB* and 10 individuals from *PopC*. We applied *DATES* on *PopC* with *PopA* and *PopB* as sources and show the decay curves for different timing and effective population size of founder events. Note, none of the simulations show significant exponential fits and all dates include 0 in the estimated CI.

Further, we simulated a target population with a severe bottleneck without recovery to present (without any admixture), where *PopC* undergoes bottleneck at 100 or 500 generations without recovery till present. The effective population size reduces from 12,500 to 1000 or 650 at 100 generations or 500 generations and is maintained at the low size to present (Figure S2.16). Using *DATES* on *PopC* as a target with *PopA* and *PopB* as sources, we observed that the ancestry covariance curves were noisy and the dates were not significant. This shows that *DATES* does not provide spurious evidence for admixture even for populations that have a complex history with strong founder events.

*macs command line:*

macs 120 1e8 -t 6e-4 -r 5e-4 -I 3 50 20 50 -en 0 2 0.013 -en 0.00202 2 1 -ej 0.00204 2 3 -ej 0.036 1 3

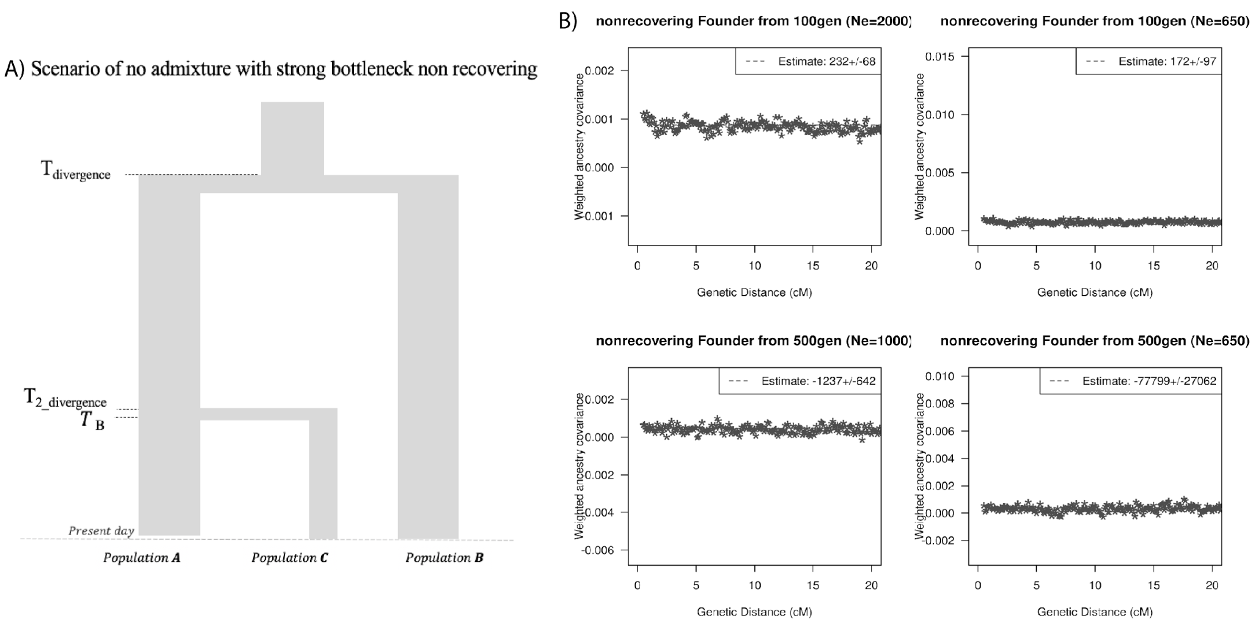


**Figure S2.16: Effect of drift, no admixture: Impact of models with no admixture, severe founder event (without recovery): A)** Demographic scenario with three populations. *PopA* and *PopB* diverged 1800 generations ago and *PopB* and *PopC* diverged 1000 generations ago. *PopC* had a bottleneck at $T_B$ generations ago with population size during bottleneck $N_B$. The population size $N_B$ is maintained to present. B) Ancestry covariance curves for *PopC*. We simulated data for 25 individuals from *PopA* and *PopB* and 10 individuals from *PopC*. We applied *DATES* on *PopC* with *PopA* and *PopB* as sources and show the decay curves for different timing and effective population size of founder events.

#### 10. Comparison with ALDER

To compare the *DATES* with ALDER, we simulated data for $n$ (=20 or 100) admixed individuals using our admixture simulator with 20% European and 80% African ancestry using CEU and YRI phased individuals from 1000 Genomes Project (5). We used French and Yoruba from the Human Genome Diversity Panel (HGDP) dataset (6) as the reference populations to represent European and African source populations respectively. We applied *DATES* and ALDER to the same dataset. We ran ALDER using default settings and allowed ALDER to pick the minimum distance to start the exponential fit. ALDER estimates the date of admixture by fitting an exponential to the weighted covariance statistic with genetic distance and performs a least-squares fit using $y = Ae^{-td} + c$, where $d$ is the genetic distance in Morgans and $t$ is the number of generations since admixture. We note this differs from *DATES* which assumes the exponential decay parameter of $(t+1)$, though in practice this has little effect on the comparisons. For the timing of admixture between 10-200 generations, we observed that both methods accurately estimated the time of admixture in most cases, though *DATES* provided more precise estimates than ALDER for older admixture dates (Table S2.4).

**Table S2.4: Comparison of ALDER and *DATES* for varying samples sizes and times of admixture.** We simulated data for 20 and 100 admixed individuals using the CEU and YRI from 1000G with the mixture proportion of 20% from European and 80% African ancestry. The dates reported here for *DATES* are using exponential fit to $\lambda - 1$ generations.

| Time of admixture (gen) | Number of individuals, *n*=20 | | Number of individuals, *n*=100 | |
|---|---|---|---|---|
| | ALDER mean ± 1SE (gen) | *DATES* mean ± 1SE (gen) | ALDER mean ± 1SE (gen) | *DATES* mean ± 1SE (gen) |
| 10 | 9.3 ± 0.8 | 10.7 ± 0.6 | 10.2 ± 0.3 | 10 ± 0.3 |
| 20 | 19.4 ± 1.3 | 19.7 ± 0.8 | 20.2 ± 0.3 | 20.3 ± 0.3 |
| 30 | 28.5 ± 1.7 | 30.8 ± 1.5 | 30.6 ± 0.9 | 30.5 ± 0.7 |
| 40 | 40.9 ± 2 | 40.3 ± 1.5 | 40.6 ± 0.7 | 40.6 ± 0.4 |
| 50 | 47.9 ± 3.6 | 49.6 ± 1.6 | 50 ± 1.1 | 50.9 ± 0.7 |
| 60 | 55.7 ± 2.7 | 60.3 ± 1.5 | 62 ± 2.2 | 63.2 ± 1 |
| 70 | 71.4 ± 4 | 74 ± 2.7 | 74 ± 2.2 | 72.4 ± 1.3 |
| 80 | 80.6 ± 4.8 | 82.5 ± 2.9 | 85.3 ± 2.3 | 84.4 ± 1.1 |
| 90 | 87.8 ± 4.2 | 88.9 ± 3 | 94.1 ± 2.7 | 92.9 ± 1.3 |
| 100 | 93.7 ± 4.9 | 98.1 ± 2.9 | 101.9 ± 3.9 | 103.6 ± 1 |
| 110 | 121.4 ± 5.4 | 118.2 ± 3.7 | 120.7 ± 4.3 | 115.5 ± 1.8 |
| 120 | 116.5 ± 8.5 | 128.4 ± 3.9 | 121.2 ± 5.1 | 121.5 ± 1.7 |
| 130 | 138.2 ± 9.2 | 133.7 ± 4.6 | 130.2 ± 4.8 | 132.8 ± 1.7 |
| 140 | 134.5 ± 17.5 | 142.4 ± 7 | 144.9 ± 7.3 | 145.3 ± 3.1 |
| 150 | 144.8 ± 23.8 | 149.5 ± 7.4 | 155.1 ± 7 | 157.5 ± 2.8 |

| | | | | |
|---|---|---|---|---|
| 160 | 141.9 ± 11.3 | 166.7 ± 5.9 | 154.5 ± 8.5 | 161.7 ± 2.4 |
| 170 | 173.4 ± 13.7 | 175.1 ± 6.9 | 170.3 ± 6.2 | 173.5 ± 3 |
| 180 | 204.6 ± 17.8 | 195.5 ± 7.1 | 174.2 ± 7 | 180.7 ± 3.3 |
| 190 | 221.3 ± 23.9 | 210.4 ± 9.4 | 191.2 ± 16.2 | 197.2 ± 4.6 |
| 200 | 202.8 ± 11.1 | 196 ± 6.5 | 188.5 ± 16.5 | 202.6 ± 4.7 |

Next, we generated 10 simulated individuals with missing genotypes varying between 5–60% (in increments of 5%) as described in Note S2.1 and applied both *DATES* and ALDER. Using the same setup in both methods, we inferred that *DATES* reliably recovers the time of admixture even with high missing proportions such as 60% (Figure S2.6). However, ALDER becomes every noisy with large proportions of missing data (>40%). For older dates (>100 generations), we observed biased estimates even with >10% missing genotypes (Figure S2.18) As the missing sites vary among individuals, admixture-LD based methods such as ALDER that combine information across individuals become noisy as there are few sites without non-missing genotypes remaining for inference. However, *DATES* performs the analysis for single individuals (using all non-missing genotypes for that individual) and then averages the inferred estimates across individuals. This provides substantial robustness to variable missingness across individuals.

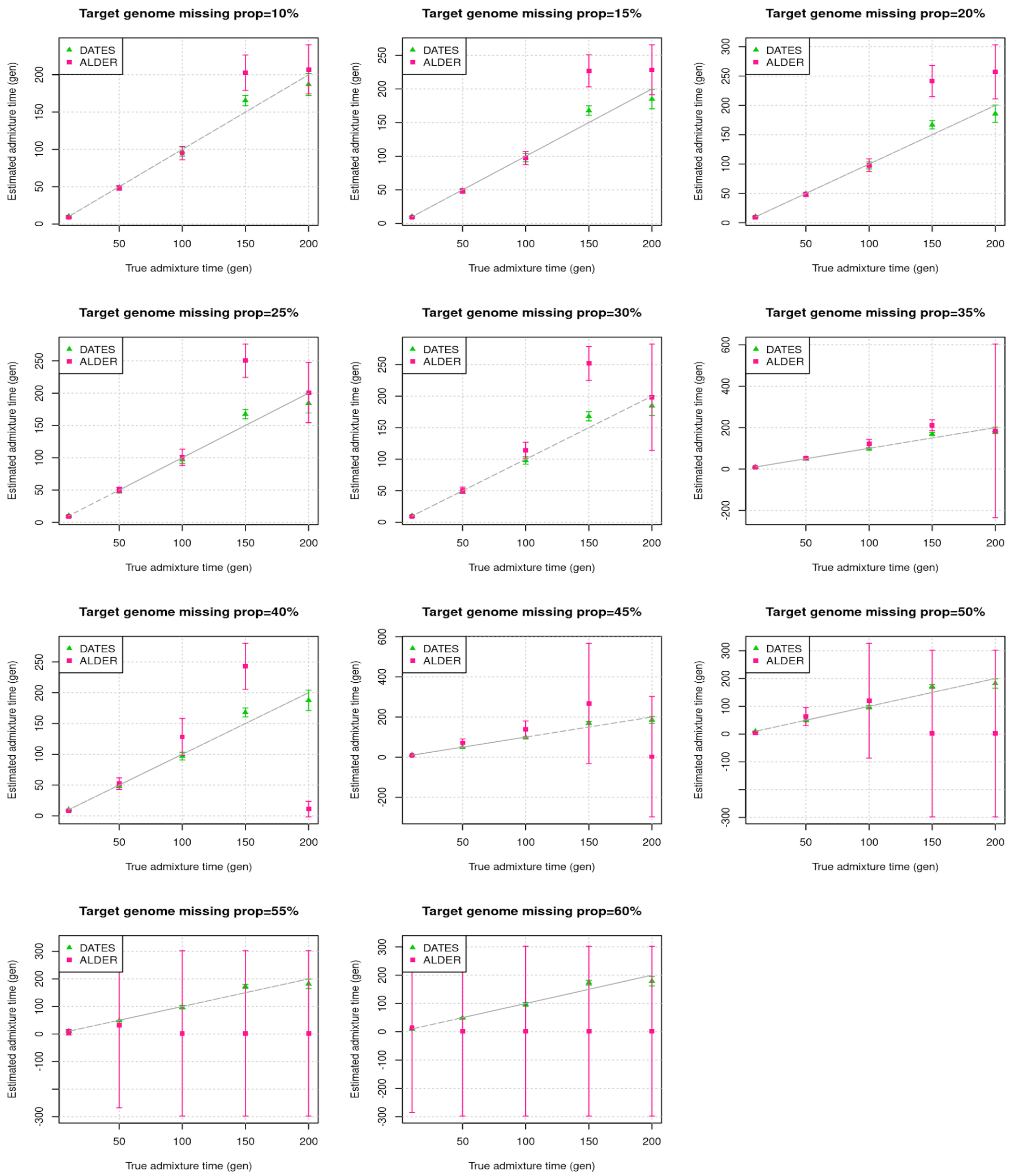


**Figure S2.17: Effect of missing genotypes on the performance of *DATES* and ALDER**: We simulated data for 10 admixed individuals with varying proportions of missing data (shown in each panel). The estimated admixture times (± 1 SE) from *DATES* (green) and ALDER (pink) are shown on Y-axis and the true time of admixture is shown on X-axis. For a fair comparison with ALDER, the dates reported here for *DATES* are using exponential fit to $\lambda - 1$ generations (instead of the default of $\lambda$ generations).

### Note S3. Comparison of versions of *DATES*

An earlier version of *DATES* (version v753) was released in Narasimhan et al. (2019) available on GitHub (https://github.com/priyamoorjani/DATES.git). This version is similar to the current version of *DATES* (v3600) though it differs in implementation. The key differences are:
a) Use of regression model vs. likelihood approach: In v753, we used a regression model to infer the residuals at each site in the genotype by conditioning on the allele frequency in the reference population and the genome-wide estimate of the admixture proportion. Instead in v3600, we use a more rigorous likelihood framework where we infer the probability of ancestry from each reference population at each site in the genome.

b) Fitting an exponential: In v753, like ALDER and Rolloff, we fit an exponential decay with the rate of $t$ generations. However, this assumes that mosaic chromosomes are formed in the generation when the gene flow occurs. However, in reality, the mixing of ancestry only begins in the following generations as the chromosomes of distinct ancestry recombine. To correctly account for this effect, we fit an exponential with the rate of ($t$+1) in *DATES* v3600. In practice, this has a minor effect on the dates reported earlier, as in most cases the uncertainty is much larger than one generation.

c) Normalized Root-Mean-Square Deviation (NRMSD) for evaluating the exponential fit: In v3600, we implemented the NRMSD to assess the goodness of fit of the exponential curve. NRMSD computes the deviation between the empirical estimate and fitted data in order to provide a statistical way to characterize the noisiness of the fitted curve. Lower values of NRMSD suggest a better fit, however, there is no clear interpretation of the absolute value of NRMSD. Based on the empirical distribution of NRMSD values in our study samples (Figure S1.3), we infer a conservative threshold of 0.7 to define a "good" fit. We caution that users should adjust this threshold based on their application and always visually inspect their exponential fits to ensure reliable results.

We compared the versions of *DATES* by performing simulations by generating 10 admixed individuals with 20% European and 80% African ancestry where the time of admixture varied between 10–300 generations (similar to Note S2). We also varied the sample sizes of the admixed population between 1-20 in increments of 5. Our estimated admixture using v753 and v3600 are highly concordant suggesting although the implementation has changed, the results are similar (Table S3.1A-B). Further, we compared the dates of admixture that were reported using the earlier version. To this end, we repeated the analysis for Narasimhan et al. (2019) for ancient South Asians and Rivollat et al. (2020) for ancient Neolithic samples in Europe (15). In both cases, we obtained consistent results as reported earlier (Table S3.2, Table S3.3).

**Table S3.1: Comparison of results using *DATES* v753 and v3600 using simulated data.**

**A) Simulated data with the target sample size (*n*) of 10 individuals**

| True time of admixture (generations) | *DATES* (V753) (mean ± SE) | *DATES* (v3600) (mean ± SE) |
|---|---|---|
| 10 | 10.0 ± 0.5 | 11.0 ± 0.6 |
| 20 | 19.1 ± 1.5 | 19.6 ± 1.5 |
| 30 | 28.0 ± 1.5 | 28.9 ± 1.3 |
| 40 | 46.1 ± 2.1 | 45.7 ± 1.8 |
| 50 | 55.5 ± 2.9 | 55.6 ± 2.8 |
| 60 | 59.4 ± 2.2 | 60.5 ± 2.4 |
| 70 | 69.3 ± 4.2 | 69.8 ± 4.0 |
| 80 | 84.2 ± 4.4 | 84.0 ± 3.9 |
| 90 | 97.7 ± 3.7 | 93.7 ± 3.6 |
| 100 | 107.4 ± 5.4 | 106.7 ± 4.5 |
| 110 | 113.6 ± 5.5 | 112.9 ± 4.7 |
| 120 | 122.7 ± 5.4 | 124.3 ± 5.5 |
| 130 | 138.6 ± 7.7 | 134.1 ± 6.2 |
| 140 | 153.2 ± 9.2 | 152.0 ± 8.4 |
| 150 | 147.5 ± 9.0 | 146.6 ± 8.2 |
| 160 | 181.4 ± 9.8 | 176.6 ± 8.2 |
| 170 | 178.0 ± 8.1 | 175.9 ± 7.4 |
| 180 | 180.7 ± 10.6 | 182.3 ± 9.3 |
| 190 | 172.9 ± 15.3 | 174.8 ± 12.7 |
| 200 | 204.7 ± 17.1 | 208.8 ± 13.4 |
| 210 | 194.5 ± 13.3 | 196.3 ± 11.9 |
| 220 | 255.8 ± 17.0 | 250.7 ± 13.8 |
| 230 | 251.9 ± 18.6 | 237.0 ± 13.0 |
| 240 | 234.7 ± 18.3 | 241.5 ± 14.2 |
| 250 | 228.3 ± 13.8 | 233.2 ± 11.9 |
| 260 | 254.2 ± 21.7 | 253.0 ± 16.1 |
| 270 | 291.4 ± 22.4 | 292.1 ± 20.0 |
| 280 | 252.4 ± 25.2 | 248.1 ± 22.4 |
| 290 | 277.9 ± 22.4 | 285.4 ± 20.5 |
| 300 | 318.6 ± 23.3 | 315.1 ± 20.3 |

**B) Simulated data with sample size (*n*) ranging between 10-20 individuals.**

| True time of admixture (generations) | Sample size | *DATES* (V753) (mean ± SE) | *DATES* (v3600) (mean ± SE) |
|---|---|---|---|
| 10 | 1 | 6.9 ± 2.5 | 7.8 ± 2.6 |
| 10 | 5 | 8.8 ± 0.8 | 9.9 ± 0.8 |
| 10 | 10 | 9.9 ± 1.2 | 10.8 ± 1.2 |
| 10 | 15 | 10.9 ± 0.7 | 11.8 ± 0.7 |
| 10 | 20 | 10.3 ± 0.6 | 11.3 ± 0.6 |
| 50 | 1 | 51.7 ± 7.1 | 51.5 ± 7.5 |
| 50 | 5 | 59.7 ± 3.5 | 58.6 ± 3.2 |
| 50 | 10 | 48.7 ± 2.7 | 50.1 ± 2.7 |
| 50 | 15 | 54.2 ± 2.1 | 54.7 ± 2.1 |
| 50 | 20 | 52.9 ± 1.9 | 53.1 ± 1.9 |
| 100 | 1 | 124.5 ± 17.6 | 122.5 ± 13.2 |
| 100 | 5 | 107.2 ± 7.6 | 108.2 ± 7.5 |
| 100 | 10 | 103.3 ± 7.7 | 103.2 ± 7.8 |
| 100 | 15 | 99.4 ± 4.5 | 100.1 ± 3.8 |
| 100 | 20 | 105.5 ± 3.9 | 103.4 ± 3.3 |
| 150 | 1 | 136.4 ± 29.2 | 144.2 ± 26.3 |
| 150 | 5 | 142.6 ± 11.4 | 143 ± 11.7 |
| 150 | 10 | 156.9 ± 9 | 158.6 ± 7 |
| 150 | 15 | 142.9 ± 7.8 | 146.1 ± 6.7 |
| 150 | 20 | 156.5 ± 5.4 | 152.9 ± 4.3 |
| 200 | 1 | 195.4 ± 88.4 | 160.2 ± 73.9 |
| 200 | 5 | 210.9 ± 20.7 | 206.7 ± 18.7 |
| 200 | 10 | 225 ± 18.7 | 219.8 ± 18 |
| 200 | 15 | 200 ± 10.6 | 197.7 ± 9 |
| 200 | 20 | 189.4 ± 11 | 190.3 ± 9 |

**Table S3.2: Comparison of results with Narasimhan, Patterson et al. 2019.**

| **Pop** | **Reference populations*** | ***DATES* (v753)** | ***DATES* (v3600)** |
|---|---|---|---|
| | | **(mean ± SE; in generations)** | **(mean ± SE; in generations)** |
| *Indus_Periphery_Pool* | AASI and Iranian-farmer-related | 71 ± 15 | 62 ± 7 |
| *SPGT* | AASI and Steppe-pastoralist-related | 26 ± 3 | 28 ± 3 |

Note:
* We used the reference populations of AASI ancestry that includes South Asians from the 1000 Genomes Project (Phase 3) including Sri Lankan Tamil from the UK (*STU.SG*) and Indian Telugu from the UK (*ITU.SG*), as well as *BIR.SG* and Iranian farmer-related ancestry including Aigyrzhal_BA, Sarazm_EN, Geoksyur_EN, Parkhai_Anau_EN, and Steppe-pastoralist-related including Central_Steppe_MLBA.

**Table S3.3: Comparison of dates of the spread of Neolithic farming from Rivollat et al. 2020.**

| **Population** | ***n*** | ***DATES* (v753)** | **Population in our study (v44 1240K)** | ***n* (v44 1240K)** | ***DATES* (v3600)#** |
|---|---|---|---|---|---|
| Bulgaria_MP_Neolithic | 9 | 8.4 ± 2.3 | Bulgaria_MalakPreslavets_N | 3 | 8.05 ± 3 |
| Serbia_Neolithic | 4 | -- | Serbia_EN | 3 | 22.8 ± 9.8 |
| Romania_EN | 2 | 32.1 ± 10.4 | Romania_EN* | 2 | 29.7 ± 7.1 |
| Croatia_Impressa | 2 | -- | Croatia_EN_Impressa | 2 | -- |
| Hungary_ALPc_MN | 23 | 21.5 ± 4.7 | Hungary_MN_ALPc | 21 | 21.9 ± 1.6 |
| Hungary_LBK_MN | 10 | 12.8 ± 5.2 | Hungary_MN_LBK | 6 | 18.6 ± 7.4 |
| Hungary_ALBK_MN | 2 | 14.8 ± 3.2 | Hungary_MN_ALBK_Szakalhat | 2 | 19.3 ± 3.3 |
| Hungary_LN | 18 | 21.5 ± 3.7 | Hungary_LN | 18 | 28.03 ± 3.8 |
| Austria_LBK_EN | 8 | 15.5 ± 4.6 | Austria_EN_LBK | 9 | 17.6 ± 2.3 |
| Czech_MN | 5 | 18.3 ± 7.7 | Czech_MN | 4 | 32.9 ± 6.3 |
| France_MN | 3 | 26.5 ± 5.6 | France_MN | 43 | 30 ± 1.3 |
| Iberia_EN | 10 | 15.6 ± 2.5 | Spain_EN | 11 | 20.6 ± 3.6 |
| Iberia_MN | 7 | 52.4 ± 4.3 | Spain_MLN | 42 | 56.3 ± 4 |
| Germany_LBK_EN | 27 | 14.4 ± 2.6 | Germany_EN_LBK | 54 | 17.4 ± 2.7 |
| Germany_Blatterhohle_MN | 4 | 12.3 ± 2.5 | Germany_Blatterhohle_MN | 4 | 16.2 ± 2.9 |
| Germany_Esperstedt_MN | 1 | -- | Germany_MN_Esperstedt | 1 | -- |
| England_Neolithic | 29 | 45.5 ± 5.5 | England_N.SG | 17 | -- |
| Wales_Neolithic | 6 | 45.3 ± 7.4 | Wales_N | 4 | 50.7 ± 3.3 |
| Scotland_Neolithic | 42 | 50.9 ± 3.8 | Scotland_N | 30 | 56.6 ± 2.9 |
| Ireland_Neolithic | 13 | 46.9 ± 7.5 | Ireland_MN.SG | 26 | 50.8 ± 2.2 |

Note:
(blue) indicates samples sizes that differ across both studies
### For *DATES*, we used pooled WHG and Anatolian farmers as the reference populations except for samples marked with *.
-- indicates cases where the results were not significant as the 95% CI includes 0

### Note S4. Comparison to other published methods

We compared the performance of *DATES* with other published methods such as Rolloff (4), ALDER (2), and Globetrotter (13). For this purpose instead of re-running all the tools, we leveraged published results from Hellenthal et al. 2014 (Table S12) where the authors compared the performance of Globetrotter and Rolloff (13). Following (13), we generated a merged dataset including samples from the Human Genome Diversity Panel (HGDP) (6), Behar et al. 2010 (16), and Henn et al. (17). Our dataset consists of 1,642 individuals and 465,543 SNPs. We did not perform any additional quality control of the dataset, except to check that the sample sizes were similar for the target and reference populations as reported in Table S12. We excluded one population "Indian" where the population label was ambiguous and could refer to many populations in the merged dataset.

We applied *DATES* and ALDER to 29 target groups using the reference populations reported in Hellenthal et al. 2014 (Table S12). We obtained significant dates of admixtures in 20/ 29 groups using *DATES*. Out of these, the estimated dates between Globetrotter and *DATES* were consistent for 14 populations (within two standard errors). In the case of the six populations that disagreed across the two methods, most of the populations appear to have a history of multiple pulses of gene flow either involving more than two populations or multiple instances of contact between the same two reference groups. For instance, previous studies have documented that Brahui has a multi-way admixture with South Asian ancestry (which are themselves admixed) (18) and sub-Saharan African ancestry (19). In this case, the result from *DATES* infers the more recent admixture, while Globetrotter infers an older date (Table S4.1). Another population, present-day Bulgarians, have ancestry from western hunter-gatherers, Near Eastern farmers, and Steppe pastoralists from Eurasia (see main text). Globetrotter instead models this group as a two-way admixture between Polish and Cypriots (Table S4.1). The results, in this case, are hard to interpret as depending on the composition of the ancestral populations either of the two ancient events or other more recent events could be captured. Finally, in the case of the west African population of Mandenka, the admixture likely occurred at multiple time points, given Mandenka's geographical location is close to North Africa (20). In these cases, Globetrotter and *DATES* could be capturing different events or the weighting of both events could differ.

We ran ALDER using default settings and allowed ALDER to pick the minimum distance and evaluate the model of admixture by comparing the one reference and two reference results. ALDER estimates the date of admixture by fitting an exponential to the weighted covariance statistic with genetic distance and performs a least-squares fit using $y = Ae^{-td} + c$, where $d$ is the genetic distance in Morgans and $t$ is the number of generations since admixture. We note this differs from *DATES* and Globetrotter which assumed the exponential decay parameter of $(t + 1)$, though in practice this has little effect on the comparisons. We observed that ALDER's formal test failed in most cases (25/29) (Table S4.1). The two main reasons for this are:

- 7/29 due to long-range shared LD where one of the reference populations appeared to be closely related to the target group;
- 18/29 due to differences in dates of single reference and two reference setups

However, the estimated dates for ALDER assuming two reference model (regardless of the formal test results) were highly concordant with *DATES*. The results of *DATES* were also highly concordant with Rolloff except for one group - Mandenka (Table S4.1).

**Table S4.1: Comparison of *DATES* and published admixture dating methods (Rolloff, Globetrotter, ALDER).**

| Population | nk | Source1 | Source2 | Rolloff | Globetrotter | ALDER formal test | ALDER_2-ref dates | *DATES* | Comments |
|---|---|---|---|---|---|---|---|---|---|
| Hazara | 22 | Mongola (10) | Iranian (13) | 23 ± 1 | 22 ± 0.9 | Long-range LD | -- | 24.6 ± 1.0 | |
| Uzbekistani | 15 | Mongola (10) | Iranian (13) | 20 ± 1.4 | 19 ± 1.1 | **SUCCEEDS** | 19.18 ± 2.22 | 21.3 ± 1.4 | |
| Uyghur | 10 | Mongola (10) | Iranian (13) | 23 ± 2.6 | 22 ± 1.3 | **SUCCEEDS** | 16.73 ± 1.38 | 22.2 ± 2.1 | |
| Makrani | 22 | Bantu Kenya (11) | Balochi (21) | 18 ± 1.8 | 18 ± 1.2 | Long-range LD | -- | 13.2 ± 1.6 | |
| Druze | 42 | Yoruba (21) | Cypriot (12) | 39 ± 7.3 | 37 ± 1.9 | FAILS | 44.02 ± 6.37 | 43.4 ± 6.1 | |
| Mozabite | 25 | Yoruba (21) | Moroccan (22) | 23 ± 1.9 | 21 ± 1.3 | Long-range LD | -- | 21.6 ± 1.8 | |
| Turkish | 17 | Mongola (10) | Iranian (13) | 28 ± 3.2 | 24 ± 1.5 | FAILS | 25.62 ± 2.48 | 28.5 ± 2.3 | |
| Brahui | 23 | Bantu Kenya (11) | Balochi (21) | 13 ± 3.4 | 20 ± 1.5 | Long-range LD | -- | 10.4 ± 1.6* | Possibly multi-way admixture (19) |
| Yemeni | 4 | Bantu Kenya (11) | Syrian (16) | 15 ± 2.3 | 14 ± 1.8 | FAILS | 6.29 ± 2.89 | 12.7 ± 1.6 | |
| Pima | 14 | Turkish (17) | Mayan (21) | 9 ± 3.6 | 6 ± 0.9 | **SUCCEEDS** | 6.29 ± 0.89 | 7.8 ± 1.1 | |
| Bantu South Africa | 8 | San Khomani (30) | Yoruba (21) | 26 ± 2.5 | 25 ± 2.3 | Long-range LD | -- | 27.9 ± 2.2 | |
| Tu | 10 | Greek (20) | Han N-China (10) | 33 ± 6.3 | 25 ± 2.3 | FAILS | 28.83 ± 2.8 | 31.3 ± 1.96 | |
| West Sicilian | 10 | Yoruba (21) | East Sicilian (10) | 26 ± 7.8 | 27 ± 3.9 | FAILS | 42.72 ± 16.34 | 37.4 ± 16.4 | |
| Cambodian | 10 | Uyghur (10) | Han (34) | 17 ± 4.7 | 20 ± 2.7 | **SUCCEEDS** | 24.28 ± 5.36 | 33.6 ± 3.7* | |
| Georgian | 20 | Adygei (17) | Greek (20) | -- | 30 ± 3.3 | FAILS | 3.15 ± 1.22 | -- | |
| Romanian | 13 | Lithuanian (10) | East Sicilian (10) | -- | 31 ± 2.6 | FAILS | -- | -- | |
| Bulgarian | 18 | Polish (16) | Cypriot (12) | -- | 28 ± 3.5 | FAILS | 40.95 ± 16.42 | 91.1 ± 24.7* | Possibly multi-way admixture (see Main text) |

| | | | | | | | | | |
|---|---|---|---|---|---|---|---|---|---|
| Hezhen | 8 | Tujia (10) | Mongola (10) | -- | 13 ± 1.3 | FAILS | 2.92 ± 1.4 | -- | |
| Oroqen | 9 | Yakut (25) | Mongola (10) | -- | 15 ± 2 | Long-range LD | -- | -- | |
| Hungarian | 18 | Cypriot (12) | Polish (16) | 65 ± 24 | 39 ± 3.5 | FAILS | 54.83 ± 25.27 | 61.8 ± 19.1 | |
| Han N-China | 10 | Turkish (17) | Tujia (10) | 37 ± 11.1 | 26 ± 3.8 | FAILS | 48.17 ± 10.36 | 44.3 ± 5.1* | |
| Daur | 9 | Tujia (10) | Mongola (10) | -- | 21 ± 1.7 | FAILS | -- | -- | |
| Greek | 20 | Polish (16) | Cypriot (12) | 69 ± 18.5 | 36 ± 3.7 | FAILS | 55.54 ± 8.93 | 62.6 ± 16.9 | |
| Melanesian | 10 | Papuan (16) | Cambodian (10) | 66 ± 12.1 | 28 ± 7.6 | FAILS | 64.91 ± 5.42 | 68.6 ± 7.1* | |
| Mandenka | 22 | Moroccan (22) | Yoruba (21) | 22 ± 10.3 | 19 ± 4.2 | FAILS | 17.25 ± 6.05 | 85.8 ± 19.0 *#Ω | Possibly multiple admixture events (20) |
| Indian | 13 | Cambodian (10) | Sindhi (23) | 91 ± 41.1 | 53 ± 8.4 | FAILS | n/a | n/a | There are multiple "Indian" groups in the dataset making it unclear which target was used |
| North Italian | 12 | Cypriot (12) | French (28) | -- | 71 ± 11.8 | FAILS | 12.44 ± 4.32 | -- | |
| Polish | 16 | French (28) | Lithuanian (10) | -- | 31 ± 5.1 | FAILS | -- | -- | |
| Tuscan | 8 | Cypriot (12) | French (28) | -- | 35 ± 6.1 | FAILS | -- | -- | |
| San Namibia | 5 | Sandawe (28) | San Khomani (30) | -- | 48 ± 8.9 | Long-range LD | -- | -- | |

NOTE

- Columns 1-5 include results from Table S12 from Hellenthal et al. 2014. We only show significant dates (|Z| > 2)
- Following Hellenthal et al., we created a merged dataset of the Human Genome Diversity Panel, Henn et al. and Behar et al. containing 1642 individuals and 465543 SNPs. This dataset was used for ALDER and *DATES* analysis.
- Standard errors in *DATES* were estimated using chromosome jackknife (see Methods)

-- indicates results where the inferred results were not significant, either the method failed or the 95% CI included 0

*- indicates *DATES* estimates that significantly differ from Globetrotter estimates (not within 2 SE)

#- indicates *DATES* estimates that significantly differ from Rolloff results

Ω- indicates *DATES* estimates that significantly differ from ALDER results

n/a- indicates target population was unclear

### Note S5. Modeling population mixture in ancient Europe

To understand the admixture history of ancient Europeans, we used *qpAdm* (8) to examine the sources of ancestry in diverse ancient European groups from the Human Origins 1240K dataset (21). Following previous studies (2, 8, 15, 22), we used a core set of seven outgroups (O7) containing Mbuti Pygmies, Ethiopian Mota, Russian Ust' Ishim hunter-gatherers (HG), Russian Mal'ta HG, Papuan, Onge, and Han that are symmetrically related to all ancient European individuals. For some analysis, we supplemented this set with additional outgroups that were necessary (see Model B-D). For all analyses, we used the default parameters of *qpAdm*. We chose the most parsimonious model, i.e., fitting the data with the minimum number of sources for each set of results. We focused on models where the p-value > 0.05. For the results described below, we considered the following models:

**Model-A**: O7 (Mbuti Pygmies, Ethiopian Mota, Russian Ust' Ishim HG, Russian Mal'ta HG, Papuan, Onge, and Han)

**Model-B:** O7, WHG and EHG

**Model-C**: O7 and EHG

**Model-D**: O7, EHG and Levant HG

**Model-E**: O7 and Italy North Villabruna HG

##### Mesolithic hunter-gatherers

We applied *qpAdm* to test if the individuals from central Europe, the Baltic region, and Scandinavia can be modeled as a mixture of western hunter-gatherers (WHG) and eastern hunter-gatherers (EHG). Using Model-A and WHG and EHG as sources, we obtained a good fit for most groups, except Estonia MN Comb Ware Culture. We report the details of estimated proportions in Supplementary Table S5.1.1. To confirm that the target populations do not harbor Anatolian farmer-related ancestry, we applied *D*-statistics of the form *D*(Mbuti, *target*, WHG, Anatolian farmers) where *target* = Mesolithic hunter-gatherers. We observed that none of the target groups have a stronger affinity to Anatolian farmers compared to WHG (Supplementary Table S5.2). Additionally, for Iron Gates HGs, we also tried a three-way model with Anatolian-related groups as previous studies have shown that the Iron Gates HGs have a strong affinity to Anatolian hunter-gatherers (AHG), compared to other European HGs. To this end, we used Model-A with WHG, EHG, and AHG as sources. We showed that this model is feasible (*p*-value > 0.05), however, the ancestry proportion of the AHG is not significant (Supplementary table S5.1.2).

**Supplementary Table S5.1.1: *qpAdm* analysis of Mesolithic hunter-gatherers.** We modeled the Mesolithic hunter-gatherers (target) from Europe as a two-way mixture with sources as WHG and EHG using Model-A.

| Target | Region | WHG | EHG | SE_WHG | SE_EHG | p-value |
|---|---|---|---|---|---|---|
| Norway_Mesolithic.SG | Scandinavia | 0.250 | 0.750 | 0.059 | 0.059 | 0.433 |
| Norway_N_HG.SG | Scandinavia | 0.267 | 0.733 | 0.070 | 0.070 | 0.113 |
| Sweden_HG.SG | Scandinavia | 0.875 | 0.125 | 0.132 | 0.132 | 0.215 |
| Sweden_Motala_HG | Scandinavia | 0.438 | 0.562 | 0.041 | 0.041 | 0.325 |
| Sweden_Mesolithic.SG | Scandinavia | 0.480 | 0.520 | 0.050 | 0.050 | 0.372 |
| Estonia_EMN_Narva | Baltic sea region | 0.476 | 0.524 | 0.068 | 0.068 | 0.649 |
| Estonia_MN_CCC | Baltic sea region | 0.240 | 0.760 | 0.083 | 0.083 | *0.004* |
| Estonia_N_CombCeramic.SG | Baltic sea region | 0.172 | 0.828 | 0.090 | 0.090 | 0.846 |
| Latvia_HG | Baltic sea region | 0.604 | 0.396 | 0.031 | 0.031 | 0.622 |
| Latvia_MN | Baltic sea region | 0.606 | 0.394 | 0.053 | 0.053 | 0.710 |
| Latvia_MN_Comb_Ware.SG | Baltic sea region | 0.054 | 0.946 | 0.071 | 0.071 | 0.721 |
| Lithuania_EMN_Narva | Baltic sea region | 0.797 | 0.203 | 0.038 | 0.038 | 0.411 |
| Lithuania_Mesolithic | Baltic sea region | 0.903 | 0.097 | 0.068 | 0.068 | 0.722 |
| Ukraine_N | Central Europe | 0.428 | 0.572 | 0.031 | 0.031 | 0.533 |
| Hungary_Koros | Central Europe | 0.936 | 0.064 | 0.061 | 0.061 | 0.077 |
| Iron Gates | Central Europe | 0.850 | 0.150 | 0.029 | 0.029 | 0.272 |

Note: p-values < 0.05 shown in red highlight models that were a poor fit.

**Supplementary Table S5.1.2: *qpAdm* analysis of Iron Gates HGs**. We modeled the Iron Gates hunter-gatherers (target) from Europe as a three-way mixture with sources as WHG, EHG and AHG using Model-A.

| Target | Region | WHG | EHG | AHG | SE_WHG | SE_EHG | SE_AHG | *p*-value |
|---|---|---|---|---|---|---|---|---|
| Iron Gates | Central Europe | 0.638 | 0.224 | 0.138 | 0.109 | 0.043 | 0.072 | 0.926236 |

**Supplementary Table S5.2: *D*-statistics to assess the affinity of Mesolithic HG groups to WHG or Anatolian farmers.** We performed D-statistics *D*(Mbuti, target, WHG, Anatolian farmers) where target = Mesolithic hunter-gatherers. A negative *D*-score indicates the target shares more alleles with WHG than Anatolian farmers.

| Target | *D*-score (Z-score) | Target closer to WHG or Anatolian farmer |
|---|---|---|
| Norway_Mesolithic.SG | -0.1028 (-32.043) | WHG |
| Norway_N_HG.SG | -0.1023 (-27.869) | WHG |
| Sweden_HG.SG | -0.0805 (-17.809) | WHG |
| Sweden_Motala_HG | -0.1253 (-42.481) | WHG |
| Sweden_Mesolithic.SG | -0.1243 (-43.667) | WHG |
| Estonia_EMN_Narva | -0.1265 (-38.037) | WHG |
| Estonia_MN_CCC | -0.0895 (-26.438) | WHG |
| Estonia_N_CombCeramic.SG | -0.0922 (-21.145) | WHG |
| Latvia_HG | -0.1515 (-60.979) | WHG |
| Latvia_MN | -0.1367 (-44.476) | WHG |
| Latvia_MN_Comb_Ware.SG | -0.0857 (-24.305) | WHG |
| Lithuania_EMN_Narva | -0.1564 (-56.820) | WHG |
| Lithuania_Mesolithic | -0.1738 (-46.433) | WHG |
| Ukraine_N | -0.0989 (-40.671) | WHG |
| Hungary_Koros | -0.1593 (-46.340) | WHG |
| IronGates | -0.1526 (-60.960) | WHG |

##### The Near Eastern farmers

Previous analysis has suggested that early Anatolian farmers can be modeled as a mixture of Anatolian hunter-gatherers (AHG) and Iran Neolithic. The AHG were in turn were a mixture of WHG and Levant Neolithic groups (23). We confirmed this model using previously suggested sources and outgroups using Model-B (Supplementary Table S5.3). The early Anatolian farmers mixed with Levant Neolithic groups to form the genetic ancestry of Anatolian farmers, that migrated to the west to Europe and in the east to form the Chalcolithic groups of Seh Gabi and Hajji Firuz. Using Model-D and Anatolian farmers and Iran Neolithic groups as sources, we obtained a good fit for Iran Seh Gabi individuals. We find that Hajji Firuz individuals were better modeled as a mix of Iran Seh Gabi and Anatolian farmer-related ancestry using Model-C (24). We report details of all models and estimated admixture proportions in Supplementary Table S5.3.

**Supplementary Table S5.3: *qpAdm* analysis of Neolithic and Chalcolithic Near Eastern Farmers.** We modeled the Anatolian farmers as a two-way mixture of AHG and Iran Neolithic groups using Model-B. The Chalcolithic farmers were modeled as a two-way mixture of Neolithic farmers and Iranian farmers with outgroups from Model-C and Model-D.

**Neolithic Farmers: Early** Anatolian farmers

| Target | Outgroups | AHG | Iran_N | SE AHG | SE_Iran_N | p-value |
|---|---|---|---|---|---|---|
| Early Anatolian farmers | Model-B | 0.891 | 0.109 | 0.044 | 0.044 | 0.964802 |

**Chalcolithic Farmers**

Hajji_Firuz_C

| Target | Outgroups | Iran_C_SehGabi | Anatolian farmers | SE Iran_Seh_Gabi | SE Anatolian farmers | p-value |
|---|---|---|---|---|---|---|
| Hajji_Firuz_C | Model-C | 0.705 | 0.295 | 0.055 | 0.105 | 0.105 |

Iran_C_SehGabi

| Target | Outgroups | Iran_N | Anatolian farmers | SE Iran_N | SE Anatolian farmers | p-value |
|---|---|---|---|---|---|---|
| Iran_C_SehGabi | Model-D | 0.653 | 0.347 | 0.069 | 0.069 | 0.595 |

##### Neolithic Europeans

Following previous studies, we applied the two-way mixture of WHG and Anatolian farmers mixture to data from 94 Neolithic European groups, sampled from diverse regions and time periods in Europe. To this end, we used Model-E with O7 and Italy Villabruna HG as outgroups. For 70/94 groups, this model was found to be a good fit ($p$-value > 0.05) (Supplementary Table SD). For the remaining 24 groups, we investigated if removing outliers or including additional gene flow from other HG groups could provide a good fit for the data. First, we performed *qpAdm* analysis per sample and removed outliers that had a different ancestry source compared to the majority of the individuals in the target population. Using this approach, we found that the model of two-way mixture between WHG and Anatolian farmers was a good fit for 14 groups (Supplementary Table SE). Next, we explored if the source of the HG ancestry was not WHG alone, as many Mesolithic HG groups were admixed and had some proportion of EHG or GoyetQ2 ancestry in addition to WHG. To this end, we added either EHG or GoyetQ2 HGs as additional sources with WHG and Anatolian farmers. In six cases, adding EHG and GoyetQ2 provided a good fit, highlighting the diversity in HG ancestry across Neolithic groups. The model of European HGs and Anatolian farmers failed in six groups ($p$-value < 0.05) (Supplementary Table SD).

To confirm these groups do not have Steppe pastoralist-related ancestry, we applied *D*-statistics of the form *D*(Mbuti, *target*, Anatolian farmers, Yamnaya Steppe pastoralists), where target = Neolithic Europeans. In this setup, if the target shares excess alleles with Steppe pastoralists, then we excluded the target from the *DATES* analysis as in that case the inferred dates would likely reflect the timing of the recent event involving Steppe pastoralists groups (vs. gene flow of HG and Anatolian farmers). Based on this analysis, we excluded 4 groups that showed significantly higher sharing with Steppe pastoralists than Near Eastern farmers (Supplementary Table SF).

##### Eurasian Steppe pastoralists

We explored the model of the ancestry of Steppe pastoralists groups from the early Bronze Age (EBA) and middle and late Bronze Age (MLBA) from Russia, the Urals, and Kazakhstan. For modeling EBA steppe pastoralists groups, we used EHG_pooled and Iran_N pooled as sources (details of the pooling in Table SA) using the outgroups given by Model-A. We observed that this model provides a good fit to most Yamnaya and Afanasievo groups except for two Yamnaya groups from Baden and Kalmykia (Supplementary Table S5.4).

**Supplementary Table S5.4: *qpAdm* analysis of Early Bronze age Steppe pastoralists.** We modeled the EBA Steppe Pastoralists as a two-way mixture of EHG related ancestry and Iranian Farmer ancestry using Model-A.

| Target | EHG pooled | Iran_N pooled | SE_EHG pooled | SE_Iran_N pooled | p-value |
|---|---|---|---|---|---|
| Russia_Samara_EBA_Yamnaya | 0.643 | 0.357 | 0.024 | 0.024 | 0.055 |
| Russia_Afanasievo | 0.658 | 0.342 | 0.021 | 0.021 | 0.285 |
| Ukraine_EBA_Yamnaya | 0.664 | 0.336 | 0.055 | 0.055 | 0.6989 |
| Ukraine_Ozera_EBA_Yamnaya | 0.399 | 0.601 | 0.058 | 0.058 | 0.538 |
| Russia_Caucasus_EBA_Yamnaya | 0.567 | 0.433 | 0.039 | 0.039 | 0.099 |
| Kazakhstan_EBA_Yamnaya.SG | 0.708 | 0.292 | 0.054 | 0.054 | 0.362 |
| Hungary_LateC_EBA_Baden_Yamnaya | -0.142 | 1.142 | 0.042 | 0.042 | 0.252 |
| Russia_Kalmykia_EBA_Yamnaya.SG | 0.633 | 0.367 | 0.035 | 0.035 | *0.035* |

Note: p-values < 0.05 shown in red highlight models that were a poor fit.

Previous analysis has suggested that MLBA Steppe pastoralists groups have ancestry from Yamnaya Steppe pastoralists and Neolithic Europeans (that are a mixture of WHG + Anatolian farmer ancestry) through Corded Ware populations (24). For dating, we used a two-way model containing one reference group containing pooled individuals of (WHG and Anatolian farmers) ancestry and a second reference group as Yamnaya Steppe pastoralists. To confirm this model, we applied *qpAdm* using the same setup. We found this model provides a good fit for all Steppe MLBA populations, except Potapovka from Russia (Supplementary Table S5.5). For Potapovka, we performed *qpAdm* separately for each individual and removed outliers (*p*-value < 0.05), and grouped individuals where the model provides a good fit within the limits of resolution of the method (Supplementary Table S5.6).

**Supplementary Table S5.5: *qpAdm* analysis of Middle late Bronze age (MLBA) Steppe pastoralists.** We modeled the Steppe pastoralists from MLBA as a two-way mixture of EBA steppe groups and Neolithic related ancestry using outgroups Model-A.

| **Target** | **Steppe pastoralists** | **WHG + Anatolian Farmer** | **SE_Steppe pastoralist** | **SE_(WHG+ANF)** | **p-value** |
|---|---|---|---|---|---|
| Sintashta | 0.631 | 0.369 | 0.029 | 0.029 | 0.398 |
| Russia_Potapovka* | 0.795 | 0.205 | 0.133 | 0.133 | 0.426 |
| Andronovo | 0.622 | 0.378 | 0.056 | 0.056 | 0.321 |
| Kazakhstan_Maitan_MLBA_Alakul | 0.625 | 0.375 | 0.037 | 0.037 | 0.346 |
| Russia_Srubnaya_Alakul.S | 0.757 | 0.243 | 0.039 | 0.039 | 0.222 |
| Srubnaya | 0.687 | 0.313 | 0.036 | 0.036 | 0.136 |
| Kazakhstan_MLBA_Aktogai | 0.725 | 0.275 | 0.046 | 0.046 | 0.729 |
| Kazakhstan_MLBA_Kairan | 0.589 | 0.411 | 0.053 | 0.053 | 0.380 |
| Kazakhstan_Shoendykol MLBA_Fedorovo | 0.731 | 0.269 | 0.060 | 0.060 | *0.0003* |

Note: p-values < 0.05 shown in red highlight models that were a poor fit.

**Supplementary Table S5.6: Admixture modeling per sample for MLBA Steppe groups**. Per individual *qpAdm* analysis of Russia Potapovka group using Steppe and pooled samples of WHG and Anatolian farmers as sources using Model-A as outgroups.

| **Instance ID** | **Instance ID** | **Steppe pastoralist** | **WHG+ANF** | **SE Steppe pastoralist** | **SE (WHG+ ANF)** | **p-value** | **Comment** |
|---|---|---|---|---|---|---|---|
| I0418 | Russia_Potapovka | 0.795 | 0.205 | 0.133 | 0.133 | 0.426213 | |
| I0246 | Russia_Potapovka | 1.232 | -0.232 | 0.121 | 0.121 | 0.878173 | infeasible |

Note: p-values < 0.05 shown in red highlight models that were a poor fit.

#### Bronze Age Europeans

We applied *qpAdm* to 109 Bronze Age Europeans from diverse geographic regions in Europe. Following previous publications, we used Model-E that includes the seven standard outgroups along with Villabruna HG that is needed to reliably model the ancestry in Bronze Age individuals (exclusion of this population leads to poor fit for many Bronze Age groups). We found the three-way model with WHG, Anatolian farmers, and Yamnaya Steppe pastoralists as sources that provided a good fit for most groups (83/109) (Supplementary Table SH). For the 28 groups where the previous model failed, we ran *qpAdm* separately for each individual to identify outliers that might be responsible for the model to fail. By excluding outliers, we found the three-way ancestry model was a good fit for 11 additional populations (Supplementary Table SI). Next, we added EHG or GoyetQ2 to test if the source of the HG ancestry was not reliably modeled. The latter had minimal effect for any target population. We excluded 17 groups that could not be modeled as a three-way mixture with the tested source populations (Supplementary Table S16). Further, to ensure that all samples had Steppe pastoralist-related ancestry, we performed a model competition to test if the two-way including WHG and Anatolian farmers or the three-way model including WHG, Anatolian farmers and Steppe pastoralists provides a better fit to the data. We excluded 20 groups out of 111 groups as the parsimonious model including only two reference populations was a better fit in these cases (Supplementary Table SH). After filtering, we retained 79 Bronze Age groups for dating the spread of Steppe pastoralist-related ancestry across Europe.

### Note S6. Formation of early Steppe pastoralists groups

The beginning of the Bronze Age was accompanied by the spread of Steppe-pastoralist-related groups from the Pontic-Caspian steppe region into Europe. These groups derived most of their ancestry from populations related to early Bronze Age populations of Yamnaya and Afanasievo from the Eurasian steppe (8). To understand the formation of the early Steppe pastoralists, we applied qpAdm and *DATES* to infer the ancestry proportions and timing of admixture in these groups.

Previous studies had shown that the early Steppe pastoralists derive ancestry from Eastern hunter-gatherers (EHG) and Caucuses hunter-gatherers (CHG) (25, 26). The CHG ancestry is maximized in Iranian Neolithic Farmers. Using data from 8 early Steppe pastoralists groups (seven Yamnaya-related and one Afanasievo group from Russia), we tested if each group could be modeled as a mixture of EHG_pooled and Iran_N_pooled reference populations (Supplementary Table SA). We observed that this provides a good fit for most Yamnaya samples except for individuals from the Baden culture of Hungary and Kalmykia from Russia (Supplementary Table S5.4).

To infer the timing of this admixture, we applied *DATES* to each of 6 early Steppe pastoralists groups using EHG_pooled and Iran_N_pooled reference populations. We observed significant dates ($Z > 2$) in 5 groups with similar admixture across all groups (within two standard errors) (Supplementary Table S6.1). The Yamnaya and Afanasievo cultures were genetically and culturally very similar and have been suggested to have very recent common ancestry (27). We confirmed this by estimating the genetic distances between the groups using *smartpca* (inbreed: YES) and found the $F_{ST}$ across groups is very low (~0.00-0.006) (Supplementary Table S6.2). Thus to infer more precise admixture dates, we pooled all the Yamnaya samples which show a good fit using *qpAdm* and had similar ancestry profiles with Afanasievo individuals and obtained a date of ~4,100 BCE (~4,000-4,300 BCE).

**Supplementary Table S6.1: *DATES* admixture times for EBA Steppe pastoralists groups.** *DATES* results for Yamnaya and Afanasievo groups using EHG_pooled and Iran_N_pooled as sources.

| **Sample** | ***DATES*** **mean ± SE (generations)** | **Average age across samples (BCE)** | **Admixture time (BCE)** |
|---|---|---|---|
| Russia_Afanasievo | 46 ± 4 | 2847 | 4135 ± 112 |
| Russia_Samara_EBA_Yamnaya | 41 ± 8 | 2934 | 4082 ± 224 |
| Kazakhstan_EBA_Yamnaya.SG | 52 ± 13 | 2940 | 4396 ± 364 |
| Russia_Caucasus_EBA_Yamnaya | 38 ± 14 | 2941 | 4005 ± 392 |
| Ukraine_EBA_Yamnaya | 61 ± 12 | 2900 | 4608 ± 336 |
| Pooled Early Steppe Pastoralists[#] (Yamnaya + Afanasievo) | 46 ± 3 | 2881 | 4169 ± 84 |

Note:
### Pooled Early Steppe Pastoralists include the following samples: Kazakhstan_EBA_Yamnaya.SG, Russia_Caucasus_EBA_Yamnaya, Russia_Samara_EBA_Yamnaya, Ukraine_EBA_Yamnaya, Ukraine_EBA_Yamnaya_published, Russia_Samara_EBA_Yamnaya_published2, Russia_Afanasievo

**Supplementary Table S6.2: Genetic distance ($F_{ST}$) in early Steppe pastoralists groups.** We performed smartpca using the option inbreed: YES to estimate $F_{ST}$ across groups.

| | Pop1 | Pop2 | Pop3 | Pop4 | Pop5 | Pop6 |
|---|---|---|---|---|---|---|
| Pop1 | 0.000 | 0.000 | 0.004 | 0.006 | 0.000 | 0.000 |
| Pop2 | 0.000 | 0.000 | 0.000 | 0.000 | 0.000 | 0.000 |
| Pop3 | 0.004 | 0.000 | 0.000 | 0.003 | -0.005 | 0.000 |
| Pop4 | 0.006 | 0.000 | 0.003 | 0.000 | 0.003 | 0.000 |
| Pop5 | 0.000 | 0.000 | -0.005 | 0.003 | 0.000 | 0.000 |
| Pop6 | 0.000 | 0.000 | 0.000 | 0.000 | 0.000 | 0.000 |

Note:
Pop1=Russia_Afanasievo; Pop2=Kazakhstan_EBA_Yamnaya.SG; Pop3=Russia_Caucasus_EBA_Yamnaya; Pop4=Russia_Samara_EBA_Yamnaya; Pop5=Ukraine_EBA_Yamnaya; Pop6=Ukraine_Ozera_EBA_Yamnaya

**Figure SA. Timing of WHG and EHG admixture in Iron Gates HG samples.** The time of admixture in Iron Gates HG samples. grouped in bins of C14 age of 500 years. The C14 age is shown on X-axis and the admixture time in BCE for corresponding samples is shown on the Y-axis.

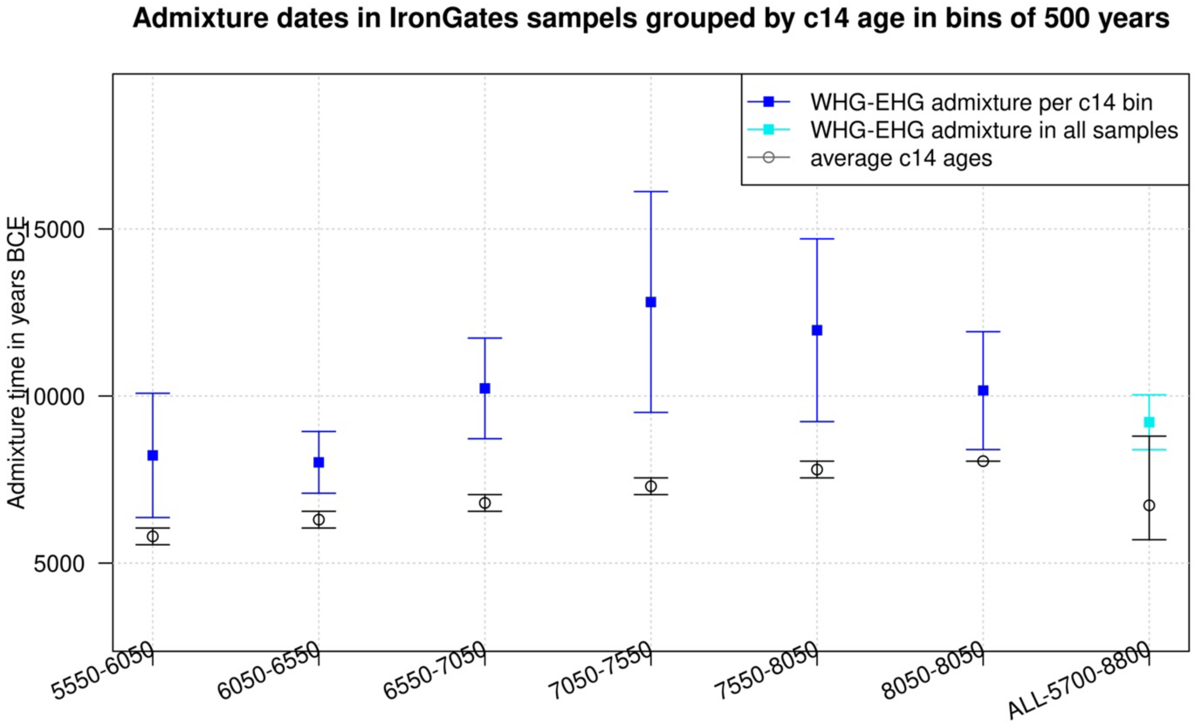


**Figure SB. *DATES* ancestry covariance decay curves.** We show the weighted ancestry covariance decay curves generated using *DATES* for all the target groups analyzed in the study. Each subplot shows the decay curve for one target population with the associated reference groups shown in the title. We plot the weighted covariance with genetic distance and obtained a date by fitting an exponential function with an affine term $y = Ae^{-\lambda d} + c$, where $d$ is the genetic distance in Morgans and $\lambda$ = ($t$+1) is the number of generations since admixture ($t$). We start the fit at genetic distance ($d$) > 0.5cM to minimize confounding with background LD and estimate a standard error by performing a weighted block jackknife removing one chromosome in each run. For each target, in the legend, we show the inferred average dates of admixture (± 1 SE) in generations before the individual lived, in BCE that accounts for the average age of all the individuals in the target and the mean generation time of human populations (see Methods). We also show the NRMSD values for all fitted curves and the plots with NRMSD > 0.7 are shown in grey. The colors match the population events described in main text and figures: formation of the Mesolithic hunter-gatherers (blue), formation of early Anatolian farmers (red), Anatolian farmer gene flow in Neolithic Europeans (orange), formation of early Steppe pastoralists (dark pink), Steppe Pastoralist-related gene flow in Steppe MLBA groups (light pink) and Steppe Pastoralist-related gene flow in Bronze age Europeans (green).

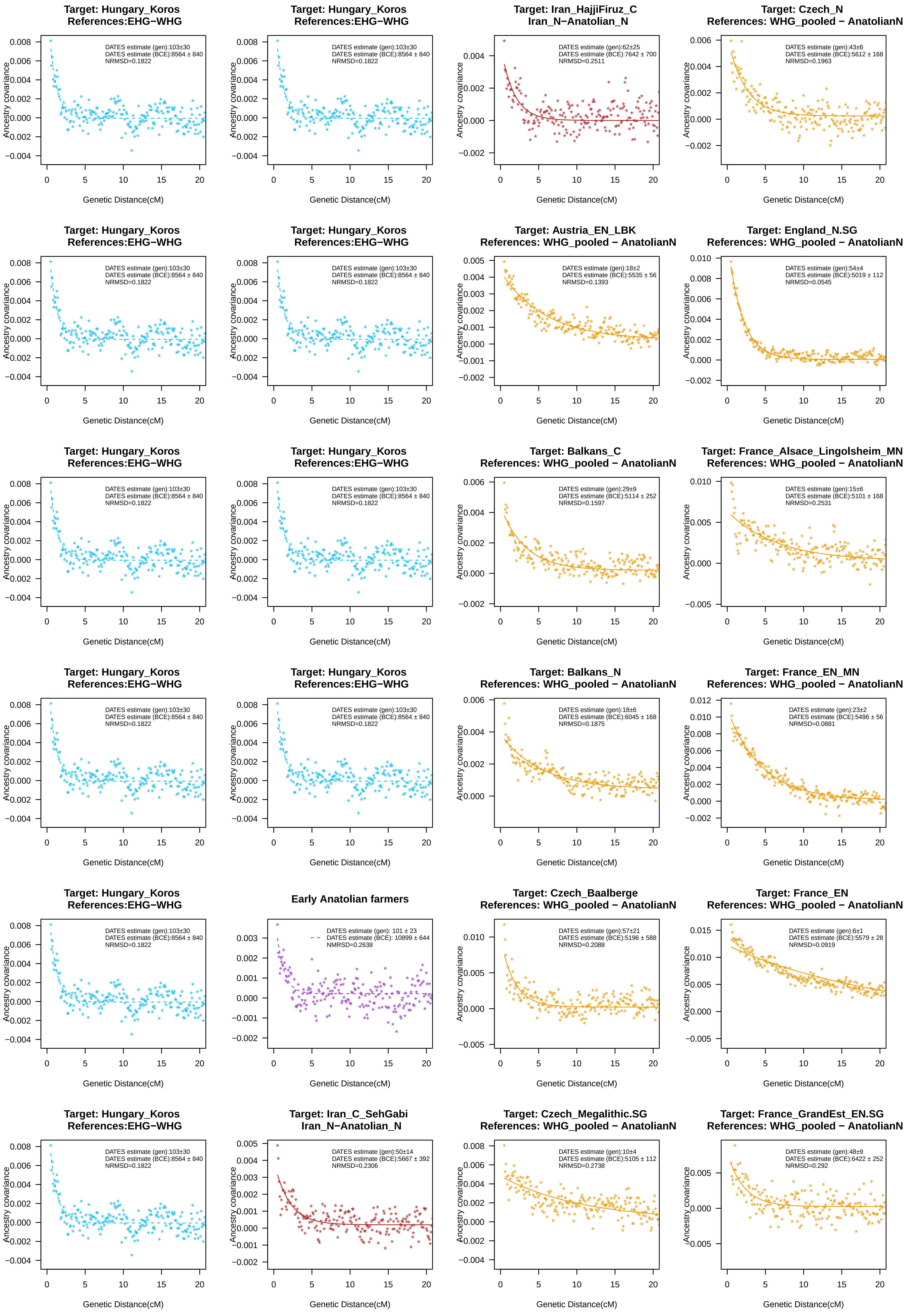
Target: Hungary_Koros
References:EHG−WHG
DATES estimate (gen):103±30
DATES estimate (BCE):8564 ± 840
NRMSD=0.1822
Target: Iran_HajjiFiruz_C
Iran_N−Anatolian_N
DATES estimate (gen):62±25
DATES estimate (BCE):7642 ± 700
NRMSD=0.2511
Target: Czech_N
References: WHG_pooled − AnatolianN
DATES estimate (gen):43±6
DATES estimate (BCE):5612 ± 168
NRMSD=0.1963
Target: Austria_EN_LBK
References: WHG_pooled − AnatolianN
DATES estimate (gen):18±2
DATES estimate (BCE):5535 ± 56
NRMSD=0.1393
Target: England_N.SG
References: WHG_pooled − AnatolianN
DATES estimate (gen):54±4
DATES estimate (BCE):5019 ± 112
NRMSD=0.0545
Target: Balkans_C
References: WHG_pooled − AnatolianN
DATES estimate (gen):29±9
DATES estimate (BCE):5114 ± 252
NRMSD=0.1597
Target: France_Alsace_Lingolsheim_MN
References: WHG_pooled − AnatolianN
DATES estimate (gen):15±6
DATES estimate (BCE):5101 ± 168
NRMSD=0.2531
Target: Balkans_N
References: WHG_pooled − AnatolianN
DATES estimate (gen):18±6
DATES estimate (BCE):6045 ± 168
NRMSD=0.1875
Target: France_EN_MN
References: WHG_pooled − AnatolianN
DATES estimate (gen):23±2
DATES estimate (BCE):5496 ± 56
NRMSD=0.0881
Early Anatolian farmers
DATES estimate (gen): 101 ± 23
DATES estimate (BCE): 10899 ± 644
NMRSD=0.2638
Target: Czech_Baalberge
References: WHG_pooled − AnatolianN
DATES estimate (gen):57±21
DATES estimate (BCE):5196 ± 588
NRMSD=0.2088
Target: France_EN
References: WHG_pooled − AnatolianN
DATES estimate (gen):6±1
DATES estimate (BCE):5579 ± 28
NRMSD=0.0919
Target: Iran_C_SehGabi
Iran_N−Anatolian_N
DATES estimate (gen):50±14
DATES estimate (BCE):5667 ± 392
NRMSD=0.2306
Target: Czech_Megalithic.SG
References: WHG_pooled − AnatolianN
DATES estimate (gen):10±4
DATES estimate (BCE):5105 ± 112
NRMSD=0.2738
Target: France_GrandEst_EN.SG
References: WHG_pooled − AnatolianN
DATES estimate (gen):48±9
DATES estimate (BCE):6422 ± 252
NRMSD=0.292
Ancestry covariance
Genetic Distance(cM)

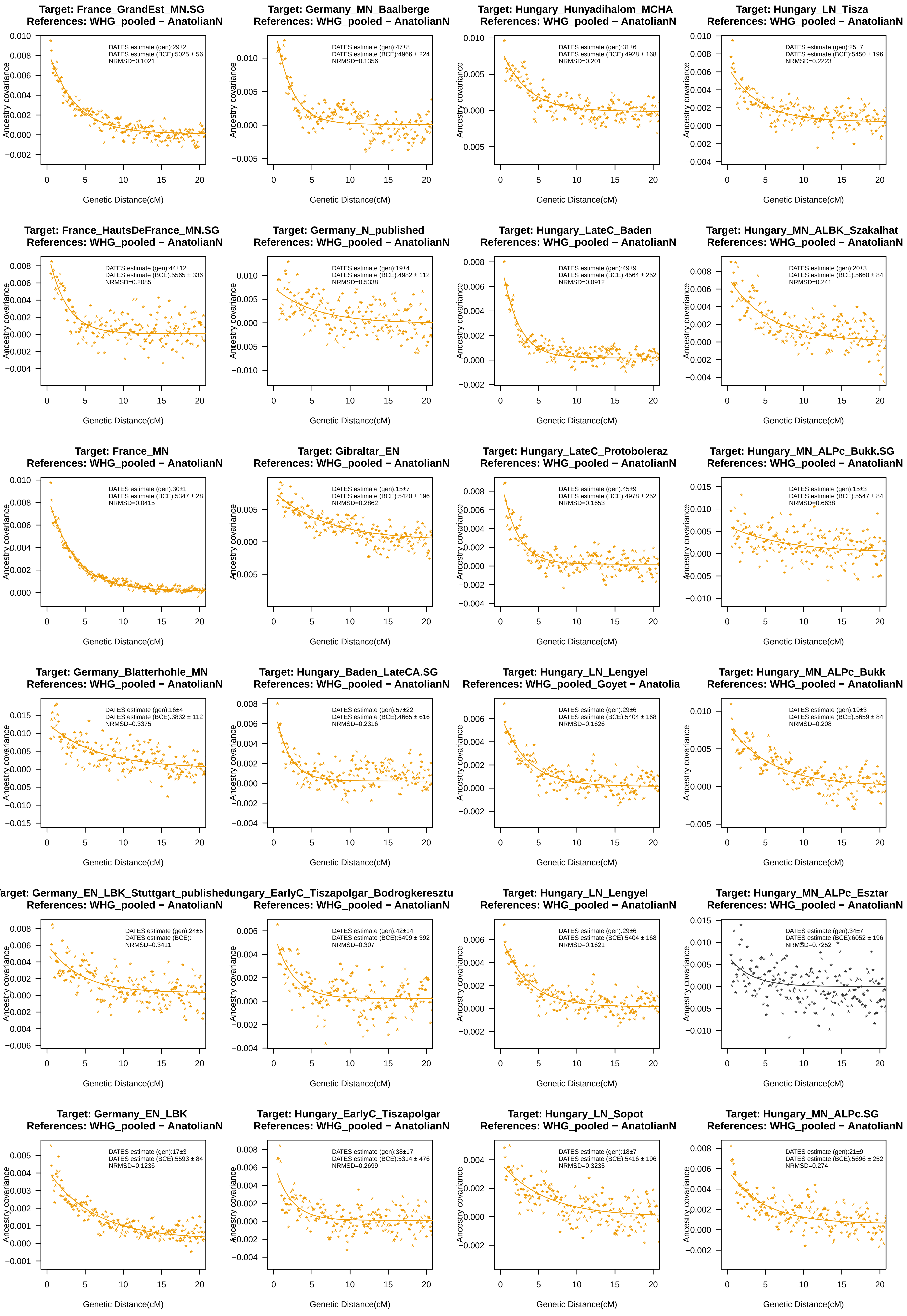
Target: France_GrandEst_MN.SG
References: WHG_pooled − AnatolianN
DATES estimate (gen):29±2
DATES estimate (BCE):5025 ± 56
NRMSD=0.1021
Target: Germany_MN_Baalberge
References: WHG_pooled − AnatolianN
DATES estimate (gen):47±8
DATES estimate (BCE):4966 ± 224
NRMSD=0.1356
Target: Hungary_Hunyadihalom_MCHA
References: WHG_pooled − AnatolianN
DATES estimate (gen):31±6
DATES estimate (BCE):4928 ± 168
NRMSD=0.201
Target: Hungary_LN_Tisza
References: WHG_pooled − AnatolianN
DATES estimate (gen):25±7
DATES estimate (BCE):5450 ± 196
NRMSD=0.2223
Target: France_HautsDeFrance_MN.SG
References: WHG_pooled − AnatolianN
DATES estimate (gen):44±12
DATES estimate (BCE):5565 ± 336
NRMSD=0.2085
Target: Germany_N_published
References: WHG_pooled − AnatolianN
DATES estimate (gen):19±4
DATES estimate (BCE):4982 ± 112
NRMSD=0.5338
Target: Hungary_LateC_Baden
References: WHG_pooled − AnatolianN
DATES estimate (gen):49±9
DATES estimate (BCE):4564 ± 252
NRMSD=0.0912
Target: Hungary_MN_ALBK_Szakalhat
References: WHG_pooled − AnatolianN
DATES estimate (gen):20±3
DATES estimate (BCE):5660 ± 84
NRMSD=0.241
Target: France_MN
References: WHG_pooled − AnatolianN
DATES estimate (gen):30±1
DATES estimate (BCE):5347 ± 28
NRMSD=0.0415
Target: Gibraltar_EN
References: WHG_pooled − AnatolianN
DATES estimate (gen):15±7
DATES estimate (BCE):5420 ± 196
NRMSD=0.2862
Target: Hungary_LateC_Protoboleraz
References: WHG_pooled − AnatolianN
DATES estimate (gen):45±9
DATES estimate (BCE):4978 ± 252
NRMSD=0.1653
Target: Hungary_MN_ALPc_Bukk.SG
References: WHG_pooled − AnatolianN
DATES estimate (gen):15±3
DATES estimate (BCE):5547 ± 84
NRMSD=0.6638
Target: Germany_Blatterhohle_MN
References: WHG_pooled − AnatolianN
DATES estimate (gen):16±4
DATES estimate (BCE):3832 ± 112
NRMSD=0.3375
Target: Hungary_Baden_LateCA.SG
References: WHG_pooled − AnatolianN
DATES estimate (gen):57±22
DATES estimate (BCE):4665 ± 616
NRMSD=0.2316
Target: Hungary_LN_Lengyel
References: WHG_pooled_Goyet − Anatolia
DATES estimate (gen):29±6
DATES estimate (BCE):5404 ± 168
NRMSD=0.1626
Target: Hungary_MN_ALPc_Bukk
References: WHG_pooled − AnatolianN
DATES estimate (gen):19±3
DATES estimate (BCE):5659 ± 84
NRMSD=0.208
arget: Germany_EN_LBK_Stuttgart_publishe
References: WHG_pooled − AnatolianN
DATES estimate (gen):24±5
DATES estimate (BCE):
NRMSD=0.3411
lungary_EarlyC_Tiszapolgar_Bodrogkereszt
References: WHG_pooled − AnatolianN
DATES estimate (gen):42±14
DATES estimate (BCE):5499 ± 392
NRMSD=0.307
Target: Hungary_LN_Lengyel
References: WHG_pooled − AnatolianN
DATES estimate (gen):29±6
DATES estimate (BCE):5404 ± 168
NRMSD=0.1621
Target: Hungary_MN_ALPc_Esztar
References: WHG_pooled − AnatolianN
DATES estimate (gen):34±7
DATES estimate (BCE):6052 ± 196
NRMSD=0.7252
Target: Germany_EN_LBK
References: WHG_pooled − AnatolianN
DATES estimate (gen):17±3
DATES estimate (BCE):5593 ± 84
NRMSD=0.1236
Target: Hungary_EarlyC_Tiszapolgar
References: WHG_pooled − AnatolianN
DATES estimate (gen):38±17
DATES estimate (BCE):5314 ± 476
NRMSD=0.2699
Target: Hungary_LN_Sopot
References: WHG_pooled − AnatolianN
DATES estimate (gen):18±7
DATES estimate (BCE):5416 ± 196
NRMSD=0.3235
Target: Hungary_MN_ALPc.SG
References: WHG_pooled − AnatolianN
DATES estimate (gen):21±9
DATES estimate (BCE):5696 ± 252
NRMSD=0.274
Ancestry covariance
Genetic Distance(cM)

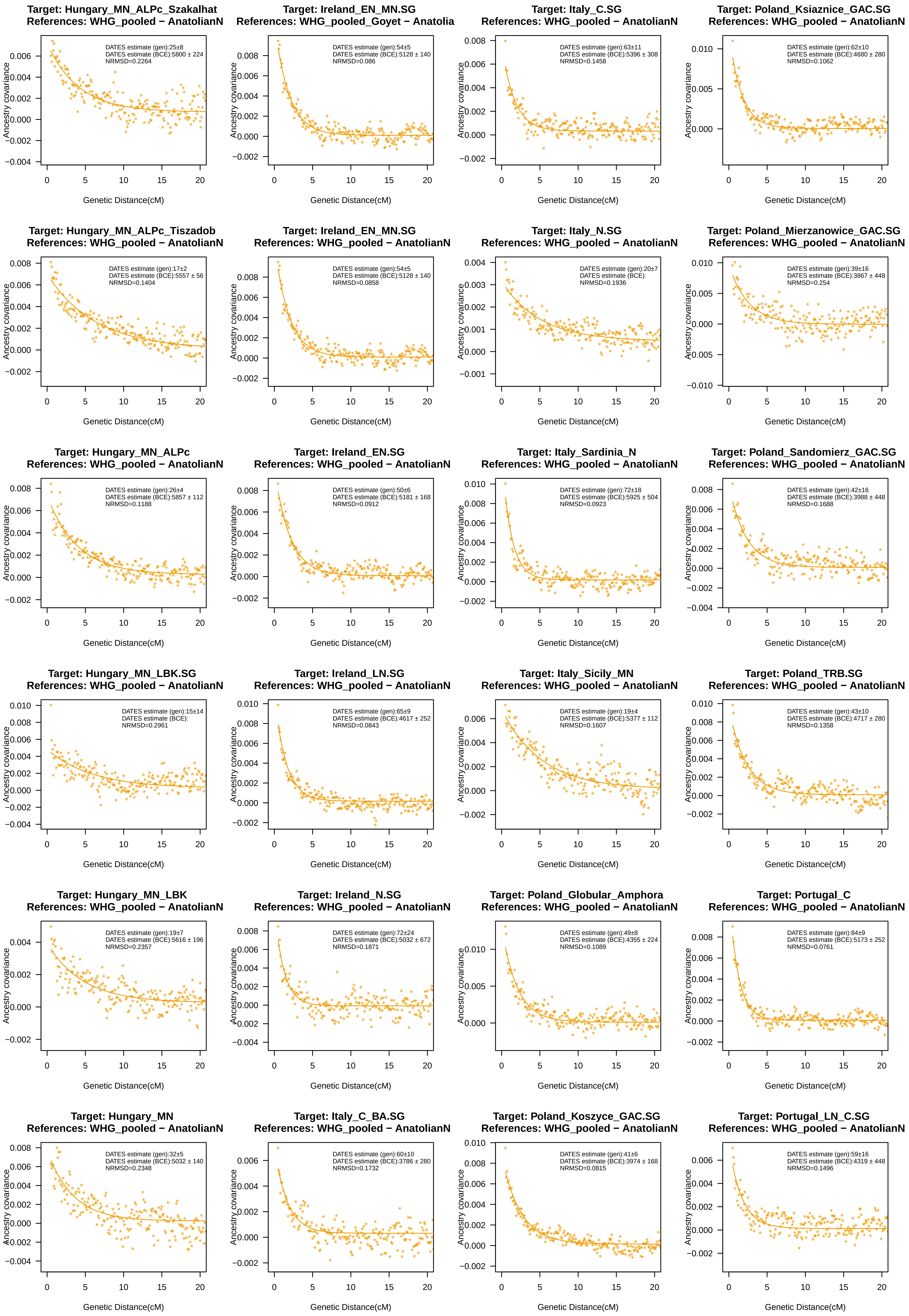
Target: Hungary_MN_ALPc_Szakalhat
References: WHG_pooled − AnatolianN
DATES estimate (gen):25±8
DATES estimate (BCE):5800 ± 224
NRMSD=0.2264
Target: Ireland_EN_MN.SG
References: WHG_pooled_Goyet − Anatolia
DATES estimate (gen):54±5
DATES estimate (BCE):5128 ± 140
NRMSD=0.086
Target: Italy_C.SG
References: WHG_pooled − AnatolianN
DATES estimate (gen):63±11
DATES estimate (BCE):5396 ± 308
NRMSD=0.1458
Target: Poland_Ksiaznice_GAC.SG
References: WHG_pooled − AnatolianN
DATES estimate (gen):62±10
DATES estimate (BCE):4680 ± 280
NRMSD=0.1062
Target: Hungary_MN_ALPc_Tiszadob
References: WHG_pooled − AnatolianN
DATES estimate (gen):17±2
DATES estimate (BCE):5557 ± 56
NRMSD=0.1404
Target: Ireland_EN_MN.SG
References: WHG_pooled − AnatolianN
DATES estimate (gen):54±5
DATES estimate (BCE):5128 ± 140
NRMSD=0.0858
Target: Italy_N.SG
References: WHG_pooled − AnatolianN
DATES estimate (gen):20±7
DATES estimate (BCE):
NRMSD=0.1936
Target: Poland_Mierzanowice_GAC.SG
References: WHG_pooled − AnatolianN
DATES estimate (gen):39±16
DATES estimate (BCE):3867 ± 448
NRMSD=0.254
Target: Hungary_MN_ALPc
References: WHG_pooled − AnatolianN
DATES estimate (gen):26±4
DATES estimate (BCE):5857 ± 112
NRMSD=0.1188
Target: Ireland_EN.SG
References: WHG_pooled − AnatolianN
DATES estimate (gen):50±6
DATES estimate (BCE):5181 ± 168
NRMSD=0.0912
Target: Italy_Sardinia_N
References: WHG_pooled − AnatolianN
DATES estimate (gen):72±18
DATES estimate (BCE):5925 ± 504
NRMSD=0.0923
Target: Poland_Sandomierz_GAC.SG
References: WHG_pooled − AnatolianN
DATES estimate (gen):42±16
DATES estimate (BCE):3988 ± 448
NRMSD=0.1688
Target: Hungary_MN_LBK.SG
References: WHG_pooled − AnatolianN
DATES estimate (gen):15±14
DATES estimate (BCE):
NRMSD=0.2961
Target: Ireland_LN.SG
References: WHG_pooled − AnatolianN
DATES estimate (gen):65±9
DATES estimate (BCE):4617 ± 252
NRMSD=0.0843
Target: Italy_Sicily_MN
References: WHG_pooled − AnatolianN
DATES estimate (gen):19±4
DATES estimate (BCE):5377 ± 112
NRMSD=0.1607
Target: Poland_TRB.SG
References: WHG_pooled − AnatolianN
DATES estimate (gen):43±10
DATES estimate (BCE):4717 ± 280
NRMSD=0.1358
Target: Hungary_MN_LBK
References: WHG_pooled − AnatolianN
DATES estimate (gen):19±7
DATES estimate (BCE):5616 ± 196
NRMSD=0.2357
Target: Ireland_N.SG
References: WHG_pooled − AnatolianN
DATES estimate (gen):72±24
DATES estimate (BCE):5032 ± 672
NRMSD=0.1871
Target: Poland_Globular_Amphora
References: WHG_pooled − AnatolianN
DATES estimate (gen):49±8
DATES estimate (BCE):4355 ± 224
NRMSD=0.1089
Target: Portugal_C
References: WHG_pooled − AnatolianN
DATES estimate (gen):84±9
DATES estimate (BCE):5173 ± 252
NRMSD=0.0761
Target: Hungary_MN
References: WHG_pooled − AnatolianN
DATES estimate (gen):32±5
DATES estimate (BCE):5032 ± 140
NRMSD=0.2348
Target: Italy_C_BA.SG
References: WHG_pooled − AnatolianN
DATES estimate (gen):60±10
DATES estimate (BCE):3786 ± 280
NRMSD=0.1732
Target: Poland_Koszyce_GAC.SG
References: WHG_pooled − AnatolianN
DATES estimate (gen):41±6
DATES estimate (BCE):3974 ± 168
NRMSD=0.0815
Target: Portugal_LN_C.SG
References: WHG_pooled − AnatolianN
DATES estimate (gen):59±16
DATES estimate (BCE):4319 ± 448
NRMSD=0.1496
Ancestry covariance
Genetic Distance(cM)

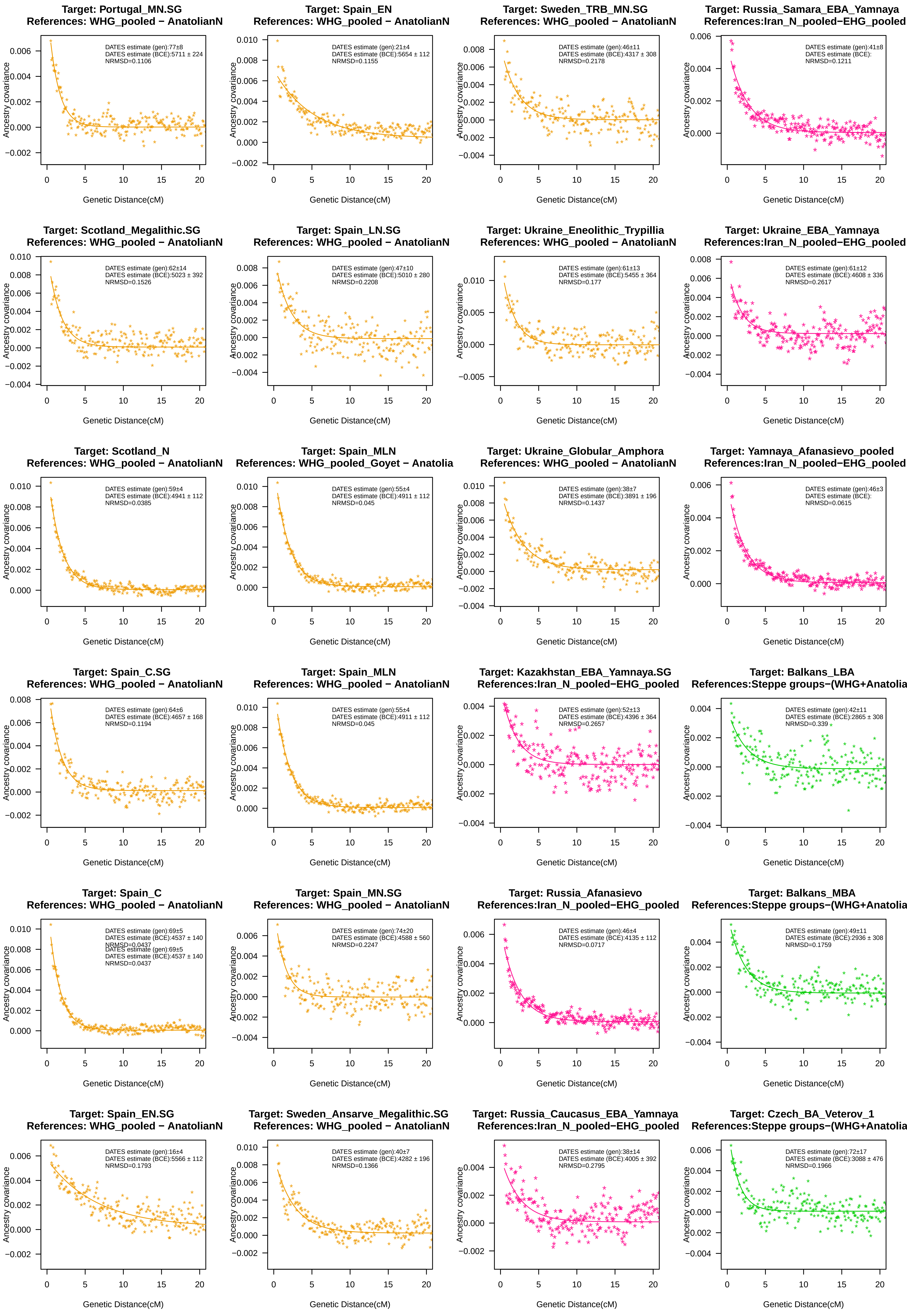

Target: Portugal_MN.SG
References: WHG_pooled − AnatolianN
DATES estimate (gen):77±8
DATES estimate (BCE):5711 ± 224
NRMSD=0.1106
Target: Spain_EN
References: WHG_pooled − AnatolianN
DATES estimate (gen):21±4
DATES estimate (BCE):5654 ± 112
NRMSD=0.1155
Target: Sweden_TRB_MN.SG
References: WHG_pooled − AnatolianN
DATES estimate (gen):46±11
DATES estimate (BCE):4317 ± 308
NRMSD=0.2178
Target: Russia_Samara_EBA_Yamnaya
References:Iran_N_pooled−EHG_pooled
DATES estimate (gen):41±8
DATES estimate (BCE):
NRMSD=0.1211
Target: Scotland_Megalithic.SG
References: WHG_pooled − AnatolianN
DATES estimate (gen):62±14
DATES estimate (BCE):5023 ± 392
NRMSD=0.1526
Target: Spain_LN.SG
References: WHG_pooled − AnatolianN
DATES estimate (gen):47±10
DATES estimate (BCE):5010 ± 280
NRMSD=0.2208
Target: Ukraine_Eneolithic_Trypillia
References: WHG_pooled − AnatolianN
DATES estimate (gen):61±13
DATES estimate (BCE):5455 ± 364
NRMSD=0.177
Target: Ukraine_EBA_Yamnaya
References:Iran_N_pooled−EHG_pooled
DATES estimate (gen):61±12
DATES estimate (BCE):4608 ± 336
NRMSD=0.2617
Target: Scotland_N
References: WHG_pooled − AnatolianN
DATES estimate (gen):59±4
DATES estimate (BCE):4941 ± 112
NRMSD=0.0385
Target: Spain_MLN
References: WHG_pooled_Goyet − Anatolia
DATES estimate (gen):55±4
DATES estimate (BCE):4911 ± 112
NRMSD=0.045
Target: Ukraine_Globular_Amphora
References: WHG_pooled − AnatolianN
DATES estimate (gen):38±7
DATES estimate (BCE):3891 ± 196
NRMSD=0.1437
Target: Yamnaya_Afanasievo_pooled
References:Iran_N_pooled−EHG_pooled
DATES estimate (gen):46±3
DATES estimate (BCE):
NRMSD=0.0615
Target: Spain_C.SG
References: WHG_pooled − AnatolianN
DATES estimate (gen):64±6
DATES estimate (BCE):4657 ± 168
NRMSD=0.1194
Target: Spain_MLN
References: WHG_pooled − AnatolianN
DATES estimate (gen):55±4
DATES estimate (BCE):4911 ± 112
NRMSD=0.045
Target: Kazakhstan_EBA_Yamnaya.SG
References:Iran_N_pooled−EHG_pooled
DATES estimate (gen):52±13
DATES estimate (BCE):4396 ± 364
NRMSD=0.2657
Target: Balkans_LBA
References:Steppe groups−(WHG+Anatolia
DATES estimate (gen):42±11
DATES estimate (BCE):2865 ± 308
NRMSD=0.339
Target: Spain_C
References: WHG_pooled − AnatolianN
DATES estimate (gen):69±5
DATES estimate (BCE):4537 ± 140
NRMSD=0.0437
DATES estimate (gen):69±5
DATES estimate (BCE):4537 ± 140
NRMSD=0.0437
Target: Spain_MN.SG
References: WHG_pooled − AnatolianN
DATES estimate (gen):74±20
DATES estimate (BCE):4588 ± 560
NRMSD=0.2247
Target: Russia_Afanasievo
References:Iran_N_pooled−EHG_pooled
DATES estimate (gen):46±4
DATES estimate (BCE):4135 ± 112
NRMSD=0.0717
Target: Balkans_MBA
References:Steppe groups−(WHG+Anatolia
DATES estimate (gen):49±11
DATES estimate (BCE):2936 ± 308
NRMSD=0.1759
Target: Spain_EN.SG
References: WHG_pooled − AnatolianN
DATES estimate (gen):16±4
DATES estimate (BCE):5566 ± 112
NRMSD=0.1793
Target: Sweden_Ansarve_Megalithic.SG
References: WHG_pooled − AnatolianN
DATES estimate (gen):40±7
DATES estimate (BCE):4282 ± 196
NRMSD=0.1366
Target: Russia_Caucasus_EBA_Yamnaya
References:Iran_N_pooled−EHG_pooled
DATES estimate (gen):38±14
DATES estimate (BCE):4005 ± 392
NRMSD=0.2795
Target: Czech_BA_Veterov_1
References:Steppe groups−(WHG+Anatolia
DATES estimate (gen):72±17
DATES estimate (BCE):3088 ± 476
NRMSD=0.1966
Ancestry covariance
Genetic Distance(cM)

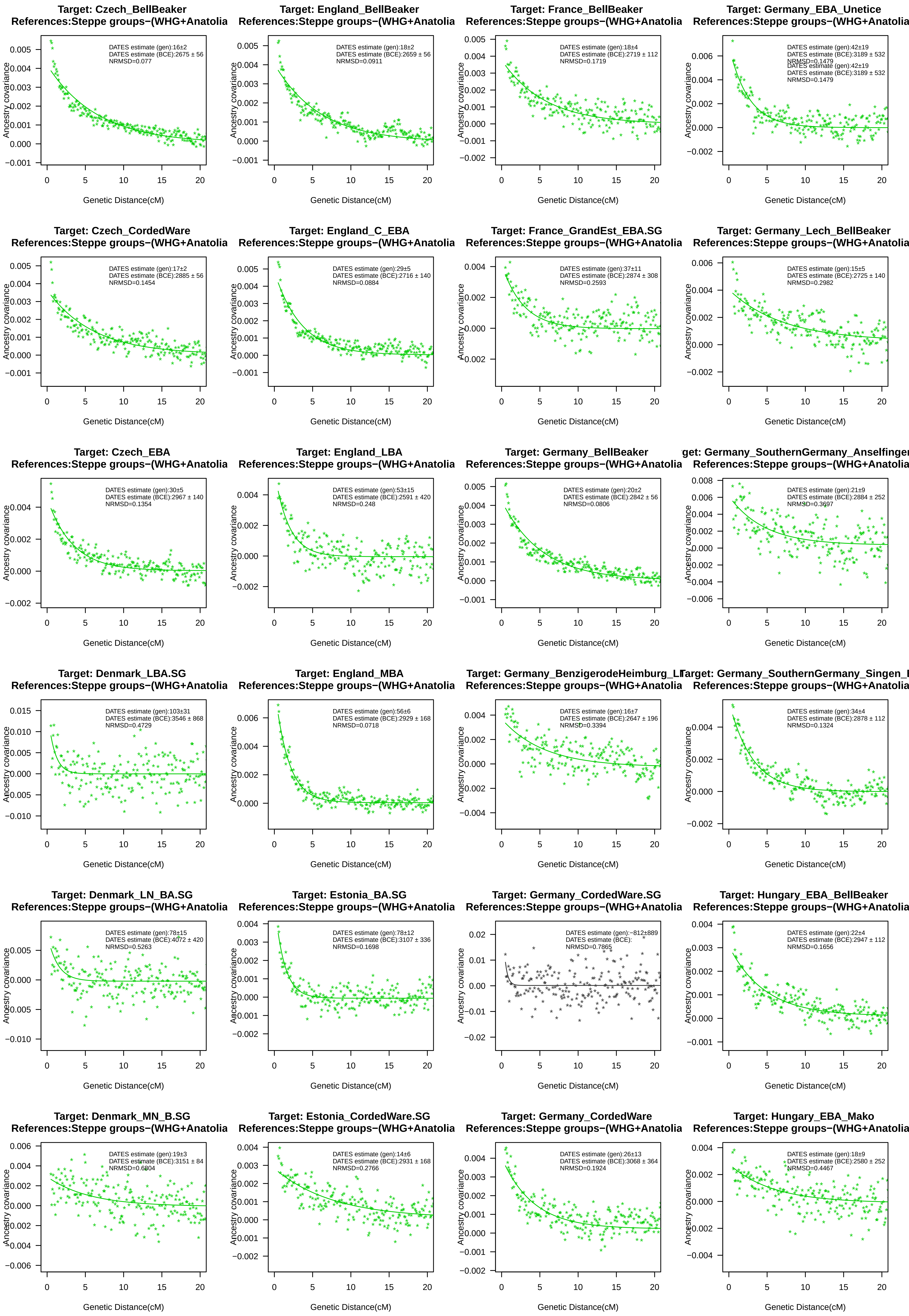
Target: Czech_BellBeaker
References:Steppe groups−(WHG+Anatolia
DATES estimate (gen):16±2
DATES estimate (BCE):2675 ± 56
NRMSD=0.077
Target: England_BellBeaker
References:Steppe groups−(WHG+Anatolia
DATES estimate (gen):18±2
DATES estimate (BCE):2659 ± 56
NRMSD=0.0911
Target: France_BellBeaker
References:Steppe groups−(WHG+Anatolia
DATES estimate (gen):18±4
DATES estimate (BCE):2719 ± 112
NRMSD=0.1719
Target: Germany_EBA_Unetice
References:Steppe groups−(WHG+Anatolia
DATES estimate (gen):42±19
DATES estimate (BCE):3189 ± 532
NRMSD=0.1479
Target: Czech_CordedWare
References:Steppe groups−(WHG+Anatolia
DATES estimate (gen):17±2
DATES estimate (BCE):2885 ± 56
NRMSD=0.1454
Target: England_C_EBA
References:Steppe groups−(WHG+Anatolia
DATES estimate (gen):29±5
DATES estimate (BCE):2716 ± 140
NRMSD=0.0884
Target: France_GrandEst_EBA.SG
References:Steppe groups−(WHG+Anatolia
DATES estimate (gen):37±11
DATES estimate (BCE):2874 ± 308
NRMSD=0.2593
Target: Germany_Lech_BellBeaker
References:Steppe groups−(WHG+Anatolia
DATES estimate (gen):15±5
DATES estimate (BCE):2725 ± 140
NRMSD=0.2982
Target: Czech_EBA
References:Steppe groups−(WHG+Anatolia
DATES estimate (gen):30±5
DATES estimate (BCE):2967 ± 140
NRMSD=0.1354
Target: England_LBA
References:Steppe groups−(WHG+Anatolia
DATES estimate (gen):53±15
DATES estimate (BCE):2591 ± 420
NRMSD=0.248
Target: Germany_BellBeaker
References:Steppe groups−(WHG+Anatolia
DATES estimate (gen):20±2
DATES estimate (BCE):2842 ± 56
NRMSD=0.0806
get: Germany_SouthernGermany_Anselfinger
References:Steppe groups−(WHG+Anatolia
DATES estimate (gen):21±9
DATES estimate (BCE):2884 ± 252
NRMSD=0.3697
Target: Denmark_LBA.SG
References:Steppe groups−(WHG+Anatolia
DATES estimate (gen):103±31
DATES estimate (BCE):3546 ± 868
NRMSD=0.4729
Target: England_MBA
References:Steppe groups−(WHG+Anatolia
DATES estimate (gen):56±6
DATES estimate (BCE):2929 ± 168
NRMSD=0.0718
Target: Germany_BenzigerodeHeimburg_LI
References:Steppe groups−(WHG+Anatolia
DATES estimate (gen):16±7
DATES estimate (BCE):2647 ± 196
NRMSD=0.3394
Target: Germany_SouthernGermany_Singen_E
References:Steppe groups−(WHG+Anatolia
DATES estimate (gen):34±4
DATES estimate (BCE):2878 ± 112
NRMSD=0.1324
Target: Denmark_LN_BA.SG
References:Steppe groups−(WHG+Anatolia
DATES estimate (gen):78±15
DATES estimate (BCE):4072 ± 420
NRMSD=0.5263
Target: Estonia_BA.SG
References:Steppe groups−(WHG+Anatolia
DATES estimate (gen):78±12
DATES estimate (BCE):3107 ± 336
NRMSD=0.1698
Target: Germany_CordedWare.SG
References:Steppe groups−(WHG+Anatolia
DATES estimate (gen):−812±889
DATES estimate (BCE):
NRMSD=0.7865
Target: Hungary_EBA_BellBeaker
References:Steppe groups−(WHG+Anatolia
DATES estimate (gen):22±4
DATES estimate (BCE):2947 ± 112
NRMSD=0.1656
Target: Denmark_MN_B.SG
References:Steppe groups−(WHG+Anatolia
DATES estimate (gen):19±3
DATES estimate (BCE):3151 ± 84
NRMSD=0.6804
Target: Estonia_CordedWare.SG
References:Steppe groups−(WHG+Anatolia
DATES estimate (gen):14±6
DATES estimate (BCE):2931 ± 168
NRMSD=0.2766
Target: Germany_CordedWare
References:Steppe groups−(WHG+Anatolia
DATES estimate (gen):26±13
DATES estimate (BCE):3068 ± 364
NRMSD=0.1924
Target: Hungary_EBA_Mako
References:Steppe groups−(WHG+Anatolia
DATES estimate (gen):18±9
DATES estimate (BCE):2580 ± 252
NRMSD=0.4467
Ancestry covariance
Genetic Distance(cM)

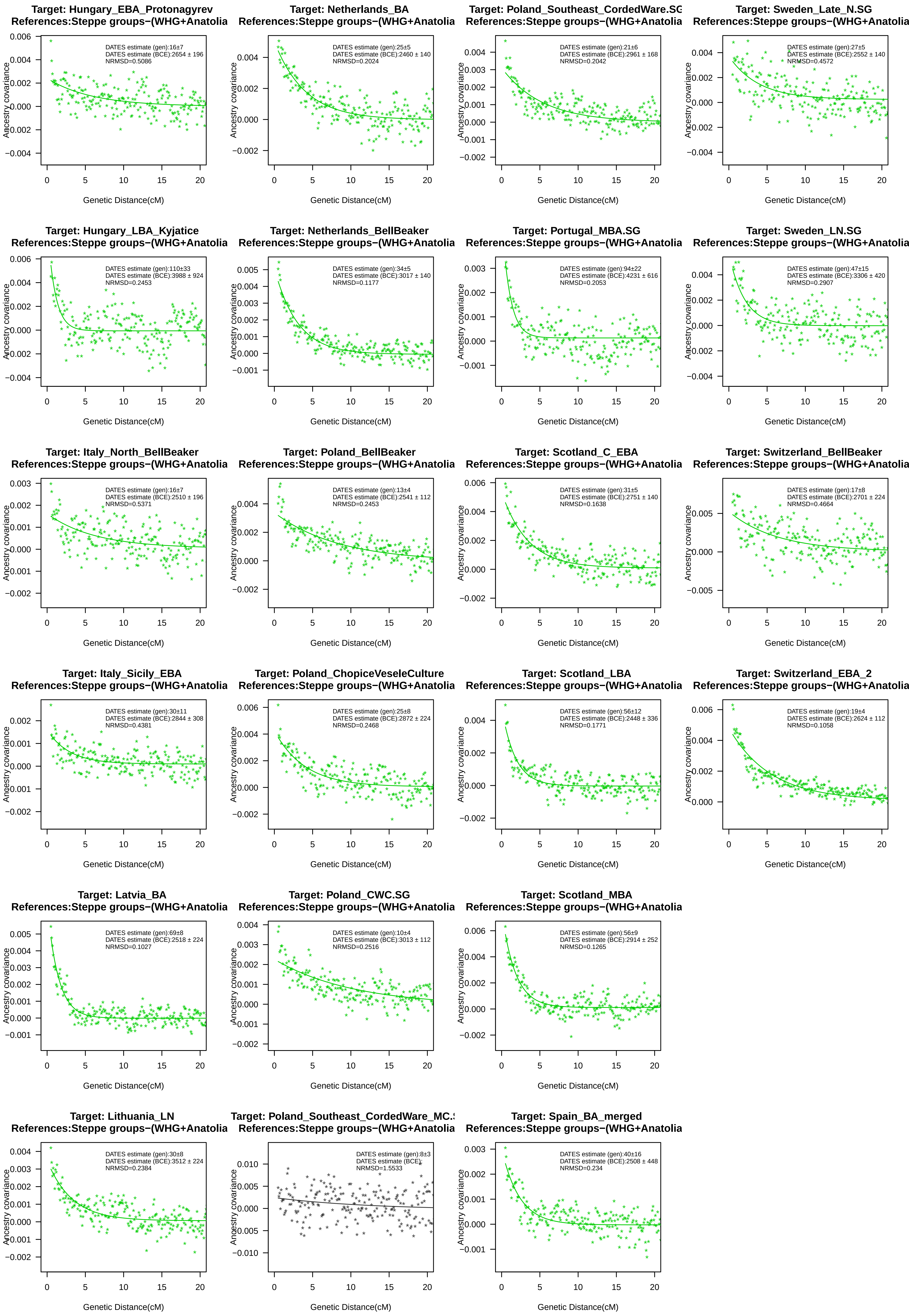

Target: Hungary_EBA_Protonagyrev
References:Steppe groups−(WHG+Anatolia
DATES estimate (gen):16±7
DATES estimate (BCE):2654 ± 196
NRMSD=0.5086
Target: Netherlands_BA
References:Steppe groups−(WHG+Anatolia
DATES estimate (gen):25±5
DATES estimate (BCE):2460 ± 140
NRMSD=0.2024
Target: Poland_Southeast_CordedWare.SG
References:Steppe groups−(WHG+Anatolia
DATES estimate (gen):21±6
DATES estimate (BCE):2961 ± 168
NRMSD=0.2042
Target: Sweden_Late_N.SG
References:Steppe groups−(WHG+Anatolia
DATES estimate (gen):27±5
DATES estimate (BCE):2552 ± 140
NRMSD=0.4572
Target: Hungary_LBA_Kyjatice
References:Steppe groups−(WHG+Anatolia
DATES estimate (gen):110±33
DATES estimate (BCE):3988 ± 924
NRMSD=0.2453
Target: Netherlands_BellBeaker
References:Steppe groups−(WHG+Anatolia
DATES estimate (gen):34±5
DATES estimate (BCE):3017 ± 140
NRMSD=0.1177
Target: Portugal_MBA.SG
References:Steppe groups−(WHG+Anatolia
DATES estimate (gen):94±22
DATES estimate (BCE):4231 ± 616
NRMSD=0.2053
Target: Sweden_LN.SG
References:Steppe groups−(WHG+Anatolia
DATES estimate (gen):47±15
DATES estimate (BCE):3306 ± 420
NRMSD=0.2907
Target: Italy_North_BellBeaker
References:Steppe groups−(WHG+Anatolia
DATES estimate (gen):16±7
DATES estimate (BCE):2510 ± 196
NRMSD=0.5371
Target: Poland_BellBeaker
References:Steppe groups−(WHG+Anatolia
DATES estimate (gen):13±4
DATES estimate (BCE):2541 ± 112
NRMSD=0.2453
Target: Scotland_C_EBA
References:Steppe groups−(WHG+Anatolia
DATES estimate (gen):31±5
DATES estimate (BCE):2751 ± 140
NRMSD=0.1638
Target: Switzerland_BellBeaker
References:Steppe groups−(WHG+Anatolia
DATES estimate (gen):17±8
DATES estimate (BCE):2701 ± 224
NRMSD=0.4664
Target: Italy_Sicily_EBA
References:Steppe groups−(WHG+Anatolia
DATES estimate (gen):30±11
DATES estimate (BCE):2844 ± 308
NRMSD=0.4381
Target: Poland_ChopiceVeseleCulture
References:Steppe groups−(WHG+Anatolia
DATES estimate (gen):25±8
DATES estimate (BCE):2872 ± 224
NRMSD=0.2468
Target: Scotland_LBA
References:Steppe groups−(WHG+Anatolia
DATES estimate (gen):56±12
DATES estimate (BCE):2448 ± 336
NRMSD=0.1771
Target: Switzerland_EBA_2
References:Steppe groups−(WHG+Anatolia
DATES estimate (gen):19±4
DATES estimate (BCE):2624 ± 112
NRMSD=0.1058
Target: Latvia_BA
References:Steppe groups−(WHG+Anatolia
DATES estimate (gen):69±8
DATES estimate (BCE):2518 ± 224
NRMSD=0.1027
Target: Poland_CWC.SG
References:Steppe groups−(WHG+Anatolia
DATES estimate (gen):10±4
DATES estimate (BCE):3013 ± 112
NRMSD=0.2516
Target: Scotland_MBA
References:Steppe groups−(WHG+Anatolia
DATES estimate (gen):56±9
DATES estimate (BCE):2914 ± 252
NRMSD=0.1265
Target: Lithuania_LN
References:Steppe groups−(WHG+Anatolia
DATES estimate (gen):30±8
DATES estimate (BCE):3512 ± 224
NRMSD=0.2384
Target: Poland_Southeast_CordedWare_MC.
References:Steppe groups−(WHG+Anatolia
DATES estimate (gen):8±3
DATES estimate (BCE):
NRMSD=1.5533
Target: Spain_BA_merged
References:Steppe groups−(WHG+Anatolia
DATES estimate (gen):40±16
DATES estimate (BCE):2508 ± 448
NRMSD=0.234
Ancestry covariance
Genetic Distance(cM)
